## Supplemental information for "The genomic and epigenomic evolutionary history of papillary renal cell carcinomas"

**Zhu et al.**

**Supplementary Method**

**Extraction of genomic DNA from fresh frozen tissues specimens**
After weight measurements, fresh frozen tissue samples (25 mg) were immediately put into 1 ml of 0.2 mg/ml Proteinase K (Qiagen) in DNA Lysis Buffer (10 mMTris-Cl (pH 8.0), 0.1 M EDTA (pH 8.0), and 0.5% (w/v) SDS) for 24 hrs at 56°C with shaking at 850 rpm in Thermomixer R (Eppendorf) until the tissue was completely lysed. Genomic DNA was extracted from fresh frozen tissue using the QIAmp DNA mini kit (Qiagen) according to the manufacturer's instructions. Each sample was eluted in volume of 200 µl AE buffer. DNA concentration was determined by Nanodrop spectrophotometer. All DNA samples were aliquoted and stored at −80 °C until use.

**DNA Processing**
DNA was quantified utilizing the QuantiFluor® dsDNA System (Promega Corporation, USA). DNA was normalized to 25ng/ul and underwent fragment analysis via AmpFLSTR™ Identifiler™ PCR Amplification Kit (ThermoFisher Scientific, USA). DNA samples are required to meet minimum mass and concentration thresholds for each assay, as well as show no evidence of contamination or profile discordance in the Identifiler assay. Samples meeting these requirements are aliquoted at the appropriate mass needed for downstream assay processing.

**Whole-genome sequencing**

Libraries were constructed and sequenced on the Illumina HiSeqX with the use of 151-bp paired-end reads for whole-genome sequencing.

**Preparation of libraries for cluster amplification and sequencing**
An aliquot of genomic DNA is taken from a stock sample at a target of 350ng in 50µL of solution to serve as the input into shearing. Samples undergo fragmentation by means of acoustic shearing using Covaris focused-ultrasonicator, targeting 385bp fragments. Following fragmentation, additional size selection is performed using a SPRI cleanup. Library preparation is performed using a commercially available kit provided by KAPA Biosystems (KAPA Hyper Prep without amplification module, product KK8505), and with palindromic forked adapters with unique 8 base index sequences embedded within the adapter (purchased from IDT). Following sample preparation, libraries were quantified using quantitative PCR (kit purchased from KAPA biosystems) with probes specific to the ends of the adapters. This assay was automated using Agilent's Bravo liquid handling platform. Based on qPCR quantification, libraries were normalized to 1.7nM. Samples are then pooled into 24-plexes and the pools are once again qPCRed. Samples were then combined with HiSeq X Cluster Amp Mix 1,2, and 3 into single wells on a strip tube using the Hamilton Starlet Liquid Handling System.

**Cluster amplification and sequencing**
Cluster amplification of the templates was performed according to the manufacturer's protocol (Illumina) using the Illumina cBot. Flowcells were sequenced on HiSeqX Sequencing-by-Synthesis Kits, then analyzed using RTA2.

**Filtering criteria of somatic mutation calling from whole genome sequencing data**
We used a revised method described by Jia-Jie Hao and colleagues to filter the somatic variants. Specifically, a variant was kept if at least 8 reads covered this variant in the normal samples and 3 reads in the tumor samples. Somatic variants with variant allele frequency (VAF) less than 0.07 were discarded. In addition, somatic variants were filtered using the VarScan[1] 'processSomatic' command with arguments tailored to our WGS samples with --min-tumor-freq 0.07, --max- normal-freq 0.02 and --p-value 0.05. The resulting somatic variants were further filtered to reduce false positives using the fpFilter Perl script (https://github.com/ckandoth/variant-filter). To remove possible germline variants from the called somatic variants, somatic variants were filtered against the dbSNP135 (https://www.ncbi.nlm.nih.gov/projects/SNP/snp_summary_byOrg.cgi?build_id=135), the 1000 genomes (phase 3 v5, http://www.internationalgenome.org/category/phase-3/), the ExAC v0.3.1 database (http://exac.broadinstitute.org/), and an in-house germline variant database from Italian population for SNPs with MAF<0.001. The filtered variants were annotated with Oncotator (version 1.1.9.0, https://github.com/broadinstitute/oncotator). To increase the sensitivity of somatic mutation calling, disease-associated variants annotated in the ClinVar database and the COSMIC database were retained. In addition, following the approach described by Stachler, et al. [2], we leveraged the multiple region sequencing and salvaged somatic variants that were detected from at least one tumor region and were missed in other tumor regions due to low VAF. In brief, Bam-readcount (https://github.com/genome/bam-readcount) was used to obtain read counts for

unique somatic variants across all tumor regions. A somatic variant was considered to be absent if either its VAF was less than 0.02 or there were fewer than three reads.

**Analysis of mutational signatures**

**Nucleosome Occupancy Analysis**

Nucleosomes are the basic units of DNA packaging, consisting of eight core histone proteins wrapping DNA sequence of about 147 bp long around itself. Consecutive nucleosomes are connected to each other by stretches of DNA called "linker DNA". To explore the relationship between mutational signatures and nucleosome occupancy, we downloaded the nucleosome occupancy signal of K562 cell line generated by micrococcal nuclease sequencing (MNase-seq) from ENCODE project [3].To examine the average nucleosome occupancy signal around the mutations, we considered all single point mutations with probability >= 0.5 to be in that signature. First, we took a window ± 1 kb centered at the mutation start position and counted all the nucleosome occupancy signals separately for each base in this window. We repeated this procedure for all mutations, accumulating and counting the signals within this 2 kb window. We then calculated the average nucleosome occupancy signal at each base by dividing the accumulated nucleosome occupancy signal by the accumulated number of counts for each base.
To interpret this nucleosome analysis, if there are no relationships between the nucleosome occupancy and the mutations in a signature, a flat line would be seen. If mutations occur at nucleosome positions we would see a peak where the mutations are centered. If the mutations occur at linker DNA stretches, there would be a trough (valley) where the mutations are centered.

**Replication Time Analysis**

The ENCODE project provides genome-wide assessment of DNA replication timing in various cell lines using sequencing-based "Repli-seq" methodology. We used MCF-7 cell line for all analysis. We downloaded wavelet-smoothed replication time signal data as well as replication peaks and replication valleys. Replication peaks, corresponding to replication initiation zones (Peaks) and replication termination zones (Valleys) were determined from local maxima and minima, respectively, in the wavelet-smoothed replication time signal data[3]. We sorted wavelet-smoothed replication time signal in descending order, divided them into ten deciles, each containing equal number of signals. Single point mutations with probability >= 0.5 for each signature are distributed into the corresponding decile in which it falls into, so that the first decile contains the mutations that are replicated the earliest and the last decile contains the mutations that are replicated the latest. The number of mutations in each decile is divided by the number of attributable bases (including 'A's, 'T's,'C's, 'G's and excluding 'N's) in the corresponding decile which gives the mutation density. Then, these mutation densities are divided by the highest mutation density which results in normalized mutation densities.

**Transcription and Replication Strand Bias Analysis**

Single point mutations are called on the + strand of the reference genome and converted into pyrimidine context. We identified the transcribed and un-transcribed strands of the genome based on hg19 NCBI RefSeq curated genes obtained from UCSC Table Browser using transcription start and end positions. Gene containing strand was annotated as un-transcribed strand whereas complementary strand was annotated as transcribed strand. We searched for overlapping single point mutations and gene transcripts. If there were at least one gene transcript on the same strand as the single point mutation, then 'un-transcribed strand' count was increased otherwise 'transcribed count' was increased. We considered all possible gene transcripts. Among 33,067 gene transcripts, 16,863 (1,639,967,346 bp) of them were on the "+" strand and 16,194 (1,570,613,951 bp) of them were on the "-" strand.
To investigate replication strand bias, we leveraged on replication time data peaks and valleys. We ordered the peaks and valleys consecutively and found the consecutive regions with consistent positive slope in terms of replication time signal between each consecutive valley and peak. In a similar manner we found the consecutive region with consistent negative slope between each peak and valley and annotated the leading and lagging strands, respectively. Note that each strand with lagging/leading strand annotations implies leading/lagging on the opposite strand. To annotate a DNA stretch, as leading or lagging strand, we discarded the latest replication termination zones ± 25,000 bp from the valley's midpoint, and we required at least 10,000 bp long DNA stretch with consistent positive or negative slope, respectively. All statistical tests for significance of strand-bias are based on Fisher exact test after FDR correction. Topography analysis of mutational signatures was performed using our previously developed methodology[4].

**Targeted Capture Sequencing**

**DNA Preparation:** For each sample, 50 ng genomic DNA was purified using Agencourt AMPure XP Reagent (Beckman Coulter Inc, Brea, CA, USA) according to manufacturer's protocol. An adapter-ligated library was prepared with the KAPA HyperPlus Kit (KAPA Biosystems, Wilmington, MA) using Bioo Scientific NEXTflex™ DNA Barcoded Adapters (Bioo Scientific, Austin, TX, USA) according to KAPA-provided protocol.

**Pre-Hybridization LM-PCR:** Genomic DNA sample libraries were amplified prior to hybridization by ligation-mediated PCR consisting of one reaction containing 20 µL library DNA, 25 µL 2x KAPA HiFi HotStart ReadyMix, and 5µL 10x Library Amplification Primer Mix (includes two primers whose sequences are: 5'-AATGATACGGCGACCACCGA-3' and 5'-CAAGCAGAAGACGGCATACGA-3'). PCR cycling conditions were as follows: 98˚C for 45 seconds, followed by 7 cycles of 98˚C for 15 s, 60˚C for 30 s, 72˚C for 30 s. The last step was an extension at 72˚C for 1 minute. The reaction was kept at 4˚C until further processing. The amplified material was cleaned with Agencourt AMPure XP Reagent (Beckman Coulter, Inc., Brea, CA, USA) according to the KAPA-provided protocol. Amplified sample libraries were quantified using Quant-iT™ PicoGreen dsDNA Reagent (Life Technologies, Carlsbad, CA, USA).

**Liquid Phase Sequence Capture:** Prior to hybridization, amplified sample libraries with unique barcoded adapters were combined in equal amounts into 1.1 µg pools for multiplex sequence capture. Sequence capture was performed with NimbleGen's SeqCap EZ Choice Library, using a custom design comprised of 254 candidate cancer driver genes (Roche NimbleGen, Inc., Madison, WI, USA). Prior to hybridization the following components were added to the 1.1 µg pooled sample library, 4 µL of NEXTflex HE Universal Oligo 1, 250 µM (5'-AATGATACGGCGACCACCGAGATCTACACTCTTTCCCTACACGACGCTCTTCCGATCT-3'), 40 µL total 25 µM NEXTflex INV-HE blocking oligos, equal volumes of each blocking oligo complementary to the barcodes in the pool (5'-CAAGCAGAAGACGGCATACGAGAT**X**GTGACT GGAGTTCAGACGTGTGCTCTTCCGATCT/C3 Spacer/-3', where **X** is 8-bases of sequence specific to adapter barcode used for library construction), and 5 µL of 1 mg/mL COT-1 DNA (Invitrogen, Inc., Carlsbad, CA, USA). Samples were dried down by puncturing a hole in the plate seal and processing in an Eppendorf 5301 Vacuum Concentrator (Eppendorf, Hauppauge, NY, USA) set to 60˚C for approximately 1 hour. To each dried pool, 7.5 µL of NimbleGen Hybridization Buffer and 3.0 µL of NimbleGen Hybridization Component A were added, and placed in a heating block for 10 minutes at 95˚C. The mixture was then transferred to 4.5 µL of EZ Choice Probe Library and hybridized at 47˚C for 64 to 72 hours. Washing and recovery of captured DNA were performed as described in NimbleGen SeqCap EZ Library SR Protocol.

**Post-Hybridization LM-PCR:** Pools of captured DNA were amplified by ligation-mediated PCR consisting of one reaction for each pool containing 20µl captured library DNA, 25 µL 2x KAPA HiFi HotStart ReadyMix, and 5µL 10x Library Amplification Primer Mix (includes two primers whose sequences are: 5'-AATGATACGGCGACCACCGA-3' and 5'-CAAGCAGAAGACGGCATACGA-3'). PCR cycling conditions were as follows: 98˚C for 45 seconds, followed by 12 cycles of 98˚C for 15 s, 60˚C for 30 s, 72˚C for 30 s. The last step was an extension at 72˚C for 1 minute. The reaction was kept at 4˚C until further processing. The amplified material was cleaned with Agencourt AMPure XP Reagent (Beckman Coulter, Inc., Brea, CA, USA) according to NimbleGen SeqCap EZ Library SR Protocol. Pools of amplified captured DNA were then quantified via Kapa's Library Quantification Kit for Illumina (Kapa Biosystems, Woburn, MA, USA) on the LightCycler 480 (Roche, Indianapolis, IN, USA).

**Sequencing:** The resulting post-capture enriched multiplexed sequencing libraries were used in cluster formation on an Illumina cBOT (Illumina, San Diego, CA, USA) and paired-end sequencing was performed using an Illumina HiSeq 4000 following Illumina-provided protocols for 2x150bp paired-end sequencing.

**Methylation analysis**

**DNA Preparation:** 400 ng of sample DNA, according to Quant-iT PicoGreen dsDNA quantitation (Life Technologies, Grand Island, NY), was treated with sodium bisulfite using the EZ-96 DNA Methylation MagPrep Kit (Zymo Research, Irvine, CA) according to manufacturer-provided protocol. Bisulfite conversion modifies non-methylated cytosines into uracil, leaving 5-methylcytosine (5mC) and 5-hydroxymethylcytosine (5hmC) unchanged. For every 95 samples, an internal control, NA07057 (Coriell Cell Repositories, Camden, NJ) was utilized to confirm the efficiency of bisulfite conversion and subsequent methylation analysis.

**Methylation Analysis:** High-throughput epigenome-wide methylation analysis, using Infinium MethylationEPIC BeadChip (Illumina Inc., San Diego, CA) which uses both Infinium I and II assay chemistry technologies, was performed according to manufacturer-provided protocol. Bisulfite-treated samples were denatured and neutralized then whole genome amplified, isothermally, to increase the amount of DNA template. The amplified product was enzymatically fragmented, precipitated and resuspended in hybridization buffer. Eight samples were applied to each BeadChip and hybridized overnight where fragmented DNA samples anneal to locus-specific 50mers (covalently linked to one of over 800,000 bead types). Two beadtypes correspond to each CpG locus for Infinium I assays: one bead type corresponds to methylated, another bead type to unmethylated state of the CpG site, while one beadtype corresponds to each CpG locus for Infinium II assays. Single-base extension of the oligos on the BeadChip, using the captured DNA as template, incorporates tagged nucleotides on the BeadChip, which are subsequently fluorophore labeled during staining. The Illumina iScan scanned the BeadChips at two wavelengths to create image and intensity files.

**Data Analysis:** The intensity files from the Illumina methylation assay on the MethylationEPIC platform were processed and analyzed with the R programming language using the R package "minfi". Briefly, raw intensity file (idats) are loaded into R using minfi. Samples are excluded if the percent of probes with detection p value greater than 0.01 is greater than 4%. Concordance is checked for both expected and unexpected replicates using the ~60 polymorphic SNPs on the array. Raw methylation beta values are normalized according to previously published methods (Fortin et al. 2014).

**Genotyping analysis**

**Illumina Infinium Genotyping Array BeadChips:** High-throughput, genome-wide SNP genotyping, using Infinium BeadChip technology (Illumina Inc. San Diego, CA), was performed at the Cancer Genomics Research Laboratory (CGR). Genotyping was performed per the manufacturer's guidelines using the Infinium automated protocol. For the Infinium HumanOmniExpress-24 chip, 20ng genomic DNA, quantitated using Quant-iT™ PicoGreen dsDNA Reagent (ThermoFisher Scientific, Waltham, MA, USA) is denatured and neutralized, then isothermally amplified by whole-genome amplification. The amplified product is enzymatically fragmented, then precipitated and re-suspended. Resuspended samples are denatured, then hybridized to locus-specific 50-mer oligonucleotides which are attached to 1-micron beads on the BeadChip. These 50-mer probes stop one base before the location of interest. Enzymatic single-base extension of the oligos on the BeadChip, using the captured DNA as a template, incorporates tagged nucleotides on the BeadChip, which are subsequently fluorophore-labeled during staining. The fluorescent label determines the genotype call for the sample. The Illumina iScan scans the BeadChips at two wavelengths to detect the fluorescent label, creating image files that are converted into genotype calls based on the detected fluorescence.

**Illumina GenomeStudio Genotyping Module v2.0:** The GenomeStudio is a Windows-based software that analyzes Illumnia genotyping data. Typically, a clustering project is created from a sample sheet to load sample intensities (*.idat files), and a SNP manifest (*.bpm file) for the type of array used. After scan, an initial clustering is performed to identify samples that need to be re-scanned from the laboratory. Once re-scan is complete, any sample that yields higher call rate by re-scan will have its intensity file substituted with a better one. Then the final clustering of all samples performed using a cluster position file (*.egt file). Once final clustering is finished, genotypes of all samples are exported through the 'Report Wizard'. Genotypes can be exported into 'Locus X DNA' format (LBD), then further converted into multiple different formats (eg. plink or glu format), or can be exported directly into PLINK format if a plug-in is installed for this function.

**Validation of SCNAs calling by FACETs**

Allele-specific copy-number alteration (SCNA) analysis was performed using FACETS5 v0.5.6 (https://github.com/mskcc/facets) with the following parameters to increase the strictness of the segments: normal read filter depth=15; window size = 5000. The 'snp-pileup' command with argument '--min-map-quality 20 --min-base-quality 20 --min-read-counts 15,0' was used to extract read counts of the reference and alternate alleles from tumor and normal tissue samples BAM files separately. SCNA events were allocated to different genotypes according to the total copy number and minor copy numbers. SCNA events were defined as chromosome-arm if the same SCNA level overlapped with at least 90% of the chromosome arm's coordinates. Focal SCNA events were defined if their individual lengths were less than half the length of a chromosome arm. The fraction of the copy-number-altered genome was defined as the fraction of the genome with either non-diploid copy-number or evidence of loss of heterozygosity. Besides SCNA events, tumor purity and ploidy were also estimated using FACETS.

**Validation of structural variants**

**Comparison between Meerkat and Novobreak SV calling**

We compared the SV results obtained by Meerkat with those obtained by Novobreak (Chong, et al., v1.1.3rc). “Filter_sv2.pl” was used to filter the SVs, and only SVs with calling quality “QUAL” above 30 were kept. We examined whether SVs called by Meerkat could be called by Novobreak and *vice versa*, by comparing both left and right breakpoints in a given window size from 0 to 10 Mbp. The results showed that Meerkat-based calling was more conservative than Novobreak regardless the window size; Novobreak detected almost twice the number of SVs detected by Meerkat. About 60% of SVs identified by Meerkat could be replicated by Novobreak, while 32% of SVs identified by Novobreak could be replicated by Meerkat. Thus, we decided to use the SVs identified by Meerkat.

**Gene fusion validation**

We selected four in-frame fusions: *MALAT-TFEB, MET-MET* deletion, *STRN-ALK*, and *EWSR1-PATZ1* for further validation. Briefly, total RNA from tumors with a fusion detected through our WGS pipeline were reverse transcribed using Superscript III (Thermo) primed with random hexamers. Subsequently, PCR using gene specific primers were performed with Platinum Taq supermix (Thermo) at 55C annealing temp. PCR products were agarose gel purified and sequenced on an ABI 3730xl. RNA samples to validate *EWSR-PATZ1* and *MALAT1-TFEB* fusions had relatively high RIN score (RIN=8 and 6, respectively). These fusions were confirmed by Sanger sequencing as valid fusion products. In contrast, RNA samples to validate the *MET-MET* deletion and *STRN-ALK* fusion were derived from FFPE samples with RIN=2.6. We were unable to validate these fusions by this method, potentially due to the poor RNA quality.

**Validation of additional structural variants**

381 structural variants, identified by Meerkat from whole genome sequencing data, were selected for validation. These events were found in pRCC1 and pRCC2 tumors with at least three samples. For each structural variant to be validated, 250bp of sequence was pulled from one side of each Meerkat breakpoint and spliced together, using the modified script “primers.pl” from the Meerkat package, such that the resulting 500bp Fasta file would serve as a reference for primer design of an amplicon that would only be generated in samples containing the structural variant. Alignment and orientation of the 500bp Fasta sequences were manually checked using UCSC BLAT. These 381 Fasta files were provided to the Ampliseq Designer v6.1.4 (ThermoFisher Scientific, Waltham, MA, USA) for custom Ampliseq panel design. 303 events had primer pairs successfully designed across the SV breakpoints and were included in a custom panel, with an average amplicon length of 352bp. A previously-designed custom panel, containing 74 amplicons, with primer pairs designed in hg19 rather than across custom SV breakpoints, was combined with this custom panel in order to ensure that samples containing only one or a few of the SV events would amplify sufficiently to generate quantifiable library. Sample DNA (30ng) was amplified using this custom AmpliSeq primer pool, and libraries were prepared following the manufacturer’s Ion AmpliSeq Library Preparation protocol. Individual samples were barcoded, pooled, templated, and sequenced on the Ion Torrent PGM Sequencer using the Ion Chef and sequenced on a 318 chip per manufacturer’s instructions. Sequence data was aligned to the custom reference used to generate the panel design, and reads were identified which spanned the SV breakpoint at 250bp.

**Comparison between SCNAs and SVs breakpoints**

We compared the breakpoints between all SCNAs (estimated by Battenberg) and SVs (estimated by Meerkat). We found that 64% of the SVs identified by Meerkat had no Battenberg breakpoint within 300KB in any sample (Supplementary Figure 17). Interestingly, the SVs that were identified as clonal by Battenberg (i.e., a breakpoint within 300KB of the SV was found by Battenberg in every sample) were dominated by inter-chromosomal translocations. In contrast, those SVs that were unobserved by Battenberg (i.e., no samples had a Battenberg breakpoint within 300KB of the SV) were dominated by tandem duplications.

These results suggest that Battenberg (and probably copy number callers in general) has poor sensitivity for calling certain types of SVs and shows the value of combined analysis of SVs and SCNAs. Notably, the median size of SVs was consistently smaller than the median SCNA size (in 73/74 samples; IQR of SVs = [36.9KB, 960.5KB], IQR of

SCNAs = [35.0MB, 89.6MB]), suggesting that many of the small SVs observed as subclonal were below the detection limit of copy number calling. In particular, 45% of the SVs that did not have a nearby copy number breakpoint were tandem duplications, whereas the SVs that were observed clonally in copy number calling were dominated by inter-chromosomal translocations (46% of the clonal variants, **Supplementary Figure 17**).

**Retrotranspon analysis**

The LINE-1 insertions (**Fig. 4a**) identified by TraFiC (Tubio et al.) were located in introns and intergenic regions. At least five insertions were judged to be *bona fide* L1HS insertions due to simultaneous presence of the following features: 1) concordant insertion position confirmed by sequence reads of different amplicons; 2) a 5' end corresponding to a portion of the L1HS consensus sequence; 3) TSD (target site duplications) of 11-17 bp and an A/T-rich insertion site (typical features of target-primed retrotransposition). The precise length of these insertions was not determined, because only the 5' and 3' ends of the inserted fragments were present in the amplicons generated in the NGS libraries. However, based on the alignment of the sequenced 5' ends with the L1HS consensus sequence (obtained from the public database www.girinst.org), no full-length L1 insertion was present. The minimal estimated length of the observed LINE-1 insertions ranged from few hundreds to about 1200 base pairs. At least three of the insertions could potentially affect the expression of proteins involved in chromatin regulation and chromosome structural maintenance and (in turn) the maintenance of genome integrity: 1) Chromosome 9: 1929235. From ENCODE analysis, this insertion appears to be located in a gene-regulatory site. *SMARCA2*, a member of the SWI/SNF family of chromatin remodeling factors, is located about 80Kb away. 2) Chromosome 7: 18991805-18992490. The insertion is located in an intron of *HDAC9* (Histone deacetylase 9), a gene whose increased or ectopic expression is involved in carcinogenesis[6]. 3) Chromosome 18: 34786106-34786799. Located in an intron of *KIAA1328* (hinderin), which binds to SMC3 (structural maintenance of chromosomes 3). SMC3 knockdown triggers genomic instability[7].

**Mutational signatures features.**

Examining the effect of genomic architecture on SNV mutational signatures revealed that mutations attributed to signature 1 were enriched in both early replicating regions of the genome and the parts of the genome with higher nucleosome occupancy (**Supplementary Figure 27**). In contrast, the mutations attributed to signatures 5 and 40 were mostly unaffected by replication timing and nucleosome occupancy. Interestingly, signatures 1, 5, and 40 exhibited a statistically significant transcriptional strand-bias. Statistically significant replication strand-bias was observed for signature 40. There were not sufficient numbers of mutations to evaluate the transcription and replication strand biases for signatures 2, 8, and 13.

*De novo* extraction of indel mutational signatures was performed across all samples using an indel classification scheme outlined in Alexandrov, et al. [8]. Three distinct indel signatures were identified and termed signatures INDEL-A, INDEL-B, and INDEL-C (**Supplementary Figure 28**). Comparison of these three *de novo* deciphered signatures to the global consensus set of mutational signatures revealed that signatures INDEL-A through INDEL-C are linear combinations of six previously known indel mutational signatures (**Supplementary Table 3**): signatures ID-1, ID-2, ID-3, ID-5, ID-6 and ID-8. Signatures INDEL-A and INDEL-B were found in most samples, while signature INDEL-C was found only in a subset of samples. The number of indels attributed to each indel mutational signature varied from a dozen to more than 800 (**Supplementary Data 16**) indicating that different indel mutational processes have been active in papillary renal cell carcinoma.

**Supplementary references:**

1. Koboldt, D.C. *et al.* VarScan 2: somatic mutation and copy number alteration discovery in cancer by exome sequencing. *Genome Res* **22**, 568-76 (2012).
2. Stachler, M.D. *et al.* Paired exome analysis of Barrett's esophagus and adenocarcinoma. *Nat Genet* **47**, 1047-55 (2015).
3. Consortium, E.P. An integrated encyclopedia of DNA elements in the human genome. *Nature* **489**, 57-74 (2012).
4. Morganella, S. *et al.* The topography of mutational processes in breast cancer genomes. *Nat Commun* **7**, 11383 (2016).

5. Shen, R. & Seshan, V.E. FACETS: allele-specific copy number and clonal heterogeneity analysis tool for high-throughput DNA sequencing. *Nucleic Acids Res* **44**, e131 (2016).
6. Lapierre, M. *et al.* Histone deacetylase 9 regulates breast cancer cell proliferation and the response to histone deacetylase inhibitors. *Oncotarget* **7**, 19693-708 (2016).
7. Ghiselli, G. SMC3 knockdown triggers genomic instability and p53-dependent apoptosis in human and zebrafish cells. *Mol Cancer* **5**, 52 (2006).
8. Alexandrov, L. *et al.* The Repertoire of Mutational Signatures in Human Cancer. *bioRxiv* (2018).
9. Alexandrov, L.B. *et al.* Clock-like mutational processes in human somatic cells. *Nat Genet* **47**, 1402-7 (2015).

**Supplementary Table 1.** Patient characteristics. They include age at diagnosis, gender, tumor size, stage and survival status.

| Subject | Age at diagnosis | Sex | Tumor size in cm | Clinical staging | Life status at the last follow-up | Survival in months |
|---|---|---|---|---|---|---|
| **pRCC2_1298_01** | 57.4 | M | 8.7 | 3 | ALIVE | 30 |
| **mixRCC_1336_01** | 55.3 | M | 2.8 | 1 | ALIVE | 30 |
| **pRCC1_1394_02** | 43.2 | M | 5 | 1 | ALIVE | 27 |
| **pRCC2_1396_01**[a] | 68.9 | M | 3 (Right) + 2 (Left) | 4 | ALIVE | 24 |
| **pRCC2_1410_02** | 43.1 | M | 5 | 1 | ALIVE | 24 |
| **pRCC1_1416_04** | 42.7 | F | 5 | 1 | ALIVE | 24 |
| **pRCC2_1429_03** | 75.3 | M | 9 | 3 | ALIVE | 22 |
| **pRCC1_1472_01** | 65.1 | M | 2.5 | 1 | ALIVE | 18 |
| **pRCC2_1479_03** | 77.3 | F | 5.2 | 1 | ALIVE | 18 |
| **pRCC2_1494_04** | 82.6 | M | 8 | 2 | ALIVE | 16 |
| **pRCC1_1501_01**[b] | 59.7 | M | 2 + 1.4 | 1 | ALIVE | 20 |
| **pRCC1_1546_01** | 40.4 | M | 3.8 | 1 | ALIVE | 12 |
| **pRCC1_1550_02** | 53.6 | M | 2.5 | 1 | ALIVE | 16 |
| **pRCC2_1552_03** | 49.6 | M | 7.4 | 2 | ALIVE | 9 |
| **pRCC2_1568_04**[b] | 68.1 | M | 1.7 + 2.4 + 7.4 | 4 | ALIVE | 2 |
| **pRCC1_1654_01** | 63.1 | M | 4.2 | 1 | ALIVE | 4 |
| **pRCC1_1662_01** | 63.2 | M | 2.5 | 1 | ALIVE | 6 |
| **pRCC1_1671_08** | 62.9 | M | 8 | 2 | ALIVE | 1 |
| **pRCC1_1689_06** | 46.3 | F | 7.2 | 2 | ALIVE | 6 |
| **rSRC_1697_10** | 58.5 | F | 7.2 | Unknown | DEAD | 7 |
| **pRCC1_1699_01** | 59.5 | F | 4.2 | 1 | ALIVE | 7 |
| **pRCC1_1708_01** | 59.4 | M | 4.5 | 1 | ALIVE | 2 |
| **pRCC1_1735_01** | 71.6 | M | 5 | 1 | ALIVE | 1 |
| **pRCC2_1782_08** | 53.2 | F | 4.8 | 3 | Unknown | Unknown |
| **pRCC2_1799_02** | 65.3 | F | 1.8 | 1 | ALIVE | 1 |
| **pRCC2_1824_13** | 51.8 | M | 10.5 | 4 | Unknown | Unknown |
| **pRCC2_1851_04** | 86.0 | F | 8.8 | 3 | Unknown | Unknown |
| **cdRCC_1929_03** | 76.0 | M | 5.7 | 3 | Unknwon | Unknown |
| **mixRCC_2028_03** | NA | M | 4.8 | 3 | ALIVE | 7 |

[a]: this subject had a bilateral cancer. The samples were taken from the larger lesion on the right side

[b]: this subject had multiple cancer lesions in the same kidney. The samples were taken from the largest lesion

**Supplementary Table 2.** Annotations of allele-specific copy number alteration types. Allele-specific copy number alteration calling based on total copy number and minor copy number. LOH: Loss of heterozygosity.

| Number of Segmentation | Total copy number | Minor copy number | Calling | Note |
|---|---|---|---|---|
| 49 | 0 | 0 | HOMD | homozygous deletion |
| 350 | 1 | 0 | DLOH | deletion LOH |
| 46 | 2 | 0 | NLOH | copy-neutral LOH |
| 1916 | 2 | 1 | HET | heterozygous |
| 34 | 2 | NA | NA | |
| 25 | 3 | 0 | ALOH | amplified LOH |
| 231 | 3 | 1 | ASCNA | allele- specific amplification |
| 5 | 3 | 2 | NA | |
| 4 | 3 | NA | NA | |
| 8 | 4 | 0 | ALOH | amplified LOH |
| 77 | 4 | 1 | ASCNA | allele- specific amplification |
| 72 | 4 | 2 | BCNA | balanced-amplification |
| 3 | 4 | NA | NA | |
| 3 | 5 | 0 | ALOH | Amplified LOH |
| 29 | 5 | 1 | ASCNA | allele- specific amplification |
| 23 | 5 | 2 | BCNA | balanced-amplification |
| 1 | 5 | 3 | NA | |
| 2 | 5 | NA | NA | |
| 4 | 6 | 1 | ASCNA | allele- specific amplification |
| 9 | 6 | 2 | BCNA | balanced-amplification |
| 7 | 6 | 3 | BCNA | balanced-amplification |
| 2 | 7 | 2 | BCNA | balanced-amplification |

**Supplementary Table 3.** Comparison between four identified SNV mutational signatures (a) and three identified indel mutational signatures (b) with previously known SNV and indels mutational signatures, respectively. SBS: Single Base Substitution; ID: indel.

| **De Novo Extracted SNVs** | **a. Global NMF SBS Signatures** | **Similarity** |
|---|---|---|
| **Signature A** | Signature **SBS-02** (8%) & Signature **SBS-13** (7%) & Signature **SBS-40** (85%) | 0.92 |
| **Signature B** | Signature **SBS-01** (10%) & Signature **SBS-05** (61%) % Signature **SBS-08** (29%) | 0.95 |
| **Signature C** | Signature **SBS-05** (40%) & Signature **SBS-40** (60%) | 0.92 |
| **Signature D** | Signature **SBS-40** (100%) | 0.94 |
| **De Novo Extracted indels** | **b. Global NMF ID Signatures** | **Similarity** |
| **Signature INDEL-A** | Signature **ID-05** (86%) & Signature **ID-08** (14%) | 0.95 |
| **Signature INDEL-B** | Signature **ID-01** (29%) & Signature **ID-02** (10%) & Signature **ID-05** (61%) | 0.93 |
| **Signature INDEL-C** | Signature **ID-03** (26%) & Signature **ID-05** (46%) & Signature **ID-06** (28%) | 0.60 |

**Supplementary Figure 1: Ploidy and purity of each sample**. Ploidy and purity of each sample was estimated based on (a) copy number and (b) density plots of single nucleotide variant allele fraction (VAF) for low purity samples. The SNV-based purity = 2*(the mode of VAF density) and is labeled in red color in panel b.

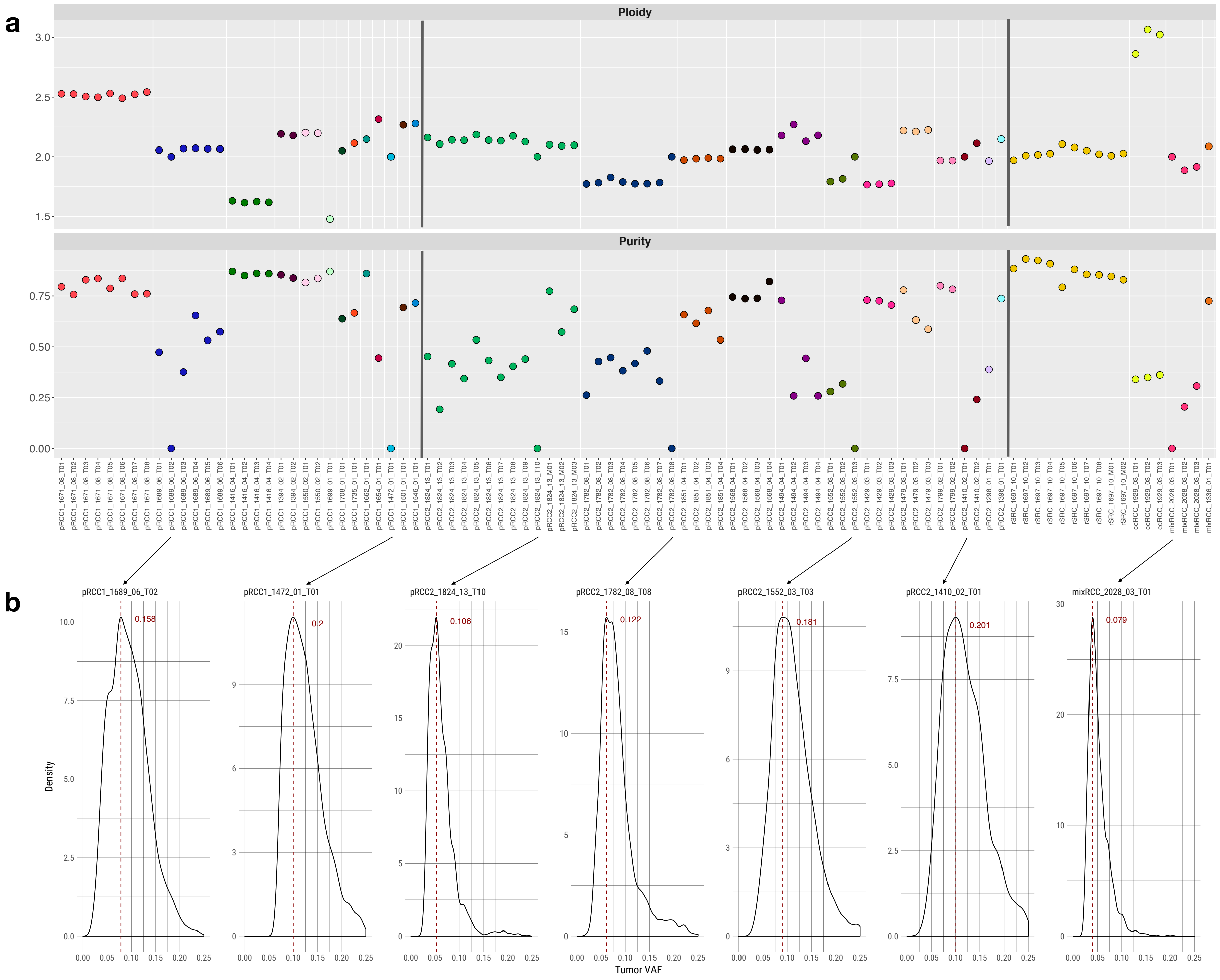

a
Ploidy
Purity
b
pRCC1_1689_06_T02
0.158
pRCC1_1472_01_T01
0.2
pRCC2_1824_13_T10
0.106
pRCC2_1782_08_T08
0.122
pRCC2_1552_03_T03
0.181
pRCC2_1410_02_T01
0.201
mixRCC_2028_03_T01
0.079
Density
Tumor VAF

**Supplementary Figure 2: Single nucleotide variants (SNVs) in known cancer driver genes.** Single nucleotide variants (SNVs) for each sample were called based on both whole-genome sequencing and deep targeted sequencing. Missense mutations, nonsense mutations and splice site mutations are annotated with different colors.

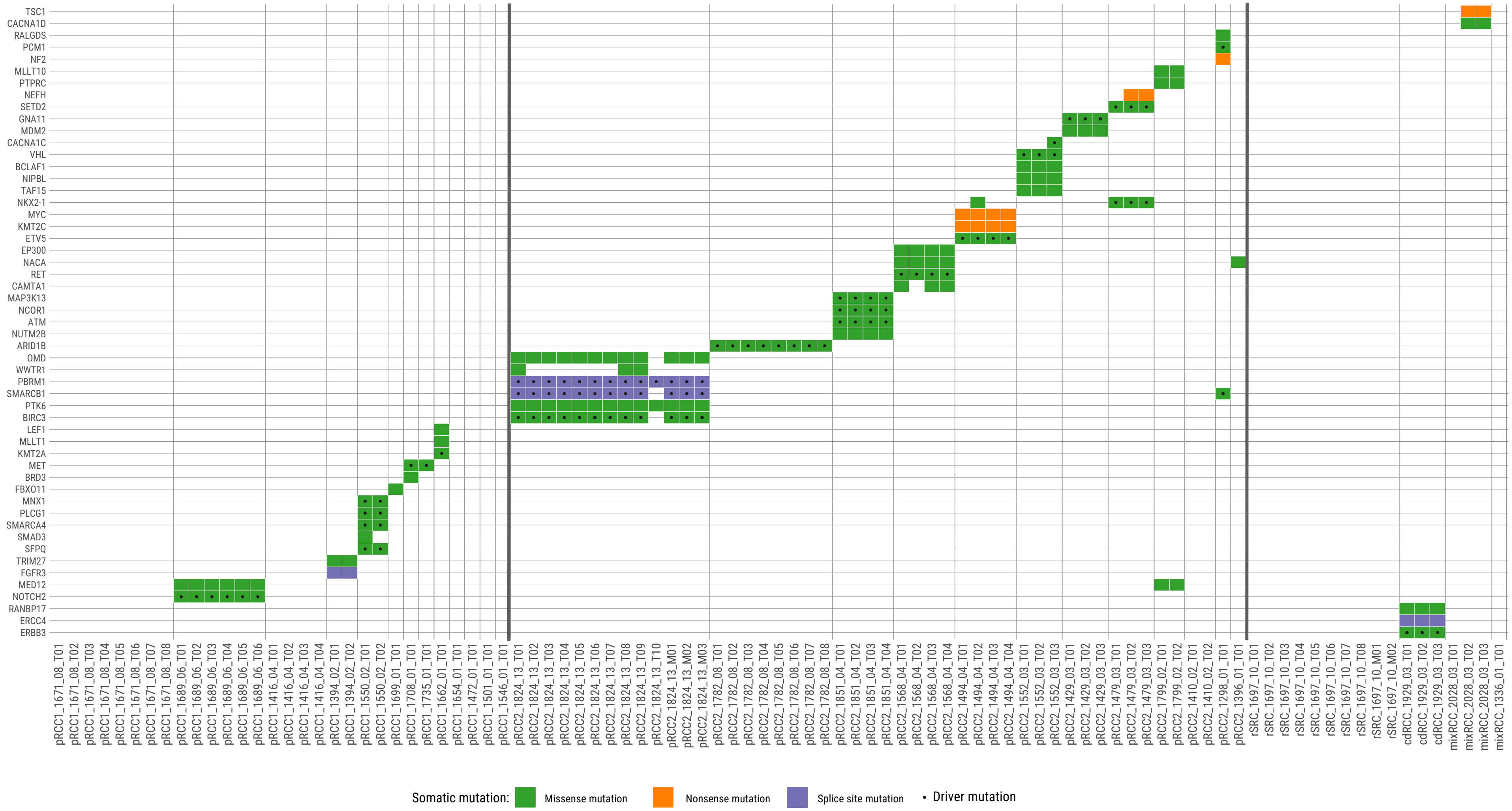
TSC1
CACNA1D
RALGDS
PCM1
NF2
MLLT10
PTPRC
NEFH
SETD2
GNA11
MDM2
CACNA1C
VHL
BCLAF1
NIPBL
TAF15
NKX2-1
MYC
KMT2C
ETV5
EP300
NACA
RET
CAMTA1
MAP3K13
NCOR1
ATM
NUTM2B
ARID1B
OMD
WWTR1
PBRM1
SMARCB1
PTK6
BIRC3
LEF1
MLLT1
KMT2A
MET
BRD3
FBXO11
MNX1
PLCG1
SMARCA4
SMAD3
SFPQ
TRIM27
FGFR3
MED12
NOTCH2
RANBP17
ERCC4
ERBB3
Somatic mutation:
Missense mutation
Nonsense mutation
Splice site mutation
Driver mutation

**Supplementary Figure 3: Phylogenetic trees and oval plots for additional tumors.**
Phylogenetic trees: the trees show the evolutionary relationships between subclones (annotated by different colors). Trunk and branch lengths are proportional to the number of substitutions in each clone cluster. Driver mutations and recurrent somatic copy number alterations are annotated on the trees. Tumor regions containing sample specific subclones are indicated on the tree leaves. Oval plots: In the top rows the ovals are ordered based on the physical sampling of the tumor regions. Ovals are nested if required by the pigeonhole principle. The first row of the plot with nested ovals is linked by lines to the ovals ordered by the phylogenetic analysis, indicating intermixing of subclones spread across 2 or more tumor regions. In the matrix, each main clone (without solid border) and subclone (with solid border) is represented as a color-coded oval. The size of the ovals is proportional to the CCF of the corresponding subclones. Each column represents a sample. Oval plots are separated by three parts: trunk (CCF = 1 in all samples), branch (present in >1 sample but not in all samples), and leaf (specific to a single sample). We also show an alternative clonality solution for tumor pRCC1_1416_04. GL: germline; amp: amplification; DLOH: hemizygous deletion loss of heterozygosity; HET: diploid heterozygous; NLOH: copy neutral loss of heterozygosity; HOMD: homozygous deletion; ASCNA: allele-specific copy number amplification; BCNA: balanced copy number amplification.

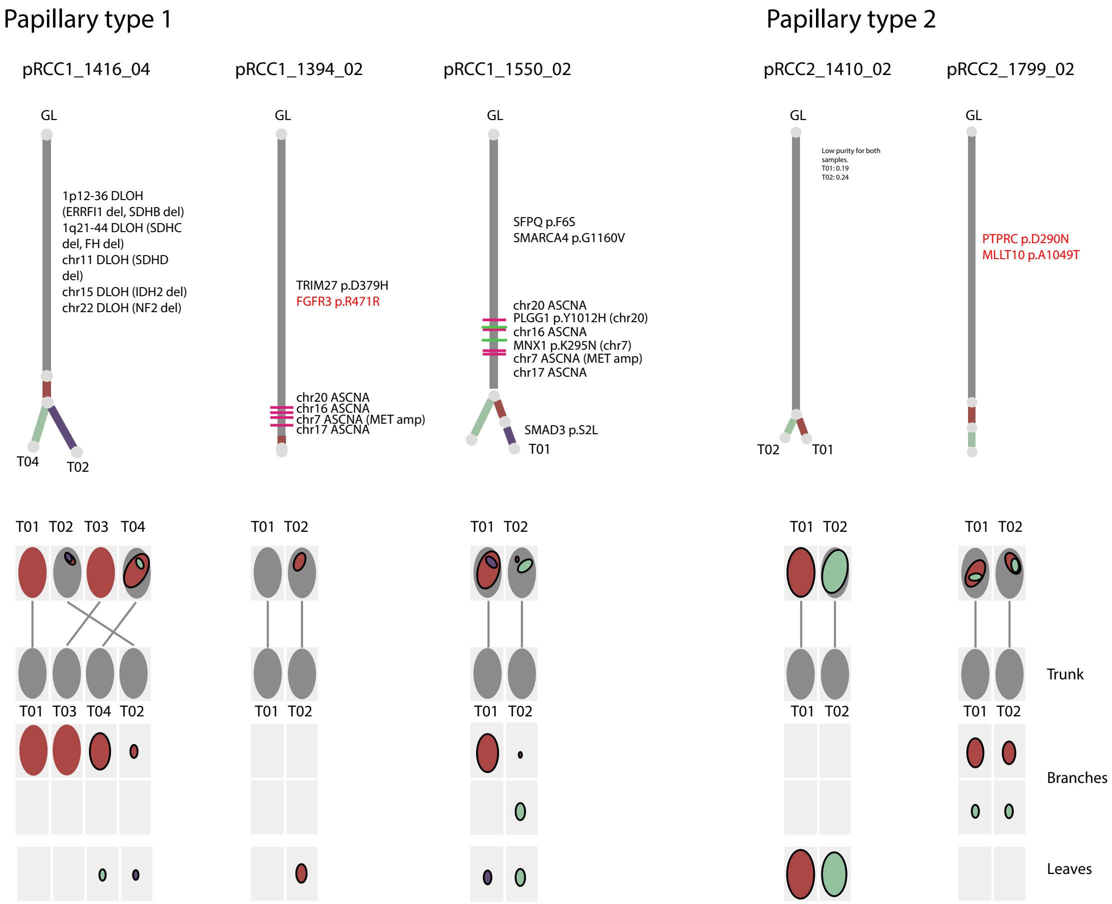
Papillary type 1
pRCC1_1416_04
GL
1p12-36 DLOH (ERRFI1 del, SDHB del)
1q21-44 DLOH (SDHC del, FH del)
chr11 DLOH (SDHD del)
chr15 DLOH (IDH2 del)
chr22 DLOH (NF2 del)
T04
T02
pRCC1_1394_02
GL
TRIM27 p.D379H
FGFR3 p.R471R
chr20 ASCNA
chr16 ASCNA
chr7 ASCNA (MET amp)
chr17 ASCNA
pRCC1_1550_02
GL
SFPQ p.F6S
SMARCA4 p.G1160V
chr20 ASCNA
PLGG1 p.Y1012H (chr20)
chr16 ASCNA
MNX1 p.K295N (chr7)
chr7 ASCNA (MET amp)
chr17 ASCNA
SMAD3 p.S2L
T01
Papillary type 2
pRCC2_1410_02
GL
Low purity for both samples.
T01: 0.19
T02: 0.24
T02
T01
pRCC2_1799_02
GL
PTPRC p.D290N
MLLT10 p.A1049T
T01 T02 T03 T04
T01 T03 T04 T02
T01 T02
T01 T02
T01 T02
T01 T02
T01 T02
T01 T02
T01 T02
T01 T02
Trunk
Branches
Leaves

**Supplementary Figure 4: Clonality of single nucleotide variants (SNVs) for each tumor**. The proportions of SNVs in trunks, internal branches and terminal branches of multi-regional trees (MRT) are presented.

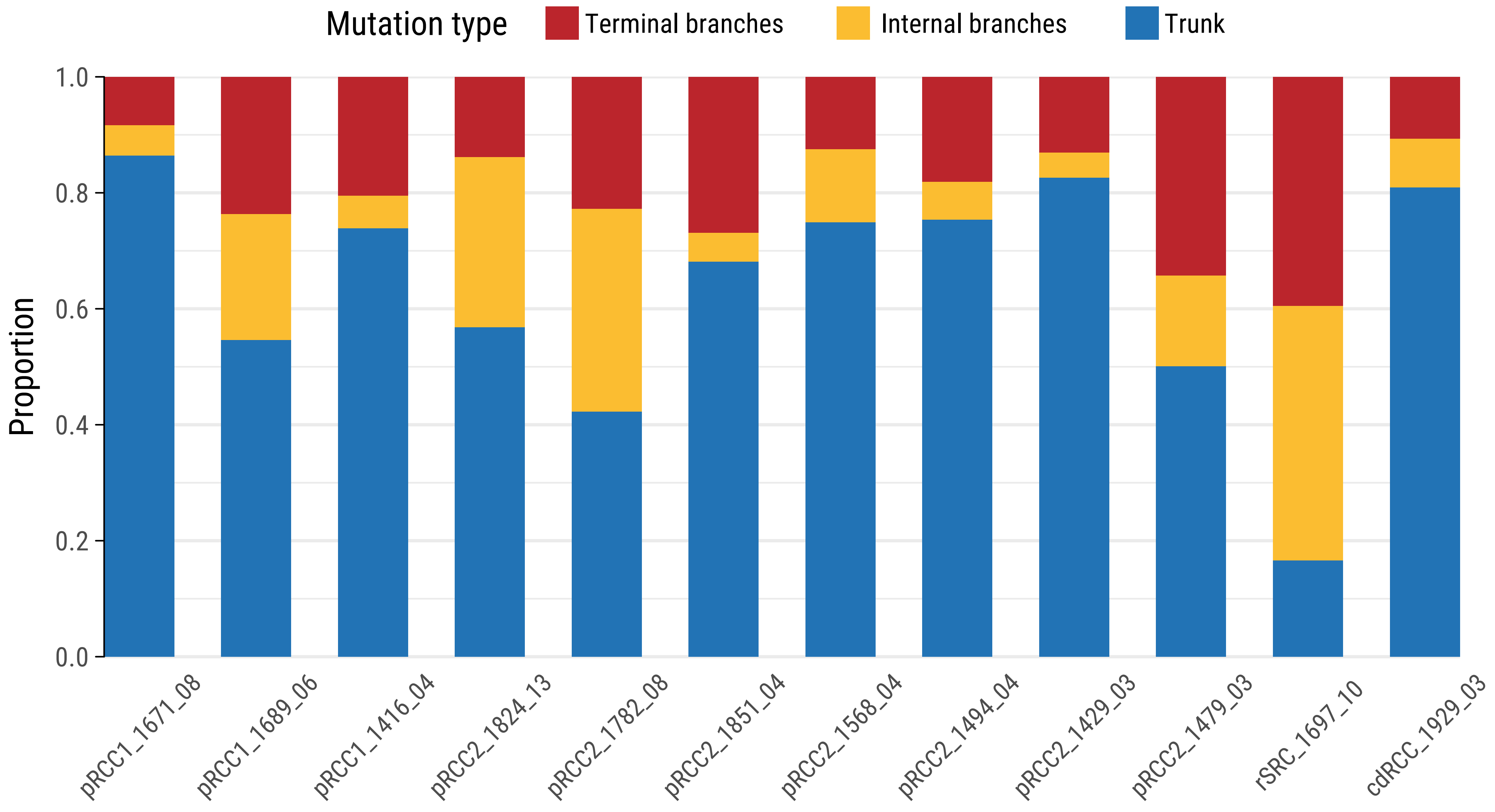
Mutation type
Terminal branches
Internal branches
Trunk
Proportion
1.0
0.8
0.6
0.4
0.2
0.0
pRCC1_1671_08
pRCC1_1689_06
pRCC1_1416_04
pRCC2_1824_13
pRCC2_1782_08
pRCC2_1851_04
pRCC2_1568_04
pRCC2_1494_04
pRCC2_1429_03
pRCC2_1479_03
rSRC_1697_10
cdRCC_1929_03

**Supplementary Figure 5: Intra-tumor heterogeneity of single nucleotide variants by genomic regions.** Genomic regions include intergenic, 1 to 5kb from the transcription starting site (TSS), promoters (0 to 1 kb from the TSS), 5'-UTRs, first exon, exon-intron boundaries, exons, introns, intron-exon boundaries, 3'-UTRs, long non-coding RNAs (lncRNAs) and enhancers (annotated by FANTOM) defined in R annotatr package (https://github.com/hhabra/annotatr).

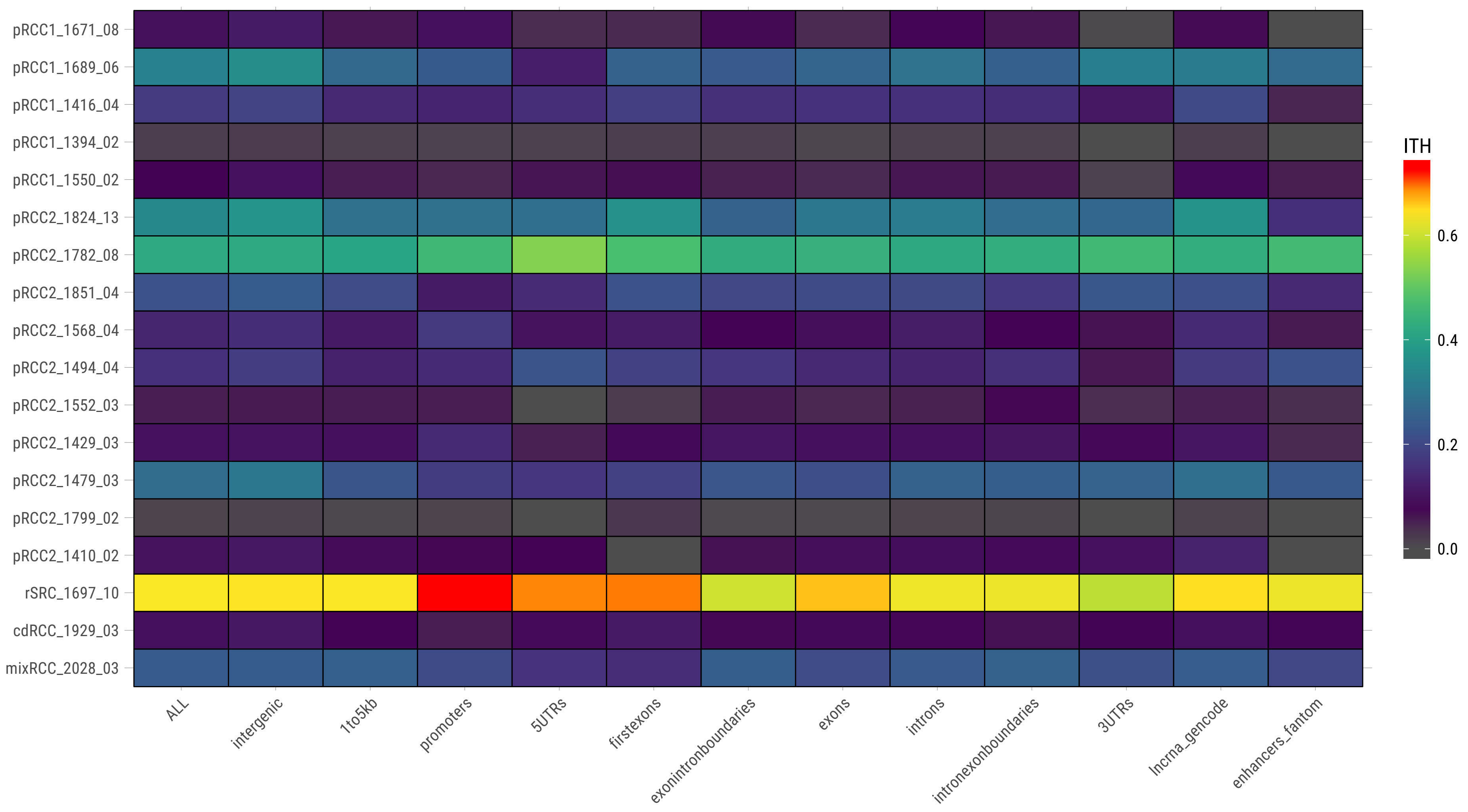

pRCC1_1671_08
pRCC1_1689_06
pRCC1_1416_04
pRCC1_1394_02
pRCC1_1550_02
pRCC2_1824_13
pRCC2_1782_08
pRCC2_1851_04
pRCC2_1568_04
pRCC2_1494_04
pRCC2_1552_03
pRCC2_1429_03
pRCC2_1479_03
pRCC2_1799_02
pRCC2_1410_02
rSRC_1697_10
cdRCC_1929_03
mixRCC_2028_03
ALL
intergenic
1to5kb
promoters
5UTRs
firstexons
exonintronboundaries
exons
introns
intronexonboundaries
3UTRs
lncrna_gencode
enhancers_fantom
ITH
0.6
0.4
0.2
0.0

**Supplementary Figure 6: Percentage of somatic copy number alterations (SCNA) across cytoband regions.** HOMD: homozygous deletion; DLOH: hemizygous deletion loss of heterozygosity; NLOH: copy neutral loss of heterozygosity; ALOH: amplified loss of heterozygosity; ASCNA: allele-specific copy number amplification; BCNA: balanced copy number amplification.

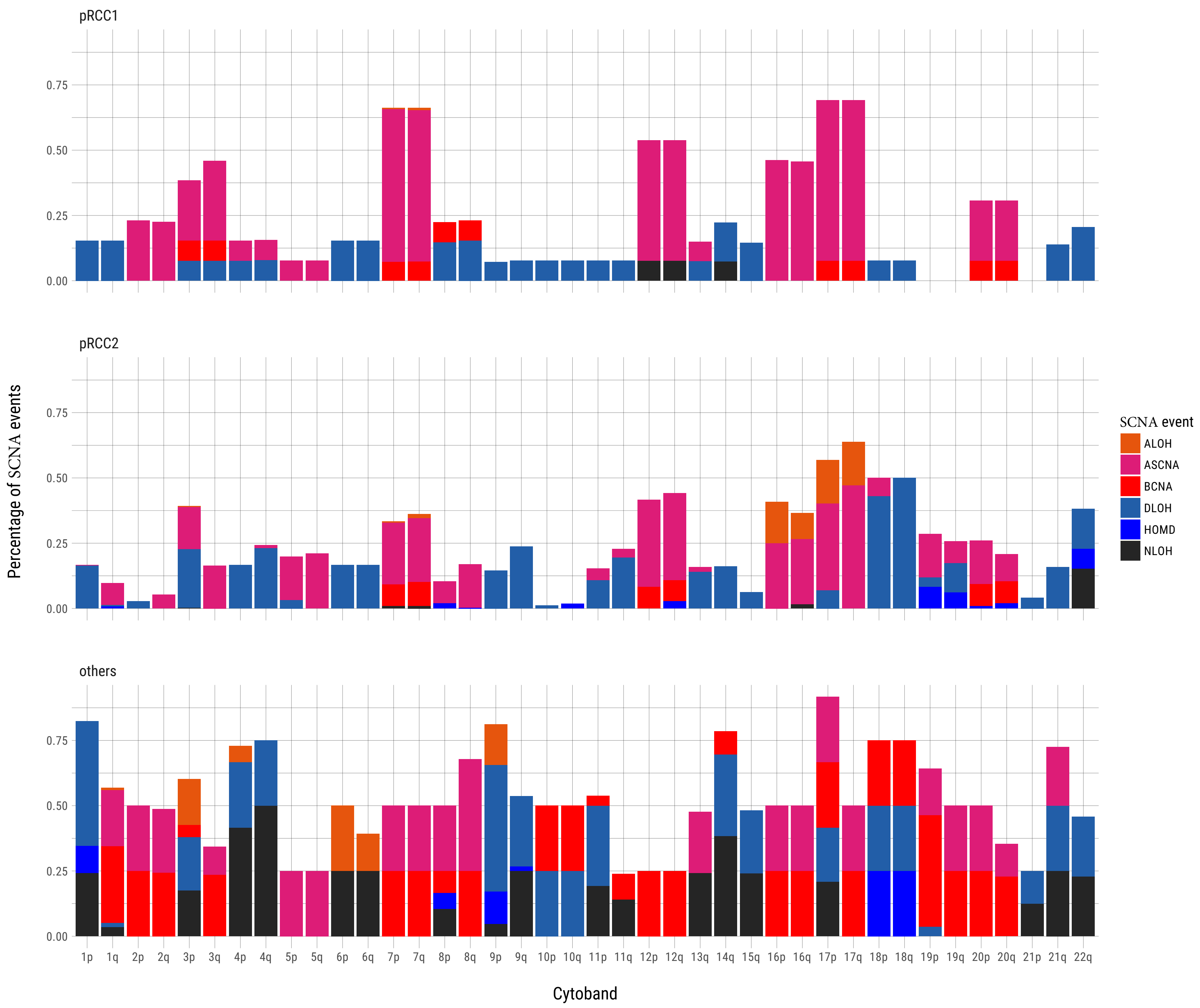
pRCC1
pRCC2
others
Percentage of SCNA events
0.00
0.25
0.50
0.75
1p 1q 2p 2q 3p 3q 4p 4q 5p 5q 6p 6q 7p 7q 8p 8q 9p 9q 10p 10q 11p 11q 12p 12q 13q 14q 15q 16p 16q 17p 17q 18p 18q 19p 19q 20p 20q 21p 21q 22q
Cytoband
SCNA event
ALOH
ASCNA
BCNA
DLOH
HOMD
NLOH

**Supplementary Figure 7: Genome-wide profiles of SCNA events of each sample.** Samples are labeled by copy number type, histology subgroup, tissue type and purity. HOMD: homozygous deletion; DLOH: hemizygous deletion loss of heterozygosity; HET: diploid heterozygous; NLOH: copy neutral loss of heterozygosity; ALOH: amplified loss of heterozygosity; ASCNA: allele-specific copy number amplification; BCNA: balanced copy number amplification.

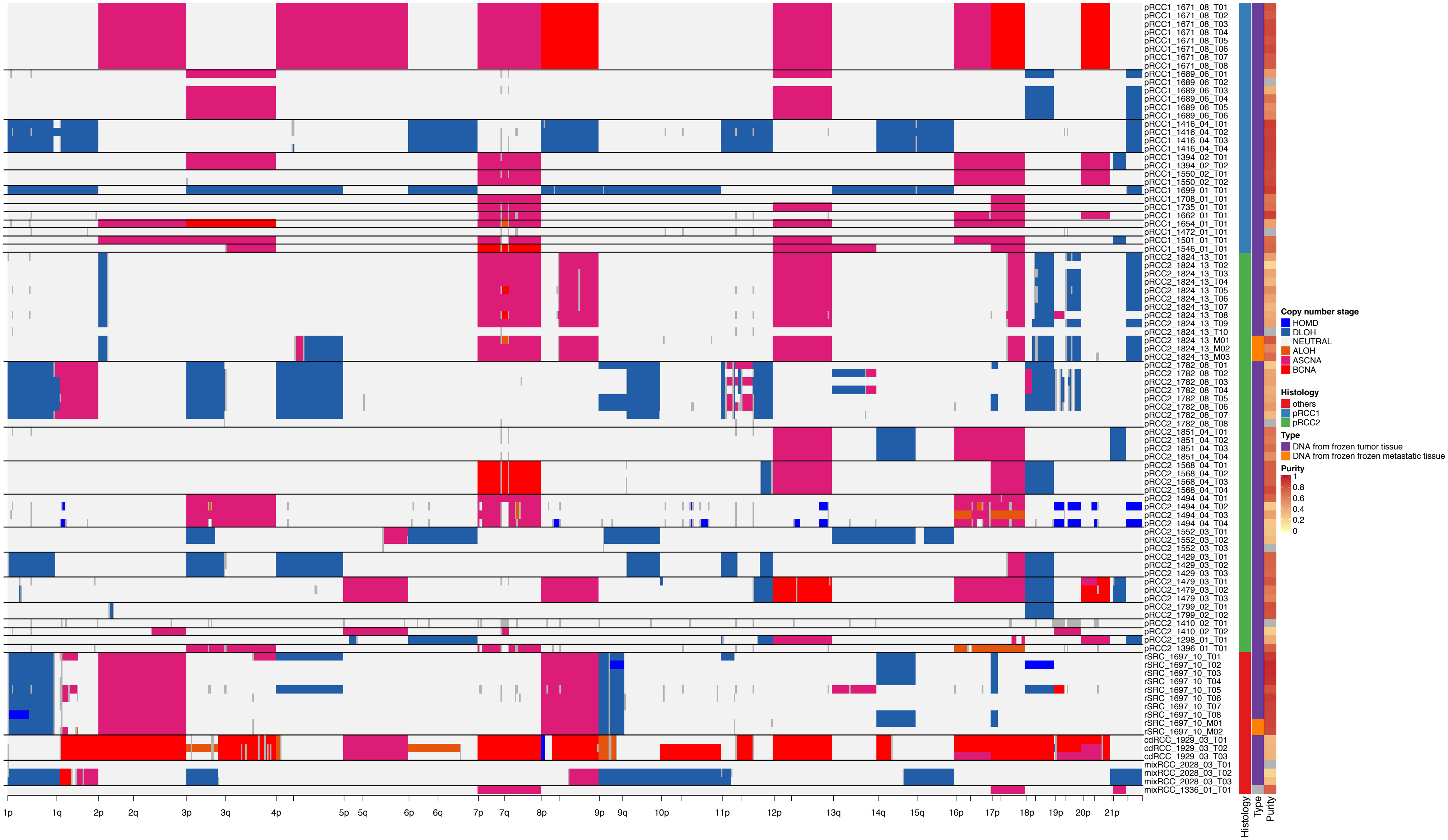

pRCC1_1671_08_T01
pRCC1_1671_08_T02
pRCC1_1671_08_T03
pRCC1_1671_08_T04
pRCC1_1671_08_T05
pRCC1_1671_08_T06
pRCC1_1671_08_T07
pRCC1_1671_08_T08
pRCC1_1689_06_T01
pRCC1_1689_06_T02
pRCC1_1689_06_T03
pRCC1_1689_06_T04
pRCC1_1689_06_T05
pRCC1_1689_06_T06
pRCC1_1416_04_T01
pRCC1_1416_04_T02
pRCC1_1416_04_T03
pRCC1_1416_04_T04
pRCC1_1394_02_T01
pRCC1_1394_02_T02
pRCC1_1550_02_T01
pRCC1_1550_02_T02
pRCC1_1699_01_T01
pRCC1_1708_01_T01
pRCC1_1735_01_T01
pRCC1_1662_01_T01
pRCC1_1654_01_T01
pRCC1_1472_01_T01
pRCC1_1501_01_T01
pRCC1_1546_01_T01
pRCC2_1824_13_T01
pRCC2_1824_13_T02
pRCC2_1824_13_T03
pRCC2_1824_13_T04
pRCC2_1824_13_T05
pRCC2_1824_13_T06
pRCC2_1824_13_T07
pRCC2_1824_13_T08
pRCC2_1824_13_T09
pRCC2_1824_13_T10
pRCC2_1824_13_M01
pRCC2_1824_13_M02
pRCC2_1824_13_M03
pRCC2_1782_08_T01
pRCC2_1782_08_T02
pRCC2_1782_08_T03
pRCC2_1782_08_T04
pRCC2_1782_08_T05
pRCC2_1782_08_T06
pRCC2_1782_08_T07
pRCC2_1782_08_T08
pRCC2_1851_04_T01
pRCC2_1851_04_T02
pRCC2_1851_04_T03
pRCC2_1851_04_T04
pRCC2_1568_04_T01
pRCC2_1568_04_T02
pRCC2_1568_04_T03
pRCC2_1568_04_T04
pRCC2_1494_04_T01
pRCC2_1494_04_T02
pRCC2_1494_04_T03
pRCC2_1494_04_T04
pRCC2_1552_03_T01
pRCC2_1552_03_T02
pRCC2_1552_03_T03
pRCC2_1429_03_T01
pRCC2_1429_03_T02
pRCC2_1429_03_T03
pRCC2_1479_03_T01
pRCC2_1479_03_T02
pRCC2_1479_03_T03
pRCC2_1799_02_T01
pRCC2_1799_02_T02
pRCC2_1410_02_T01
pRCC2_1410_02_T02
pRCC2_1298_01_T01
pRCC2_1396_01_T01
rSRC_1697_10_T01
rSRC_1697_10_T02
rSRC_1697_10_T03
rSRC_1697_10_T04
rSRC_1697_10_T05
rSRC_1697_10_T06
rSRC_1697_10_T07
rSRC_1697_10_T08
rSRC_1697_10_M01
rSRC_1697_10_M02
cdRCC_1929_03_T01
cdRCC_1929_03_T02
cdRCC_1929_03_T03
mixRCC_2028_03_T01
mixRCC_2028_03_T02
mixRCC_2028_03_T03
mixRCC_1336_01_T01
1p
1q
2p
2q
3p
3q
4p
5p
5q
6p
6q
7p
7q
8p
9p
9q
10p
11p
12p
13q
14q
15q
16p
17p
18p
19p
20p
21p
Histology
Type
Purity
Copy number stage
HOMD
DLOH
NEUTRAL
ALOH
ASCNA
BCNA
Histology
others
pRCC1
pRCC2
Type
DNA from frozen tumor tissue
DNA from frozen frozen metastatic tissue
Purity
1
0.8
0.6
0.4
0.2
0

**Supplementary Figure 8: Concordance of purity and ploidy across platforms.** The scatterplots of purity (left panel) and ploidy (middle panel) are estimated based on WGS and SNP genotyping. Right panel shows the concordance of overlapped arm-level SCNA regions across platforms for each sample.

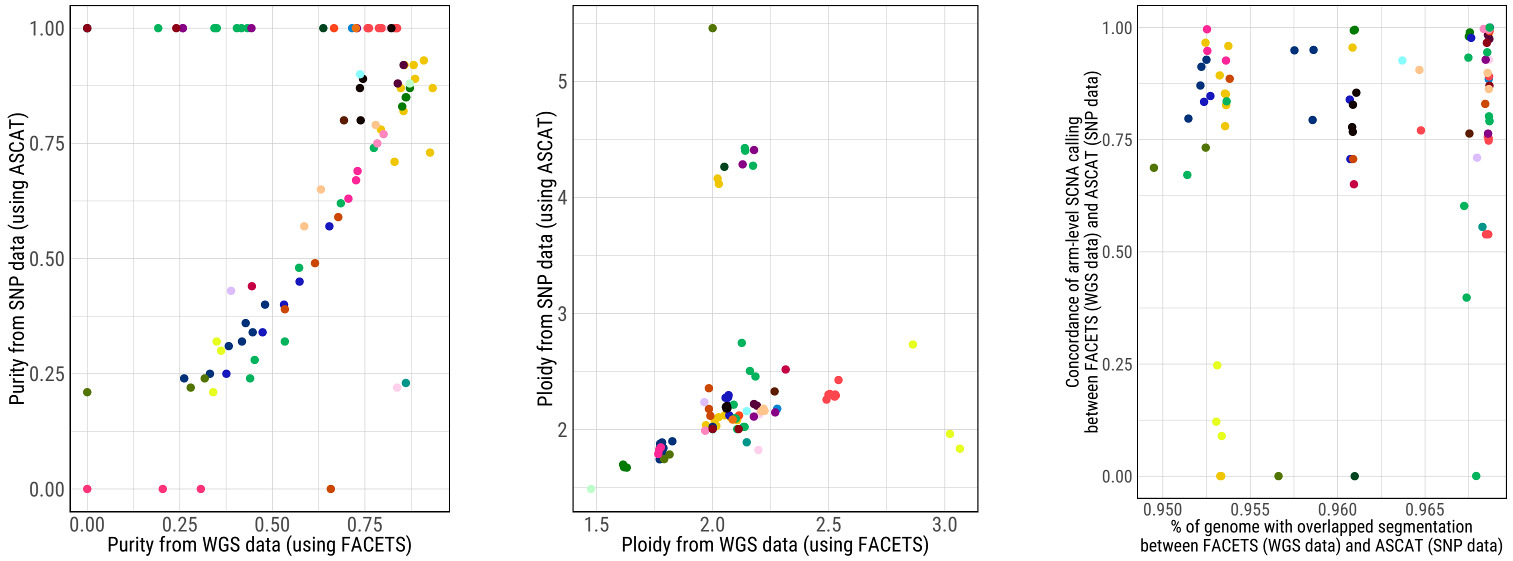
Purity from SNP data (using ASCAT)
Purity from WGS data (using FACETS)
Ploidy from SNP data (using ASCAT)
Ploidy from WGS data (using FACETS)
Concordance of arm-level SCNA calling between FACETS (WGS data) and ASCAT (SNP data)
% of genome with overlapped segmentation between FACETS (WGS data) and ASCAT (SNP data)

**Supplementary Figure 9: Cluster Dendrogram of SCNA profiles.** The similar SCNA profiles were
clustered hierarchically.

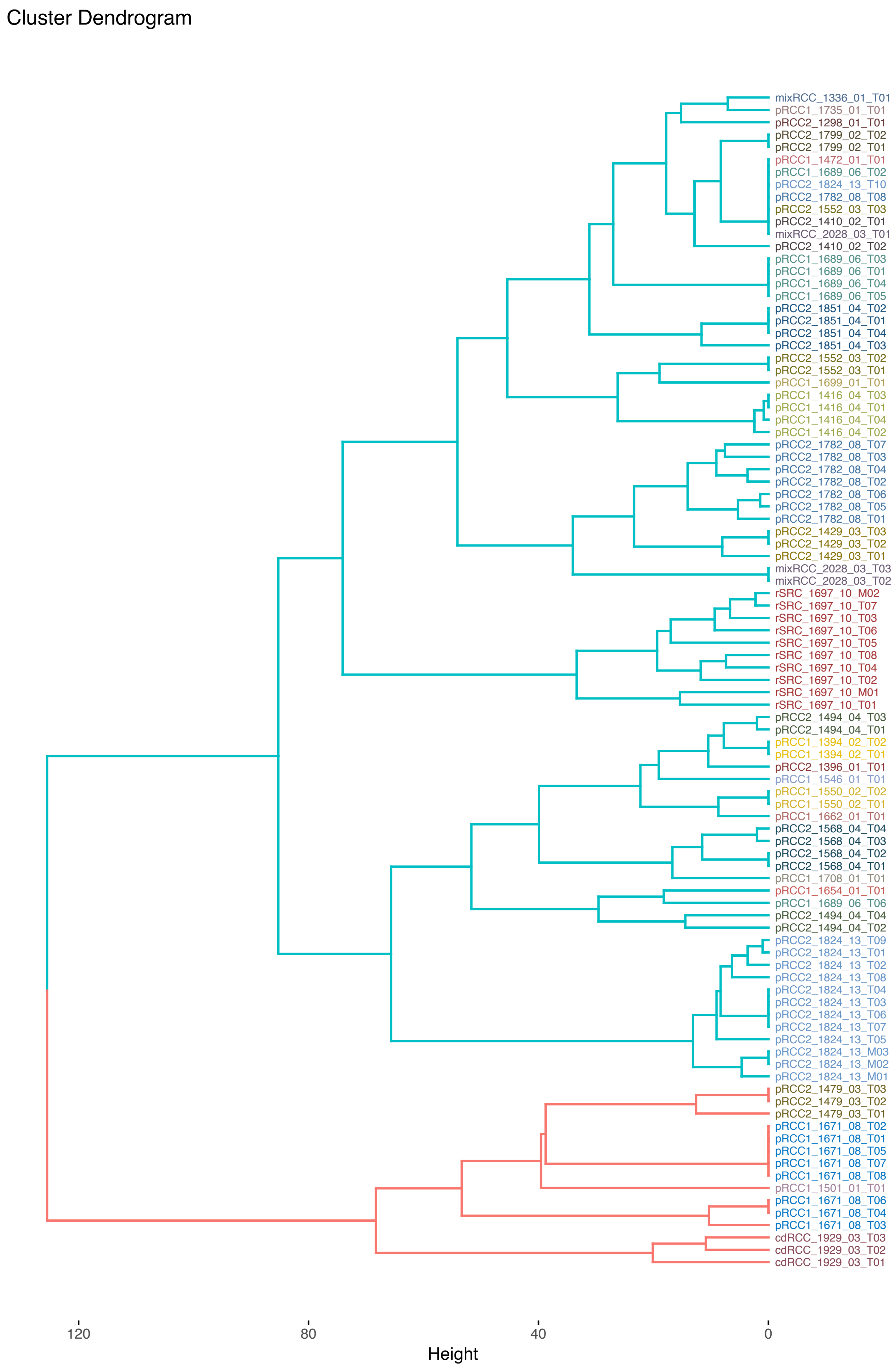
Cluster Dendrogram
mixRCC_1336_01_T01
pRCC1_1735_01_T01
pRCC2_1298_01_T01
pRCC2_1799_02_T02
pRCC2_1799_02_T01
pRCC1_1472_01_T01
pRCC1_1689_06_T02
pRCC2_1824_13_T10
pRCC2_1782_08_T08
pRCC2_1552_03_T03
pRCC2_1410_02_T01
mixRCC_2028_03_T01
pRCC2_1410_02_T02
pRCC1_1689_06_T03
pRCC1_1689_06_T01
pRCC1_1689_06_T04
pRCC1_1689_06_T05
pRCC2_1851_04_T02
pRCC2_1851_04_T01
pRCC2_1851_04_T04
pRCC2_1851_04_T03
pRCC2_1552_03_T02
pRCC2_1552_03_T01
pRCC1_1699_01_T01
pRCC1_1416_04_T03
pRCC1_1416_04_T01
pRCC1_1416_04_T04
pRCC1_1416_04_T02
pRCC2_1782_08_T07
pRCC2_1782_08_T03
pRCC2_1782_08_T04
pRCC2_1782_08_T02
pRCC2_1782_08_T06
pRCC2_1782_08_T05
pRCC2_1782_08_T01
pRCC2_1429_03_T03
pRCC2_1429_03_T02
pRCC2_1429_03_T01
mixRCC_2028_03_T03
mixRCC_2028_03_T02
rSRC_1697_10_M02
rSRC_1697_10_T07
rSRC_1697_10_T03
rSRC_1697_10_T06
rSRC_1697_10_T05
rSRC_1697_10_T08
rSRC_1697_10_T04
rSRC_1697_10_T02
rSRC_1697_10_M01
rSRC_1697_10_T01
pRCC2_1494_04_T03
pRCC2_1494_04_T01
pRCC1_1394_02_T02
pRCC1_1394_02_T01
pRCC2_1396_01_T01
pRCC1_1546_01_T01
pRCC1_1550_02_T02
pRCC1_1550_02_T01
pRCC1_1662_01_T01
pRCC2_1568_04_T04
pRCC2_1568_04_T03
pRCC2_1568_04_T02
pRCC2_1568_04_T01
pRCC1_1708_01_T01
pRCC1_1654_01_T01
pRCC1_1689_06_T06
pRCC2_1494_04_T04
pRCC2_1494_04_T02
pRCC2_1824_13_T09
pRCC2_1824_13_T01
pRCC2_1824_13_T02
pRCC2_1824_13_T08
pRCC2_1824_13_T04
pRCC2_1824_13_T03
pRCC2_1824_13_T06
pRCC2_1824_13_T07
pRCC2_1824_13_T05
pRCC2_1824_13_M03
pRCC2_1824_13_M02
pRCC2_1824_13_M01
pRCC2_1479_03_T03
pRCC2_1479_03_T02
pRCC2_1479_03_T01
pRCC1_1671_08_T02
pRCC1_1671_08_T01
pRCC1_1671_08_T05
pRCC1_1671_08_T07
pRCC1_1671_08_T08
pRCC1_1501_01_T01
pRCC1_1671_08_T06
pRCC1_1671_08_T04
pRCC1_1671_08_T03
cdRCC_1929_03_T03
cdRCC_1929_03_T02
cdRCC_1929_03_T01
120
80
40
0
Height

**Supplementary Figure 10: Focal SCNA events for each sample.** DLOH: hemizygous deletion loss of heterozygosity; HET: diploid heterozygous; NLOH: copy neutral loss of heterozygosity; ALOH: amplified loss of heterozygosity; ASCNA: allele-specific copy number amplification; BCNA: balanced copy number amplification.

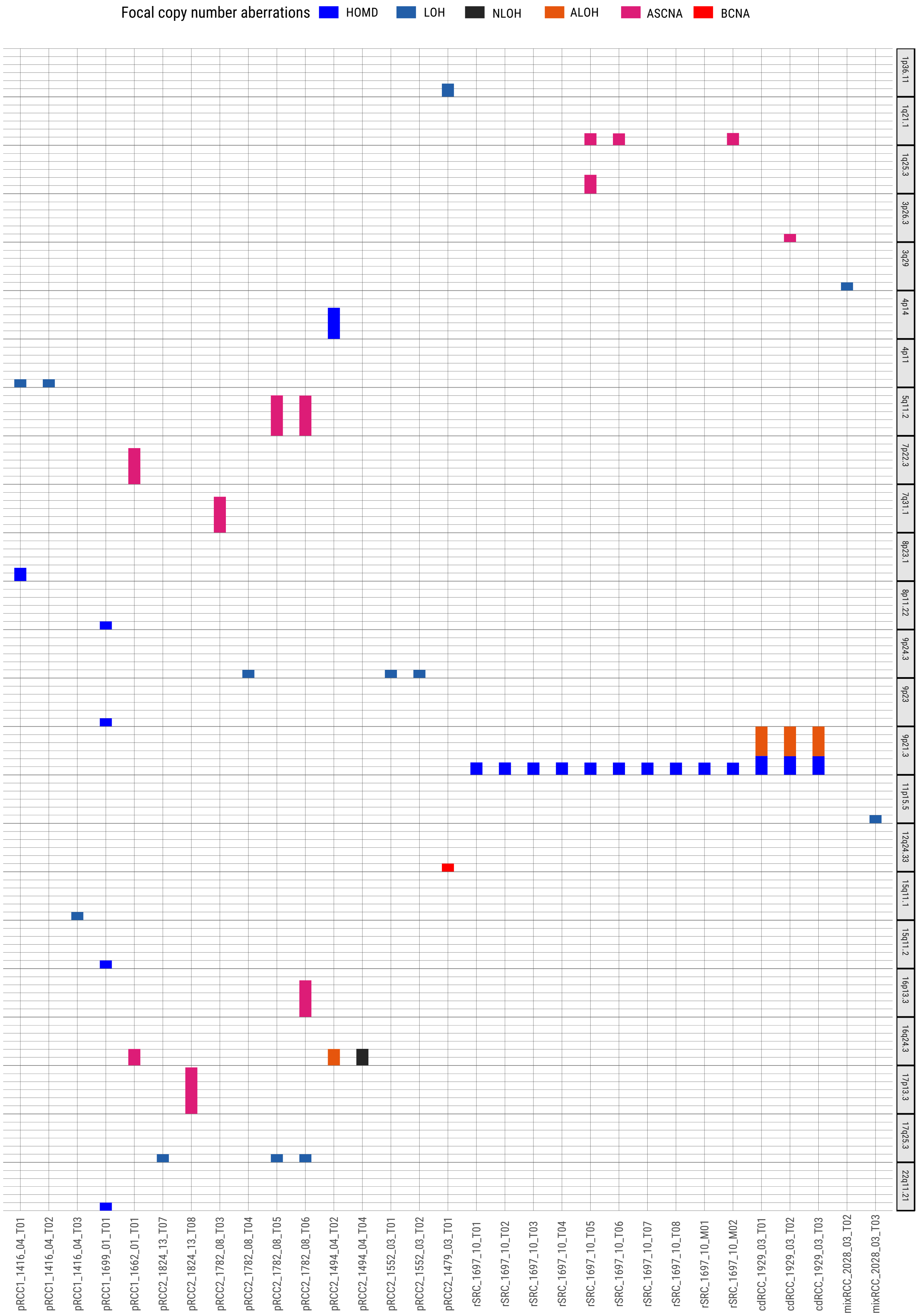
Focal copy number aberrations
HOMD
LOH
NLOH
ALOH
ASCNA
BCNA
1p36.11
1q21.1
1q25.3
3p26.3
3q29
4p14
4p11
5q11.2
7p22.3
7q31.1
8p23.1
8p11.22
9p24.3
9p23
9p21.3
11p15.5
12q24.33
15q11.1
15q11.2
16p13.3
16q24.3
17p13.3
17q25.3
22q11.21
pRCC1_1416_04_T01
pRCC1_1416_04_T02
pRCC1_1416_04_T03
pRCC1_1699_01_T01
pRCC1_1662_01_T01
pRCC2_1824_13_T07
pRCC2_1824_13_T08
pRCC2_1782_08_T03
pRCC2_1782_08_T04
pRCC2_1782_08_T05
pRCC2_1782_08_T06
pRCC2_1494_04_T02
pRCC2_1494_04_T04
pRCC2_1552_03_T01
pRCC2_1552_03_T02
pRCC2_1479_03_T01
rSRC_1697_10_T01
rSRC_1697_10_T02
rSRC_1697_10_T03
rSRC_1697_10_T04
rSRC_1697_10_T05
rSRC_1697_10_T06
rSRC_1697_10_T07
rSRC_1697_10_T08
rSRC_1697_10_M01
rSRC_1697_10_M02
cdRCC_1929_03_T01
cdRCC_1929_03_T02
cdRCC_1929_03_T03
mixRCC_2028_03_T02
mixRCC_2028_03_T03

**Supplementary Figure 11: rSRC copy number profile for chromosome 9.** The clonal focal homozygous deletion of *CDKN2A* is located at 9p21.3. For each sample, the blue dots are observed values and red lined are estimated ones; The first two panels show the profiles of logR and logOR over chromosomes; The last panel indicates estimated total copy numbers and minor copy number over chromosomes with cancer cell fraction (cf-em) in bottom, estimated by the expectation– maximization (em) algorithm.

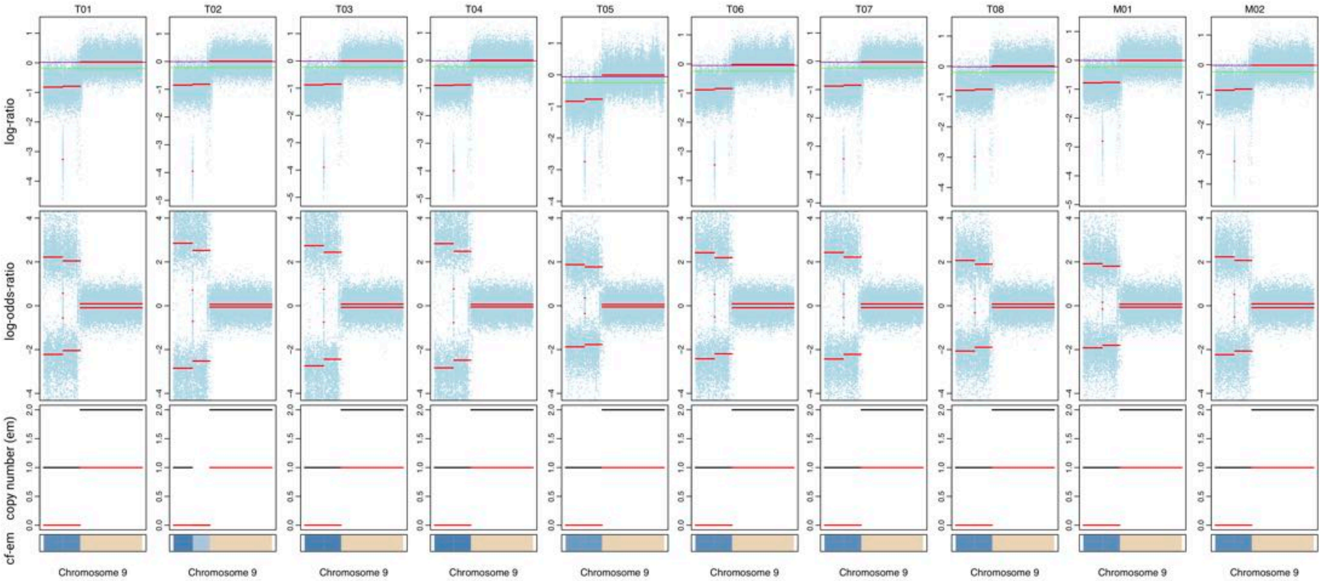

T01
T02
T03
T04
T05
T06
T07
T08
M01
M02
log-ratio
log-odds-ratio
copy number (em)
cf-em
Chromosome 9

**Supplementary Figure 12: Validation of gene fusions.** RT-PCR to detect putative fusions were performed using indicated primer sets and samples (Panel a). Individual amplicons indicated by arrows in Panel b were gel purified and sequenced by Sanger sequencing. Numbers at arrows indicate approximate molecular weights. Fusions and breakpoints detected by WGS between EWSR/PATZ1 and MALAT1/TFEB were validated by this method. The MET-MET product amplified by primer set B is the normally annotated transcript without the putative indel. RT-PCR was repeated twice for reproducibility from the same cDNA library.

**(a)**

| Forward Primer | | Reverse Primer | | | Reference |
|---|---|---|---|---|---|
| METex1F | GTTCCTGGGCACCGAAAG | METex4R | CACATTTGTCGTGGCACCAG | | novel |
| METex2F-2 | TGCCATGTGTGCATTCCCTA | METex4R | | | novel |
| STRN-ALKF | CCACAAGTTGAAATACGGGACAGAA | STRN-ALKR | TCAGCTTGTACTCAGGGCTCT | | http://www.pnas.org/content/111/11/4233.short |
| ALKex19F | TGATCCTCTCTGTGGTGACCT | STRNex4R | GTGGCTGCACTTCTGTTTCA | | http://journals.plos.org/plosone/article?id=10.1371/journal.pone.0087170#pone.0087170.s002 |
| EWS-720F | ATGGTCAACAAAGCAGCTATGGG | ZSG-S5R | GTCAGGAACCGAATGGGACGA | | Oncogene 2000 19 3799-3804 Mastrangelo et al |
| EWS-720F | | ZSG-A1R | GCAGGGCACCTTGTCTTCATG | | Oncogene 2000 19 3799-3804 Mastrangelo et al |
| EWS-720F | | PATZ1ex1R | GAGCTGGAGATGCACACTATCAG | | novel |
| AlphaRT-1f | TAA CGC ATT TAC TAA ACG CAG ACG | TFEBexon2r | AAC CCT ATG CGT GAC GCC ATG GTG G | | Ian J. Davis, 6051–6056, doi: 10.1073/pnas.0931430100 |

**(b)**

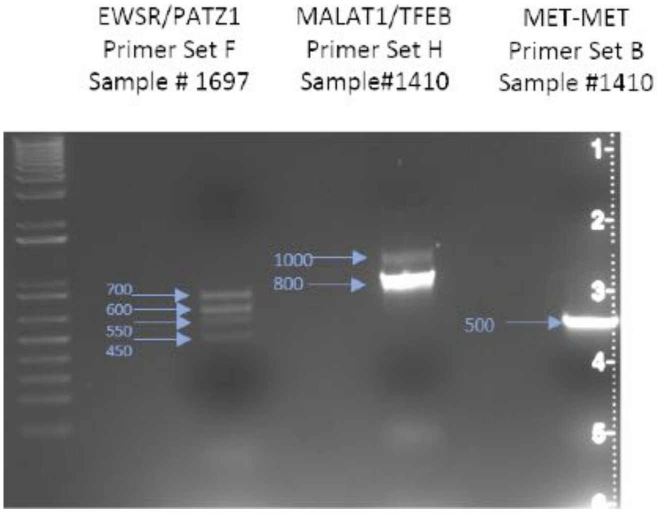

**Supplementary Figure 13: SV hotspots in rSRC_1697_10.** Each arch links the breakpoints of SV fragments with colors denoting different SV types.

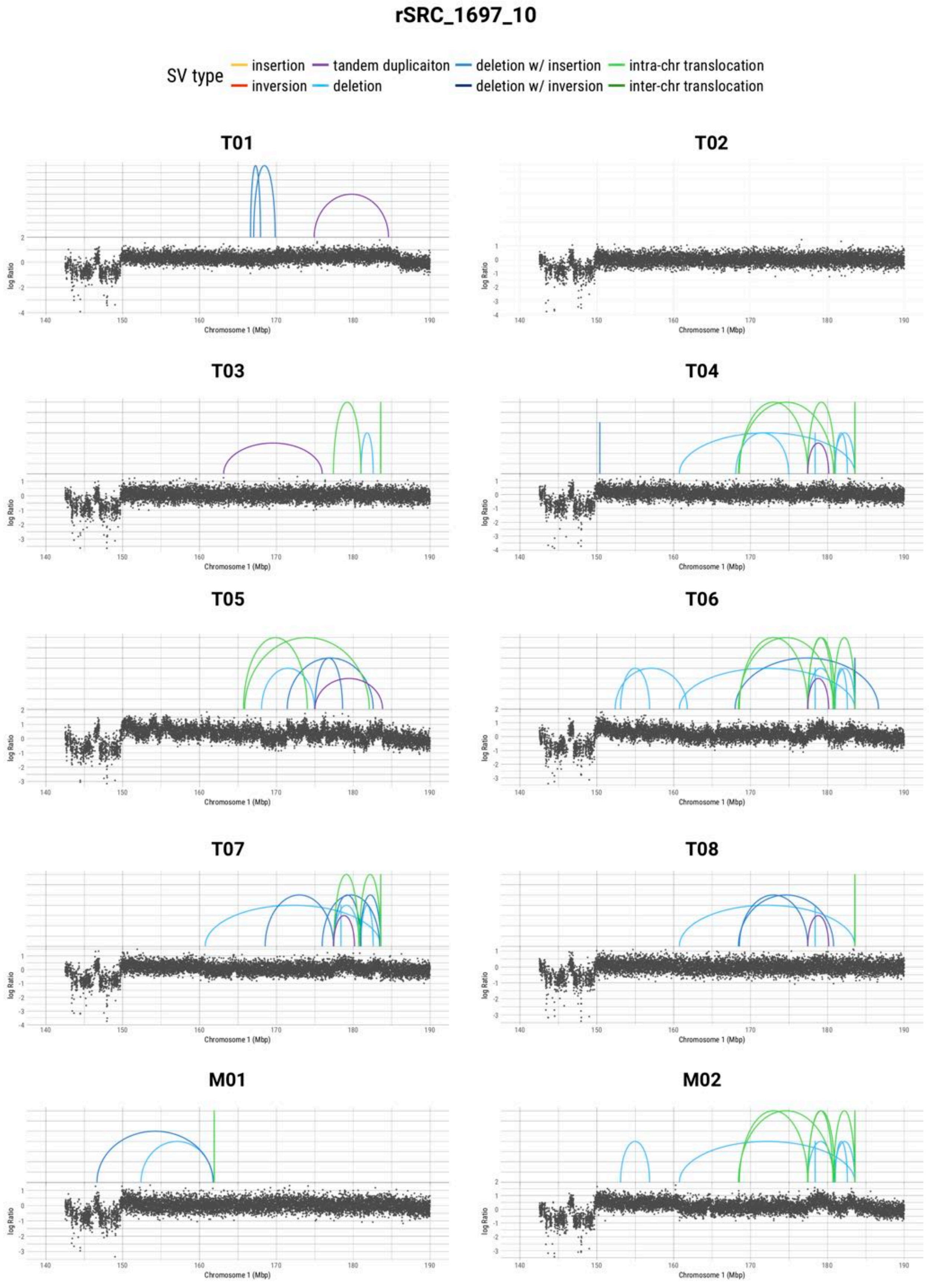

rSRC_1697_10
SV type
insertion
inversion
tandem duplicaiton
deletion
deletion w/ insertion
deletion w/ inversion
intra-chr translocation
inter-chr translocation
T01
T02
T03
T04
T05
T06
T07
T08
M01
M02
log Ratio
Chromosome 1 (Mbp)

**Supplementary Figure 14: LINE-1 retrotransposition events in pRCC2_1782_08.** Genes involved in the retrotransposons insertions are indicated.

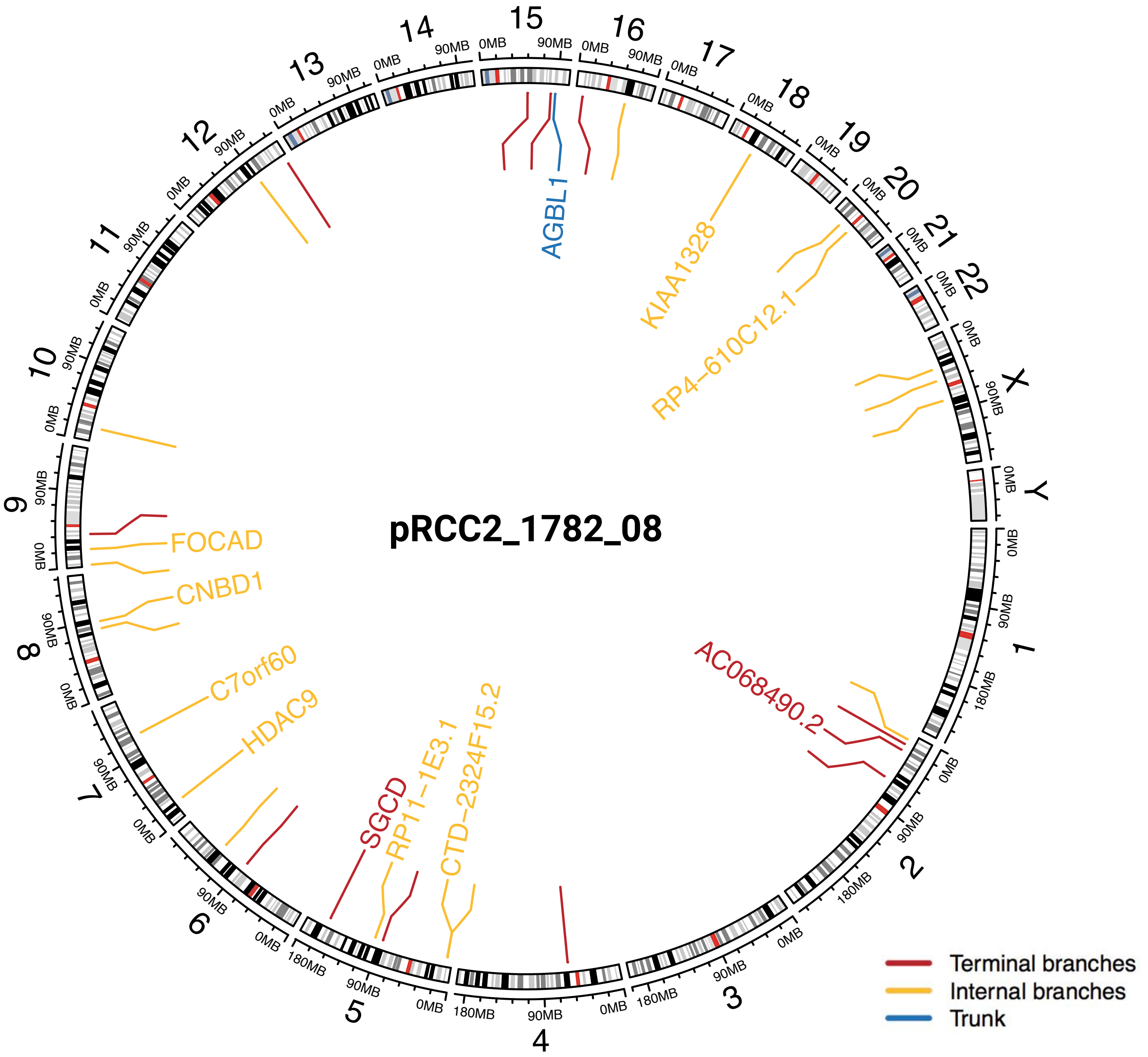

pRCC2_1782_08
AGBL1
KIAA1328
RP4-610C12.1
FOCAD
CNBD1
C7orf60
HDAC9
SGCD
RP11-1E3.1
CTD-2324F15.2
AC068490.2
Terminal branches
Internal branches
Trunk

**Supplementary Figure 15: Circos plots of SV (linked by arches) and SCNA (in the inner circle) for each tumor.** Branch and trunk events are notated by different colors.

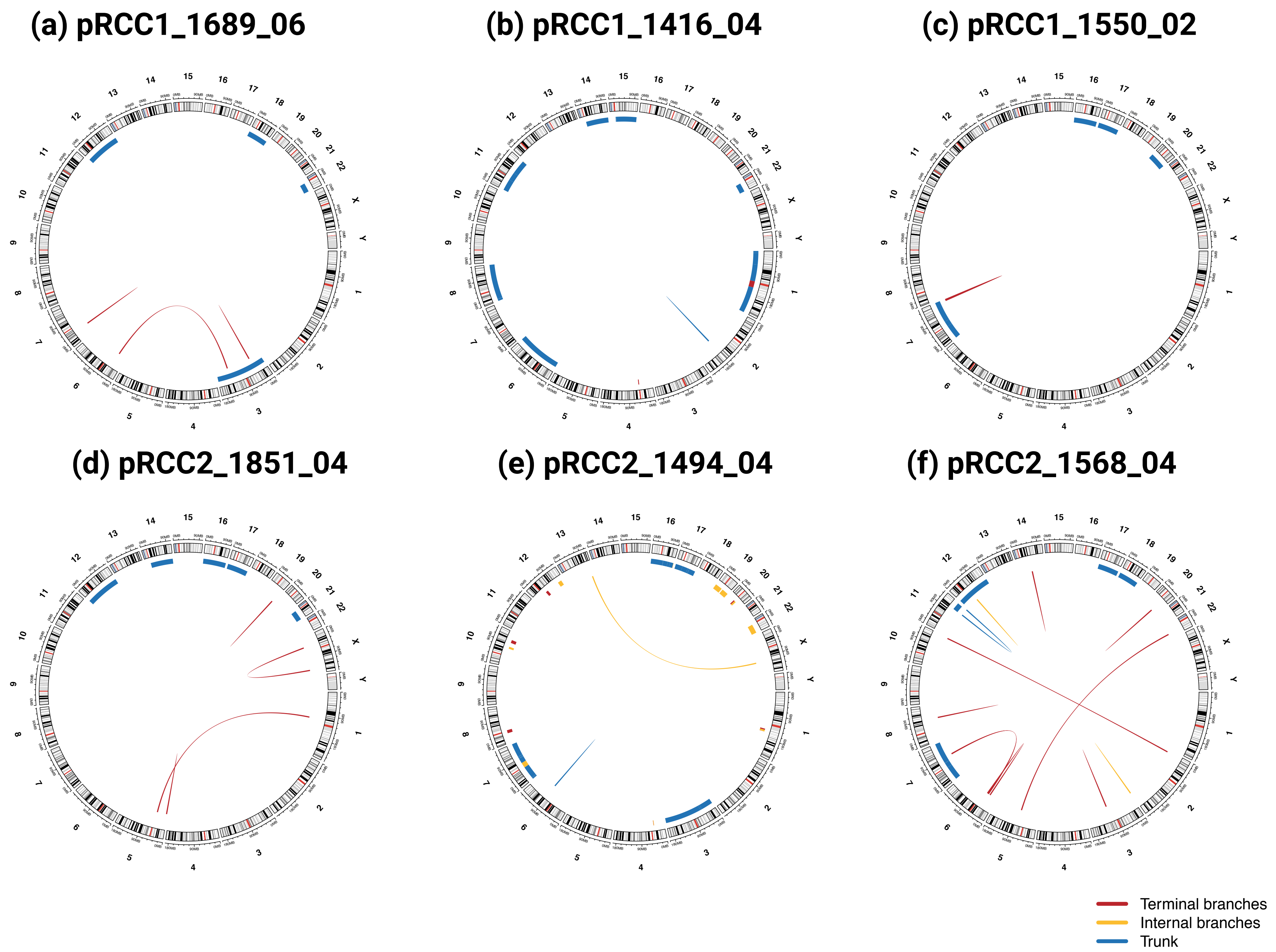

(a) pRCC1_1689_06
(b) pRCC1_1416_04
(c) pRCC1_1550_02
(d) pRCC2_1851_04
(e) pRCC2_1494_04
(f) pRCC2_1568_04
Terminal branches
Internal branches
Trunk

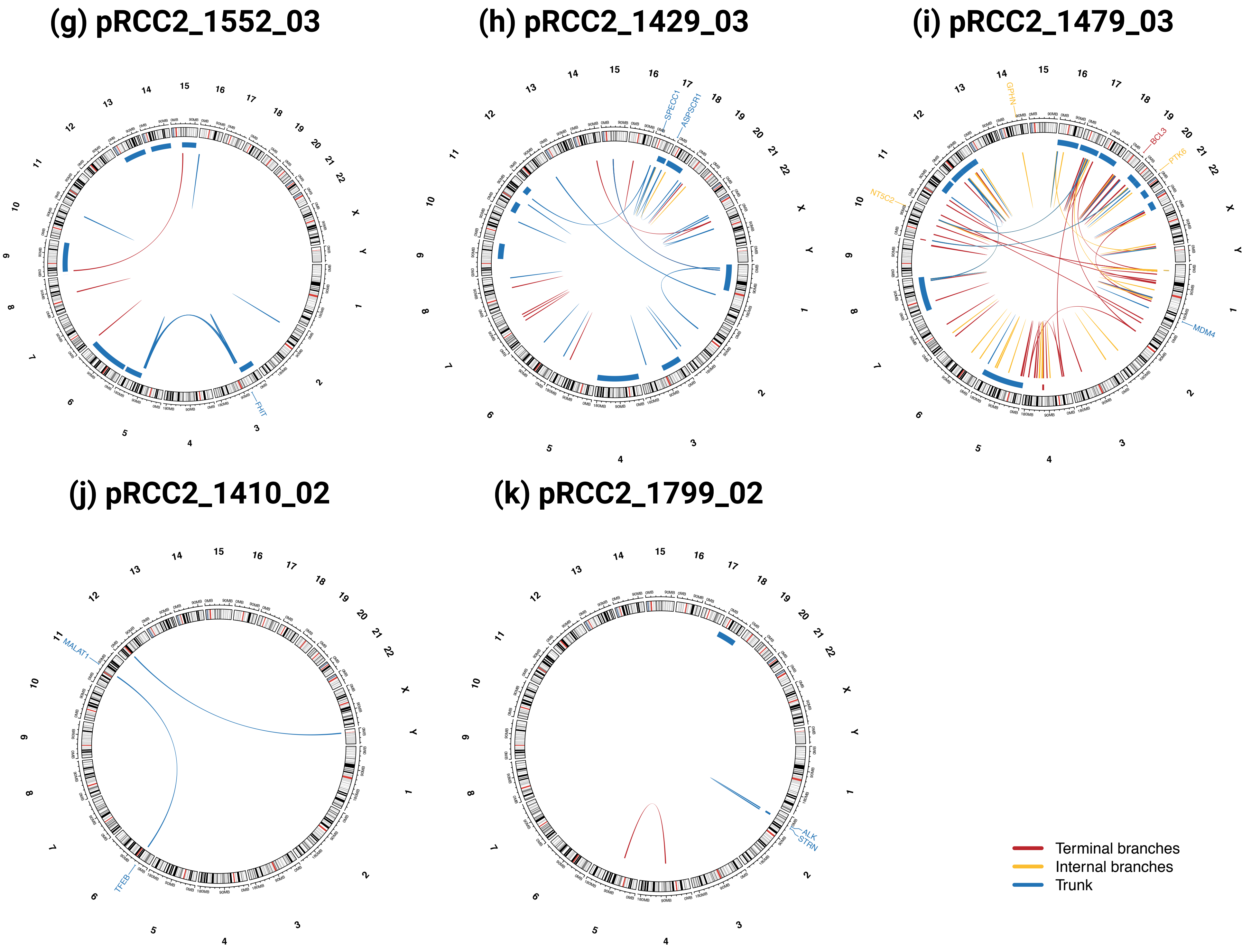

(g) pRCC2_1552_03
FHIT
(h) pRCC2_1429_03
SPECC1
ASPSCR1
(i) pRCC2_1479_03
GPHN
BCL3
PTK6
NT5C2
MDM4
(j) pRCC2_1410_02
MALAT1
TFEB
(k) pRCC2_1799_02
ALK
STRN
Terminal branches
Internal branches
Trunk

### (l) rSRC_1697_10

### (m) cdRCC_1929_03

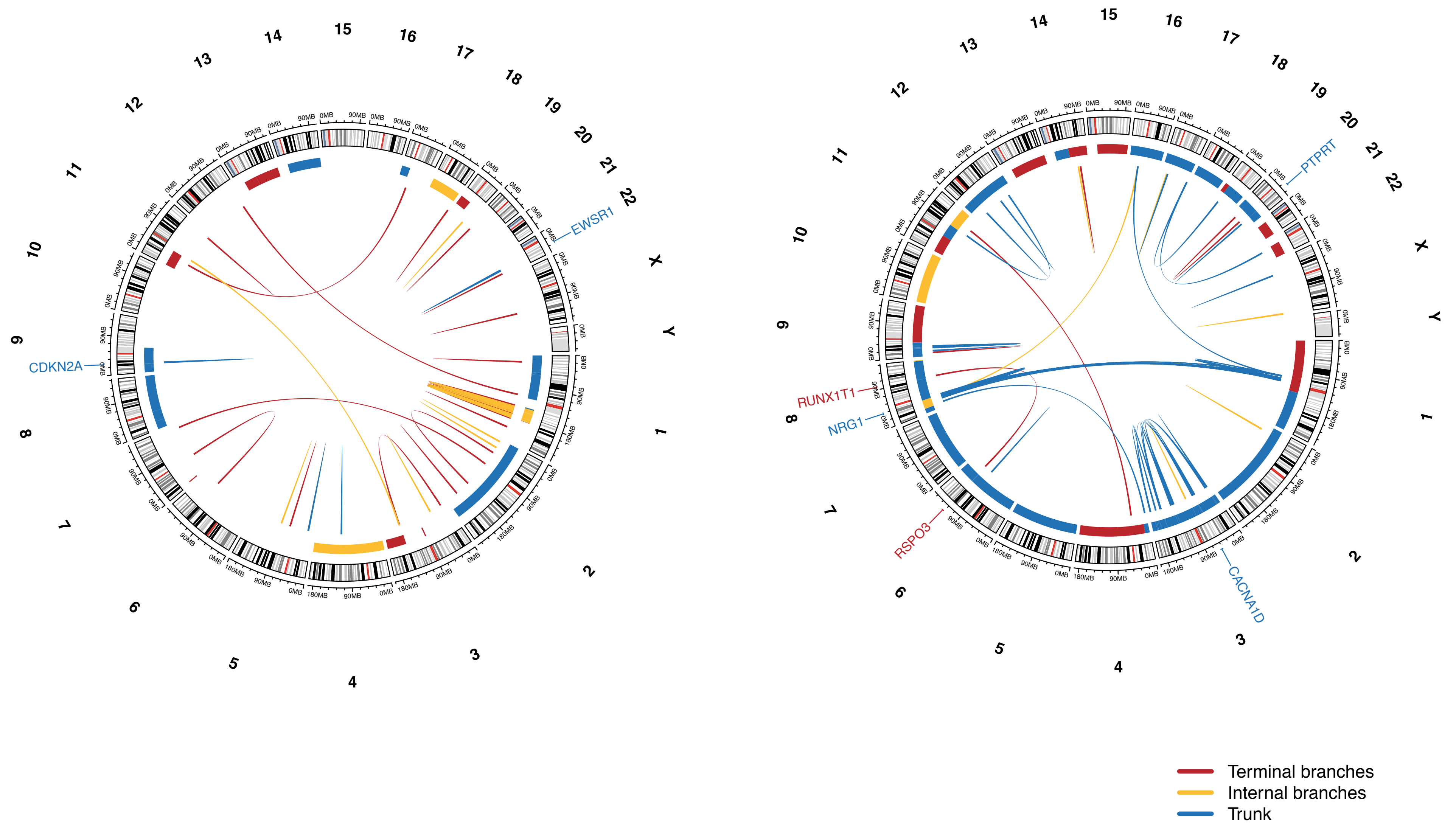

**Supplementary Figure 16: Validation of the WGS-detected SV events.** A PCR-based sequencing methodology was used (AmpliSeq) to validate the whole-genome sequencing based SV events. Read counts were colored based on their magnitude. Bottom blue bar indicates SVs shared by all regions; orange bar SVs shared by part of regions; and red bar SVs found in one region only.

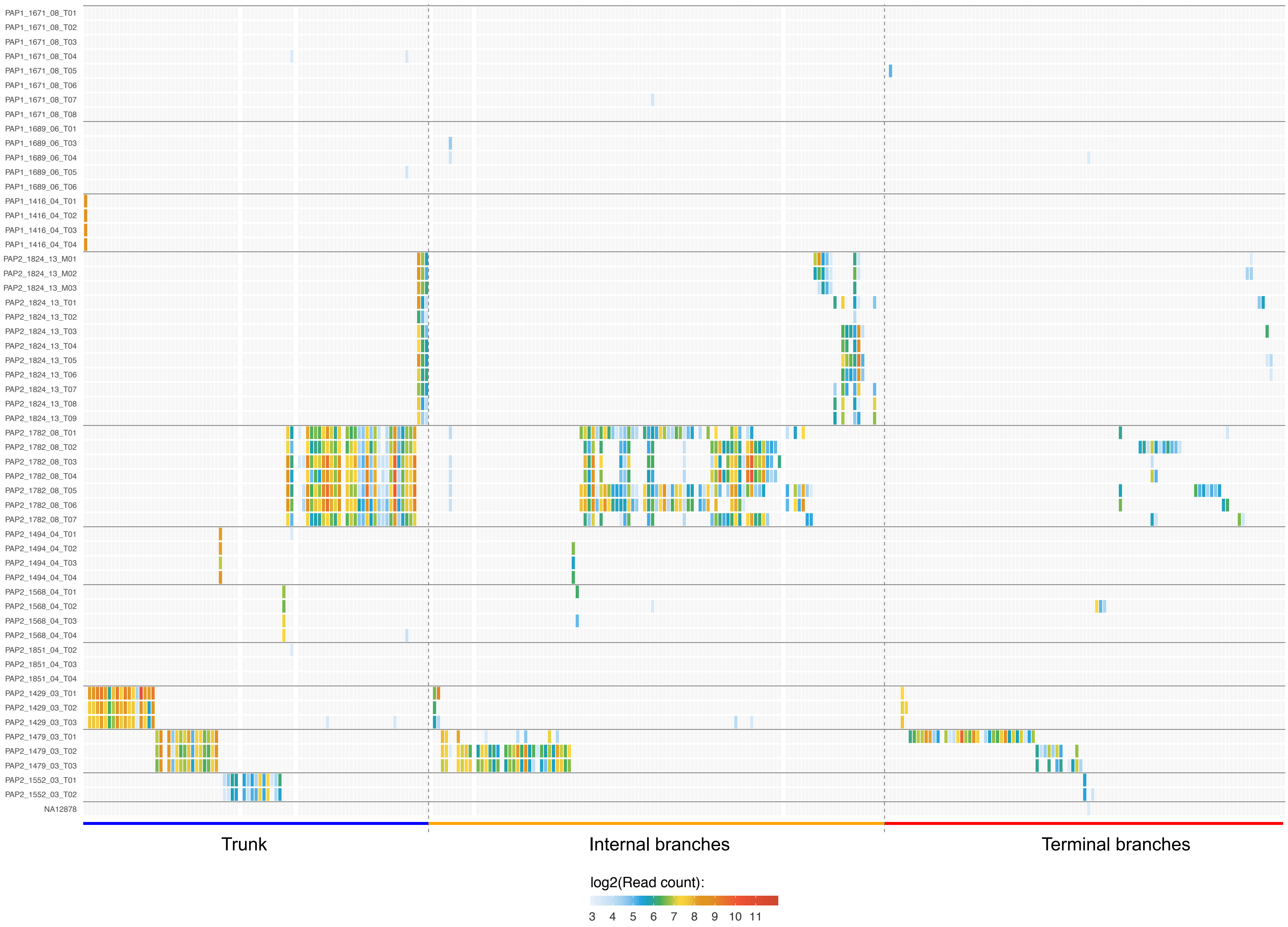

PAP1_1671_08_T01
PAP1_1671_08_T02
PAP1_1671_08_T03
PAP1_1671_08_T04
PAP1_1671_08_T05
PAP1_1671_08_T06
PAP1_1671_08_T07
PAP1_1671_08_T08
PAP1_1689_06_T01
PAP1_1689_06_T03
PAP1_1689_06_T04
PAP1_1689_06_T05
PAP1_1689_06_T06
PAP1_1416_04_T01
PAP1_1416_04_T02
PAP1_1416_04_T03
PAP1_1416_04_T04
PAP2_1824_13_M01
PAP2_1824_13_M02
PAP2_1824_13_M03
PAP2_1824_13_T01
PAP2_1824_13_T02
PAP2_1824_13_T03
PAP2_1824_13_T04
PAP2_1824_13_T05
PAP2_1824_13_T06
PAP2_1824_13_T07
PAP2_1824_13_T08
PAP2_1824_13_T09
PAP2_1782_08_T01
PAP2_1782_08_T02
PAP2_1782_08_T03
PAP2_1782_08_T04
PAP2_1782_08_T05
PAP2_1782_08_T06
PAP2_1782_08_T07
PAP2_1494_04_T01
PAP2_1494_04_T02
PAP2_1494_04_T03
PAP2_1494_04_T04
PAP2_1568_04_T01
PAP2_1568_04_T02
PAP2_1568_04_T03
PAP2_1568_04_T04
PAP2_1851_04_T02
PAP2_1851_04_T03
PAP2_1851_04_T04
PAP2_1429_03_T01
PAP2_1429_03_T02
PAP2_1429_03_T03
PAP2_1479_03_T01
PAP2_1479_03_T02
PAP2_1479_03_T03
PAP2_1552_03_T01
PAP2_1552_03_T02
NA12878
Trunk
Internal branches
Terminal branches
log2(Read count):
3 4 5 6 7 8 9 10 11

**Supplementary Figure 17: Number of SVs with nearby breakpoints in copy number, separated by type.** Unobserved SVs have no copy number breakpoint within 300KB, subclonal SVs have a copy number breakpoint in a subset of copy number calls within 300KB and clonal SVs have a copy number breakpoint within 300KB in all samples from the same tumor. Transloc = translocation, tandem = tandem duplication, invers = inversion, ins = insertion, del = deletion.

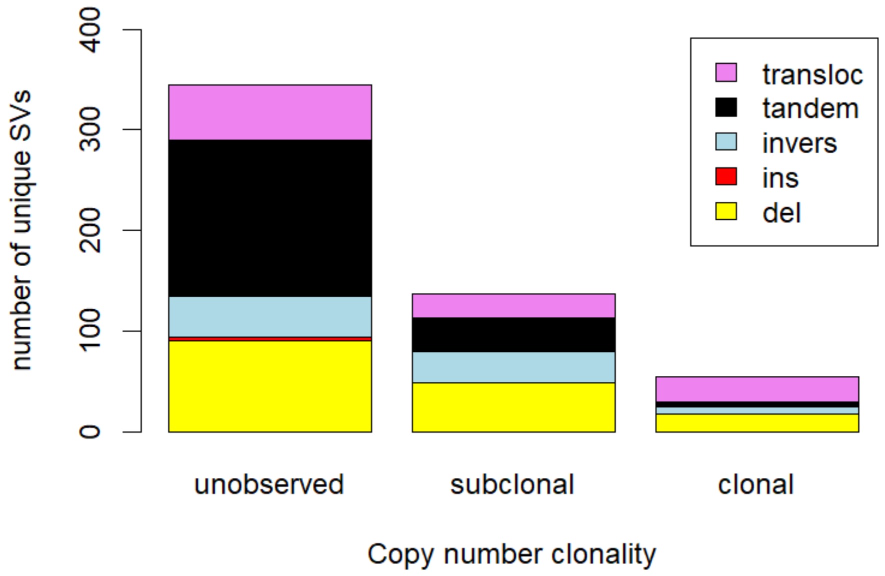

400
300
200
100
0
number of unique SVs
transloc
tandem
invers
ins
del
unobserved
subclonal
clonal
Copy number clonality

**Supplementary Figure 18: The four SNV mutational signatures identified by de novo extraction.** The mutational signature is displayed by 96 mutation types, defined by the mutated base and its sequence context immediately 3′ and 5′ on the horizontal axes. The vertical axes depict the percentage of mutations attributed to each mutation type

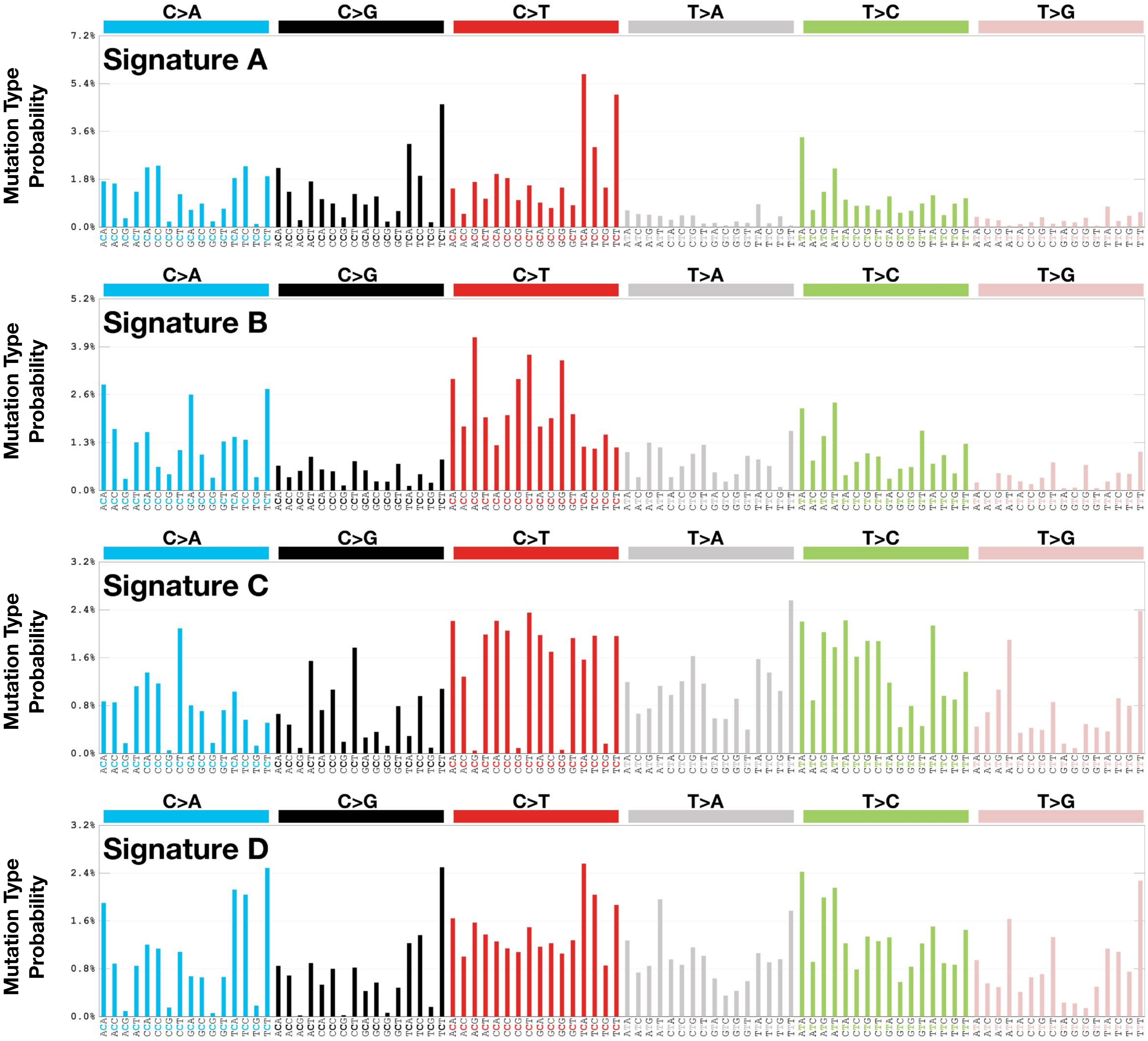
C>A
C>G
C>T
T>A
T>C
T>G
Signature A
Mutation Type Probability
7.2%
5.4%
3.6%
1.8%
0.0%
C>A
C>G
C>T
T>A
T>C
T>G
Signature B
Mutation Type Probability
5.2%
3.9%
2.6%
1.3%
0.0%
C>A
C>G
C>T
T>A
T>C
T>G
Signature C
Mutation Type Probability
3.2%
2.4%
1.6%
0.8%
0.0%
C>A
C>G
C>T
T>A
T>C
T>G
Signature D
Mutation Type Probability
3.2%
2.4%
1.6%
0.8%
0.0%

**Supplementary Figure 19:** Scatter plots of number of (a) clonal or (b) subclonal mutations assigned to signatures 1, 5, and 40 with age at diagnosis.

**a**

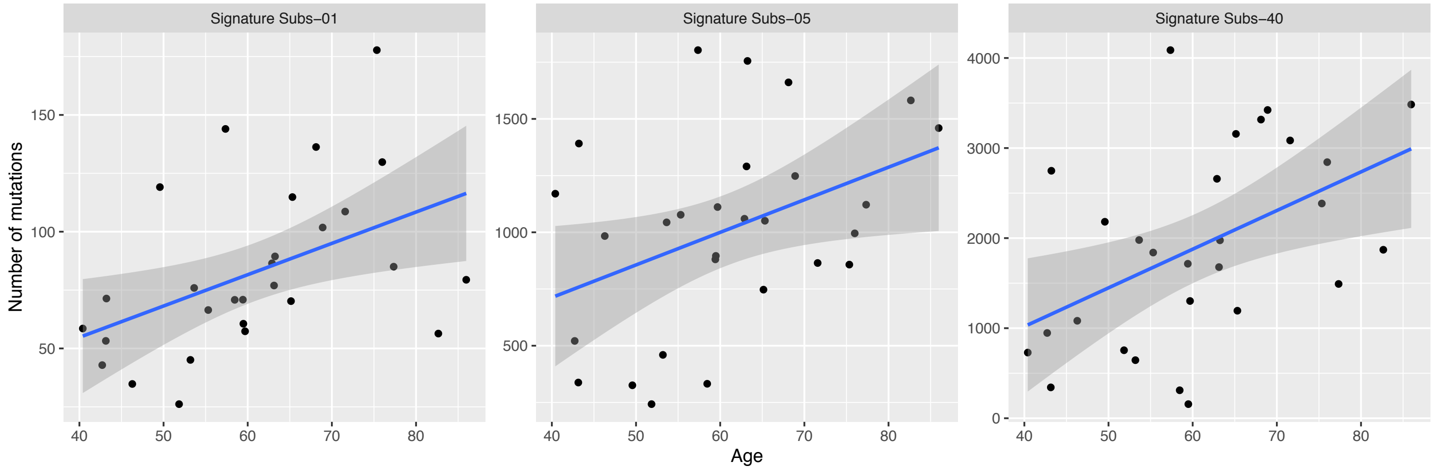

Signature Subs−01
Signature Subs−05
Signature Subs−40
Number of mutations
Age
150
100
50
1500
1000
500
4000
3000
2000
1000
0
40
50
60
70
80

**b**

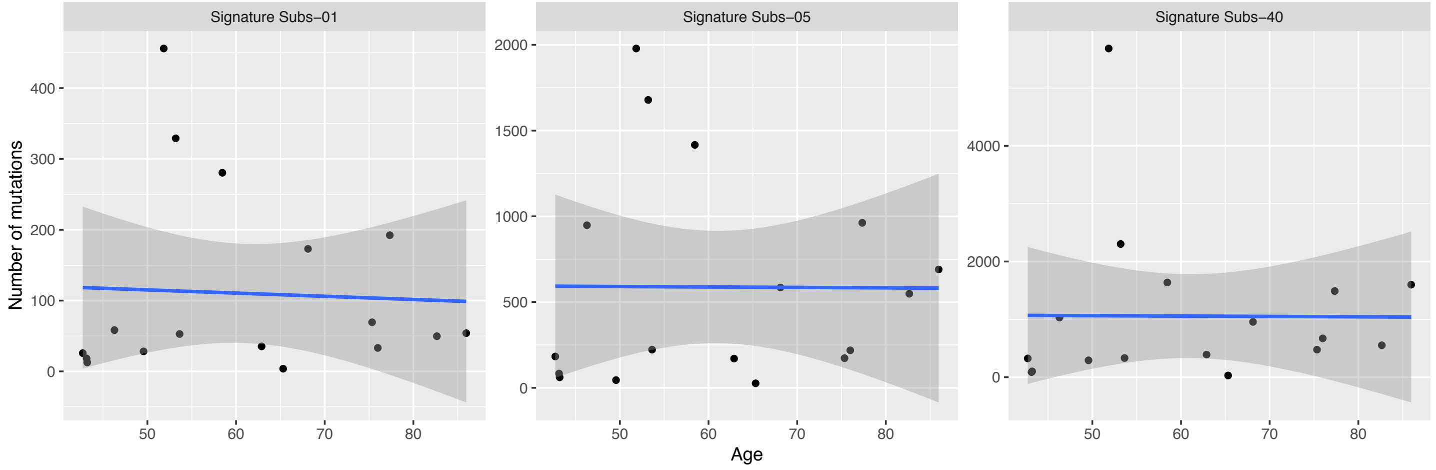

Signature Subs−01
Signature Subs−05
Signature Subs−40
Number of mutations
Age
400
300
200
100
0
2000
1500
1000
500
0
4000
2000
0
50
60
70
80

**Supplementary Figure 20: Proportions of mutational signatures.** The contribution of known mutational signatures in each sample.

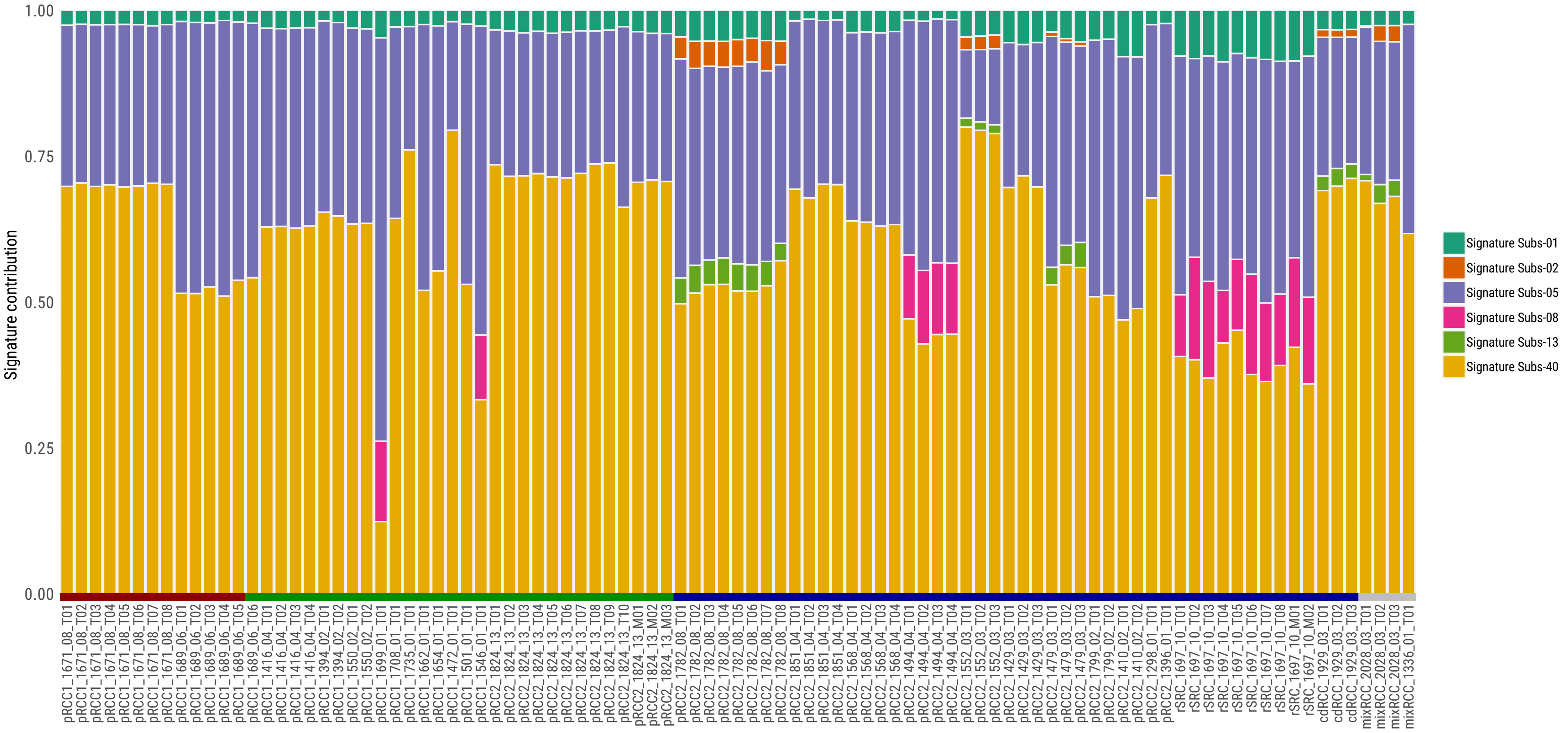

Signature contribution
1.00
0.75
0.50
0.25
0.00
Signature Subs-01
Signature Subs-02
Signature Subs-05
Signature Subs-08
Signature Subs-13
Signature Subs-40
pRCC1_1671_08_T01
pRCC1_1671_08_T02
pRCC1_1671_08_T03
pRCC1_1671_08_T04
pRCC1_1671_08_T05
pRCC1_1671_08_T06
pRCC1_1671_08_T07
pRCC1_1671_08_T08
pRCC1_1689_06_T01
pRCC1_1689_06_T02
pRCC1_1689_06_T03
pRCC1_1689_06_T04
pRCC1_1689_06_T05
pRCC1_1689_06_T06
pRCC1_1416_04_T01
pRCC1_1416_04_T02
pRCC1_1416_04_T03
pRCC1_1416_04_T04
pRCC1_1394_02_T01
pRCC1_1394_02_T02
pRCC1_1550_02_T01
pRCC1_1550_02_T02
pRCC1_1699_01_T01
pRCC1_1708_01_T01
pRCC1_1735_01_T01
pRCC1_1662_01_T01
pRCC1_1654_01_T01
pRCC1_1472_01_T01
pRCC1_1501_01_T01
pRCC1_1546_01_T01
pRCC2_1824_13_T01
pRCC2_1824_13_T02
pRCC2_1824_13_T03
pRCC2_1824_13_T04
pRCC2_1824_13_T05
pRCC2_1824_13_T06
pRCC2_1824_13_T07
pRCC2_1824_13_T08
pRCC2_1824_13_T09
pRCC2_1824_13_T10
pRCC2_1824_13_M01
pRCC2_1824_13_M02
pRCC2_1824_13_M03
pRCC2_1782_08_T01
pRCC2_1782_08_T02
pRCC2_1782_08_T03
pRCC2_1782_08_T04
pRCC2_1782_08_T05
pRCC2_1782_08_T06
pRCC2_1782_08_T07
pRCC2_1782_08_T08
pRCC2_1851_04_T01
pRCC2_1851_04_T02
pRCC2_1851_04_T03
pRCC2_1851_04_T04
pRCC2_1568_04_T01
pRCC2_1568_04_T02
pRCC2_1568_04_T03
pRCC2_1568_04_T04
pRCC2_1494_04_T01
pRCC2_1494_04_T02
pRCC2_1494_04_T03
pRCC2_1494_04_T04
pRCC2_1552_03_T01
pRCC2_1552_03_T02
pRCC2_1552_03_T03
pRCC2_1429_03_T01
pRCC2_1429_03_T02
pRCC2_1429_03_T03
pRCC2_1479_03_T01
pRCC2_1479_03_T02
pRCC2_1479_03_T03
pRCC2_1799_02_T01
pRCC2_1799_02_T02
pRCC2_1410_02_T01
pRCC2_1410_02_T02
pRCC2_1298_01_T01
pRCC2_1396_01_T01
rSRC_1697_10_T01
rSRC_1697_10_T02
rSRC_1697_10_T03
rSRC_1697_10_T04
rSRC_1697_10_T05
rSRC_1697_10_T06
rSRC_1697_10_T07
rSRC_1697_10_T08
rSRC_1697_10_M01
rSRC_1697_10_M02
cdRCC_1929_03_T01
cdRCC_1929_03_T02
cdRCC_1929_03_T03
mixRCC_2028_03_T01
mixRCC_2028_03_T02
mixRCC_2028_03_T03
mixRCC_1336_01_T01

**Supplementary Figure 21: Clonality of known mutational signatures for all samples and by subtypes.** The proportion of mutational signatures are reported for trunks and branches.

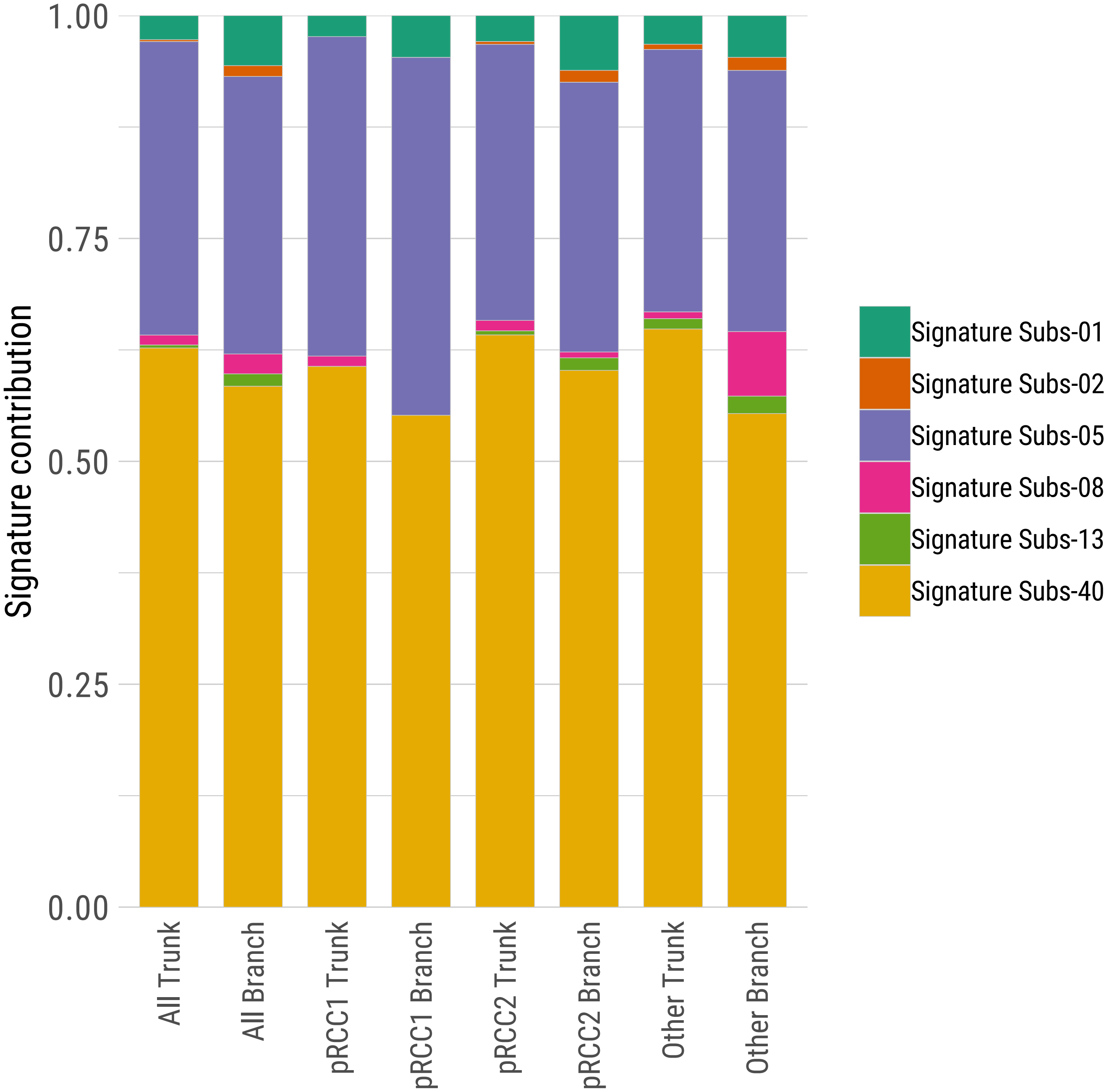

1.00
0.75
0.50
0.25
0.00
Signature contribution
All Trunk
All Branch
pRCC1 Trunk
pRCC1 Branch
pRCC2 Trunk
pRCC2 Branch
Other Trunk
Other Branch
Signature Subs-01
Signature Subs-02
Signature Subs-05
Signature Subs-08
Signature Subs-13
Signature Subs-40

**Supplementary Figure 22: Mutational signatures for primary and metastatic tumors.** The contribution of known mutational signatures in tumor pRCC2_1824_13 and rSRC_1697_10.

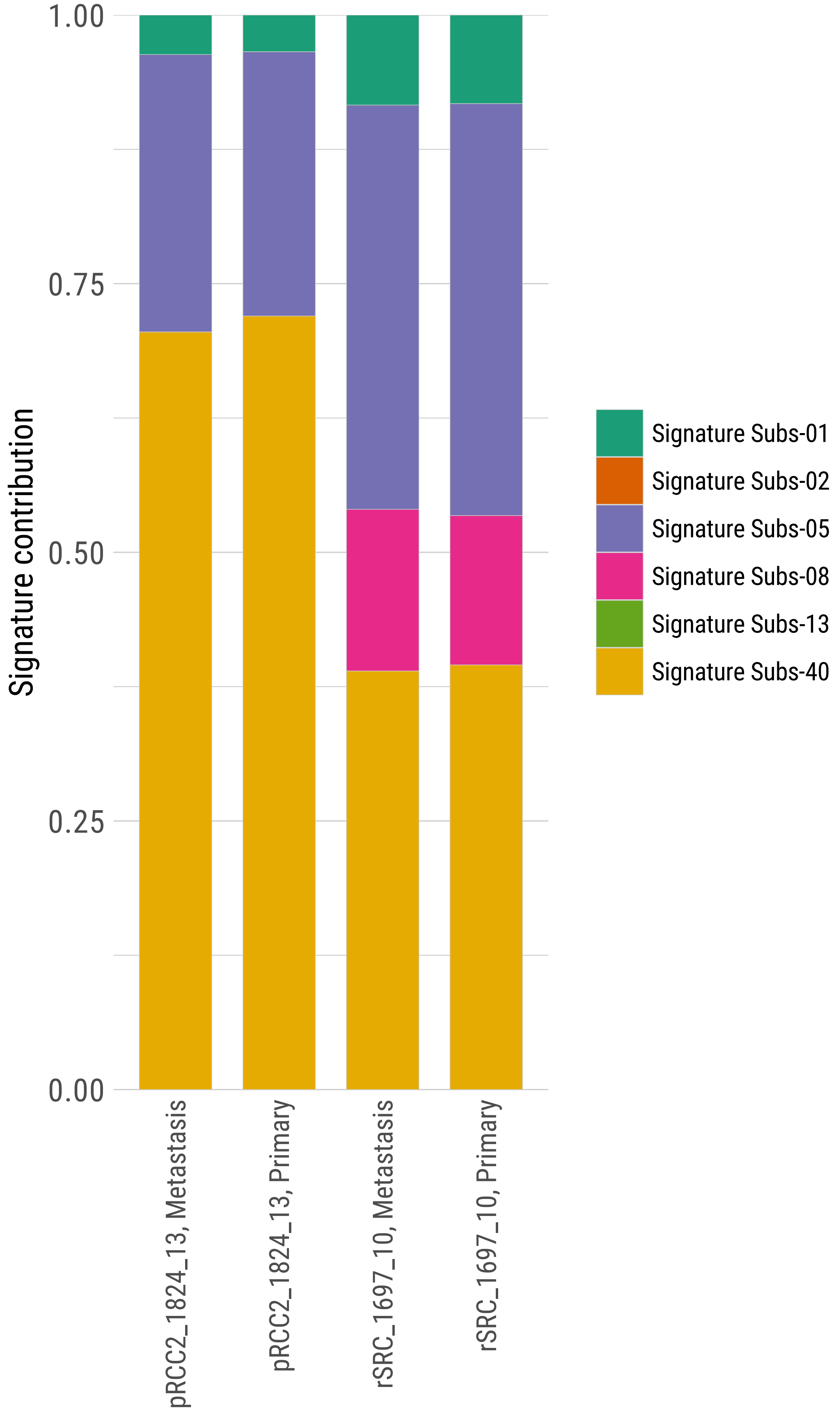
Signature contribution
1.00
0.75
0.50
0.25
0.00
pRCC2_1824_13, Metastasis
pRCC2_1824_13, Primary
rSRC_1697_10, Metastasis
rSRC_1697_10, Primary
Signature Subs-01
Signature Subs-02
Signature Subs-05
Signature Subs-08
Signature Subs-13
Signature Subs-40

**Supplementary Figure 23: Telomere length (TL) for each sample.** TL is based on the abundance of telomere motif sequence (TTAGGG/CCCTAA)4.

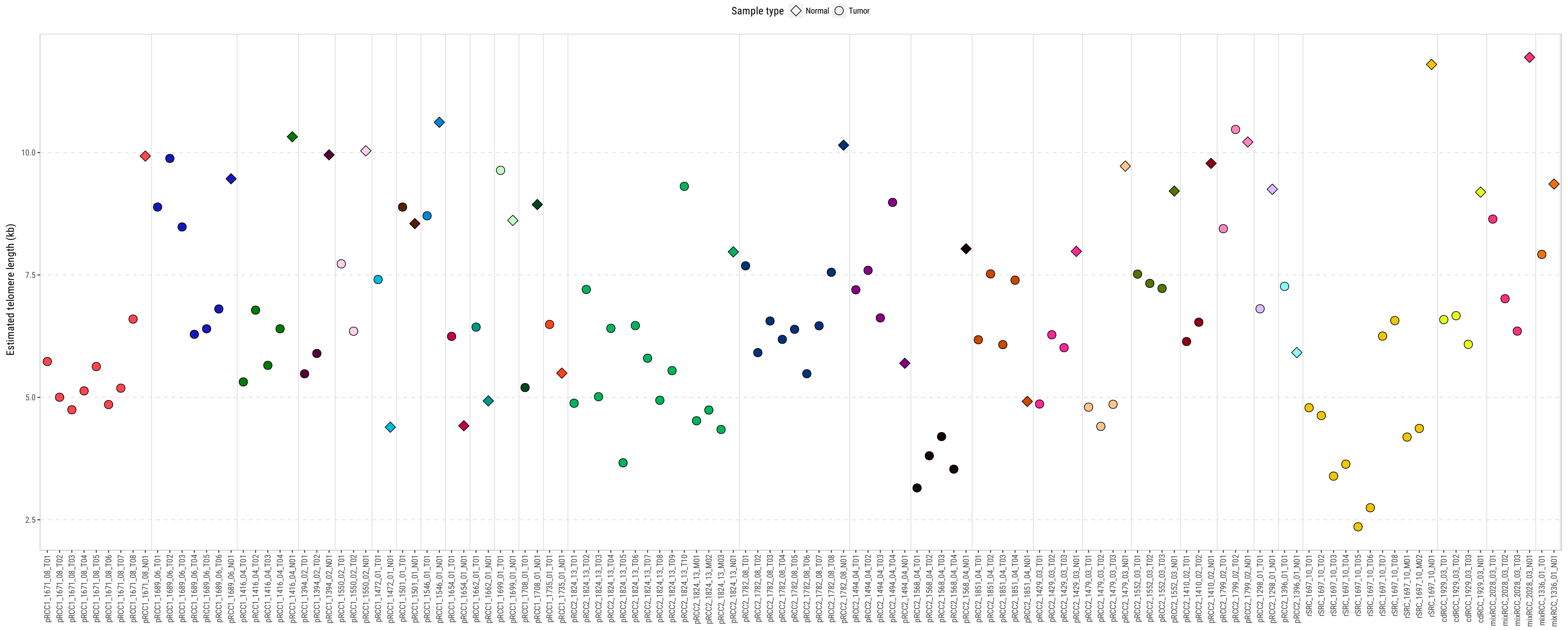
Sample type
Normal
Tumor
Estimated telomere length (kb)
10.0
7.5
5.0
2.5
pRCC1_1671_08_T01
pRCC1_1671_08_T02
pRCC1_1671_08_T03
pRCC1_1671_08_T04
pRCC1_1671_08_T05
pRCC1_1671_08_T06
pRCC1_1671_08_T07
pRCC1_1671_08_T08
pRCC1_1671_08_N01
pRCC1_1689_06_T01
pRCC1_1689_06_T02
pRCC1_1689_06_T03
pRCC1_1689_06_T04
pRCC1_1689_06_T05
pRCC1_1689_06_T06
pRCC1_1689_06_N01
pRCC1_1416_04_T01
pRCC1_1416_04_T02
pRCC1_1416_04_T03
pRCC1_1416_04_T04
pRCC1_1416_04_N01
pRCC1_1394_02_T01
pRCC1_1394_02_T02
pRCC1_1394_02_N01
pRCC1_1550_02_T01
pRCC1_1550_02_T02
pRCC1_1550_02_N01
pRCC1_1472_01_T01
pRCC1_1472_01_N01
pRCC1_1501_01_T01
pRCC1_1501_01_N01
pRCC1_1546_01_T01
pRCC1_1546_01_N01
pRCC1_1654_01_T01
pRCC1_1654_01_N01
pRCC1_1662_01_T01
pRCC1_1662_01_N01
pRCC1_1699_01_T01
pRCC1_1699_01_N01
pRCC1_1708_01_T01
pRCC1_1708_01_N01
pRCC1_1735_01_T01
pRCC1_1735_01_N01
pRCC2_1824_13_T01
pRCC2_1824_13_T02
pRCC2_1824_13_T03
pRCC2_1824_13_T04
pRCC2_1824_13_T05
pRCC2_1824_13_T06
pRCC2_1824_13_T07
pRCC2_1824_13_T08
pRCC2_1824_13_T09
pRCC2_1824_13_T10
pRCC2_1824_13_M01
pRCC2_1824_13_M02
pRCC2_1824_13_M03
pRCC2_1824_13_N01
pRCC2_1782_08_T01
pRCC2_1782_08_T02
pRCC2_1782_08_T03
pRCC2_1782_08_T04
pRCC2_1782_08_T05
pRCC2_1782_08_T06
pRCC2_1782_08_T07
pRCC2_1782_08_T08
pRCC2_1782_08_N01
pRCC2_1494_04_T01
pRCC2_1494_04_T02
pRCC2_1494_04_T03
pRCC2_1494_04_T04
pRCC2_1494_04_N01
pRCC2_1568_04_T01
pRCC2_1568_04_T02
pRCC2_1568_04_T03
pRCC2_1568_04_T04
pRCC2_1568_04_N01
pRCC2_1851_04_T01
pRCC2_1851_04_T02
pRCC2_1851_04_T03
pRCC2_1851_04_T04
pRCC2_1851_04_N01
pRCC2_1429_03_T01
pRCC2_1429_03_T02
pRCC2_1429_03_T03
pRCC2_1429_03_N01
pRCC2_1479_03_T01
pRCC2_1479_03_T02
pRCC2_1479_03_T03
pRCC2_1479_03_N01
pRCC2_1552_03_T01
pRCC2_1552_03_T02
pRCC2_1552_03_T03
pRCC2_1552_03_N01
pRCC2_1410_02_T01
pRCC2_1410_02_T02
pRCC2_1410_02_N01
pRCC2_1799_02_T01
pRCC2_1799_02_T02
pRCC2_1799_02_N01
pRCC2_1298_01_T01
pRCC2_1298_01_N01
pRCC2_1396_01_T01
pRCC2_1396_01_N01
rSRC_1697_10_T01
rSRC_1697_10_T02
rSRC_1697_10_T03
rSRC_1697_10_T04
rSRC_1697_10_T05
rSRC_1697_10_T06
rSRC_1697_10_T07
rSRC_1697_10_T08
rSRC_1697_10_M01
rSRC_1697_10_M02
rSRC_1697_10_N01
cdRCC_1929_03_T01
cdRCC_1929_03_T02
cdRCC_1929_03_T03
cdRCC_1929_03_N01
mixRCC_2028_03_T01
mixRCC_2028_03_T02
mixRCC_2028_03_T03
mixRCC_2028_03_N01
mixRCC_1336_01_T01
mixRCC_1336_01_N01

**Supplementary Figure 24: Clonality of methylation status**. The proportion of methylation levels based on the multi-regional trees (MRT) in trunks, internal branches, and terminal branches for each tumor.

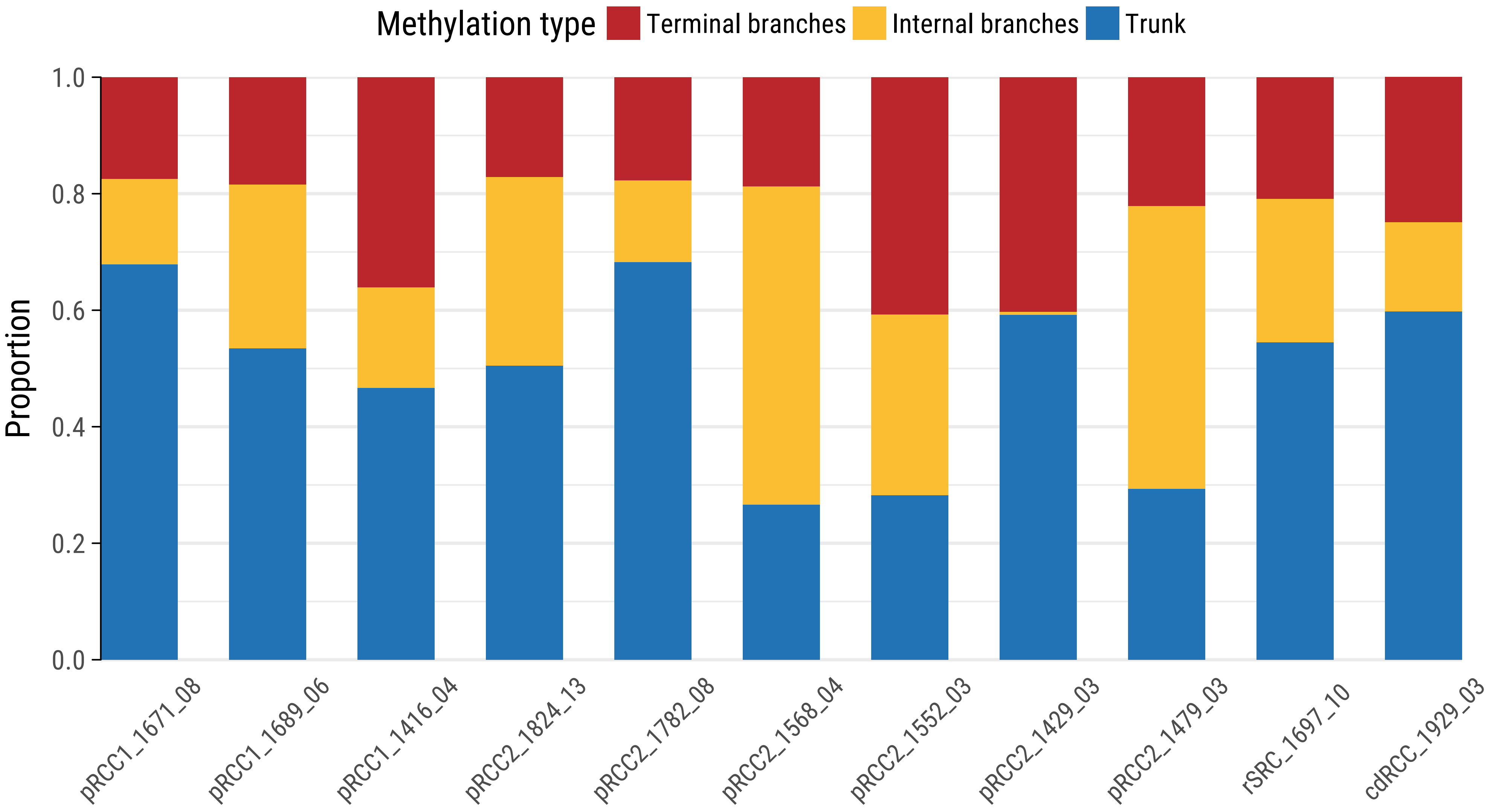
Methylation type
Terminal branches
Internal branches
Trunk
Proportion
1.0
0.8
0.6
0.4
0.2
0.0
pRCC1_1671_08
pRCC1_1689_06
pRCC1_1416_04
pRCC2_1824_13
pRCC2_1782_08
pRCC2_1568_04
pRCC2_1552_03
pRCC2_1429_03
pRCC2_1479_03
rSRC_1697_10
cdRCC_1929_03

**Supplementary Figure 25: Diagnostic close-up images of H&E stained slides of the 39 treatment-naive renal cell carcinomas studied here.** They include: 23 papillary type 1 (pRCC1); 12 papillary type 2 (pRCC2); one collecting duct carcinoma (cdRCC); one renal fibrosarcoma (rSRC); one mixed pRCC1/pRCC2 and one unclassified renal cancer with mixed features of pRCC2 and cdRCC. For the tumors with mixed histological components (mixRCC_1336_01 and mixRCC_2028_03), close-ups of both tumor components are shown.

cdRCC_1929_03

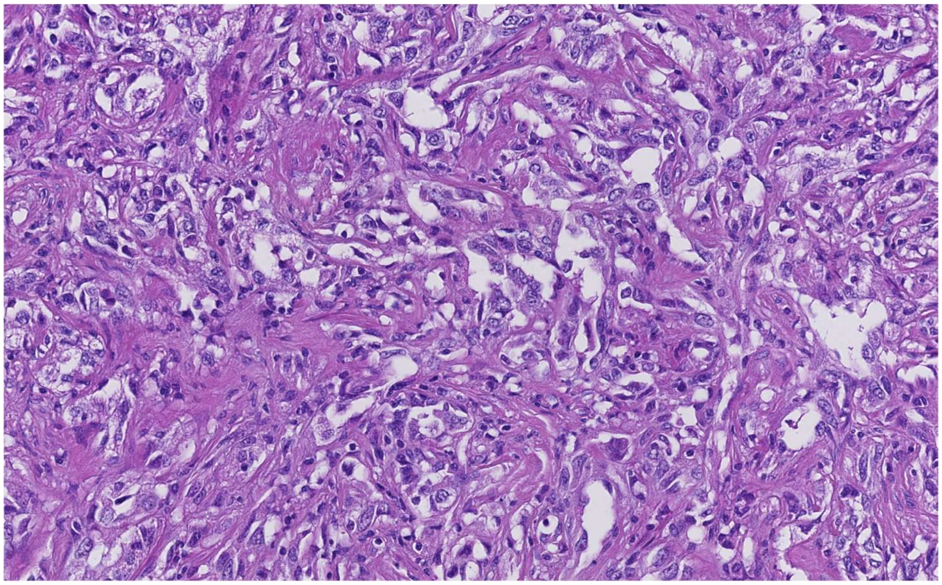

mixRCC_1336_01(1)

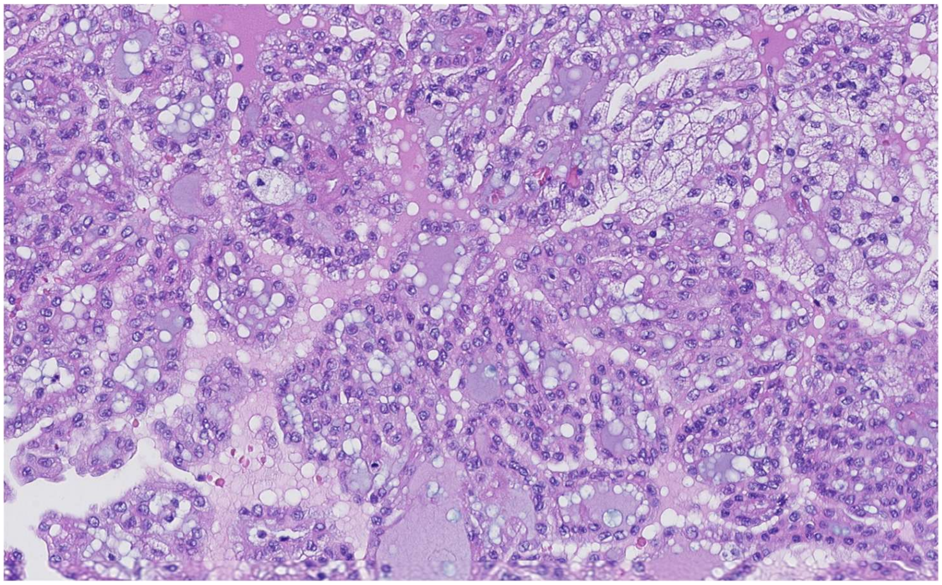

mixRCC_1336_01

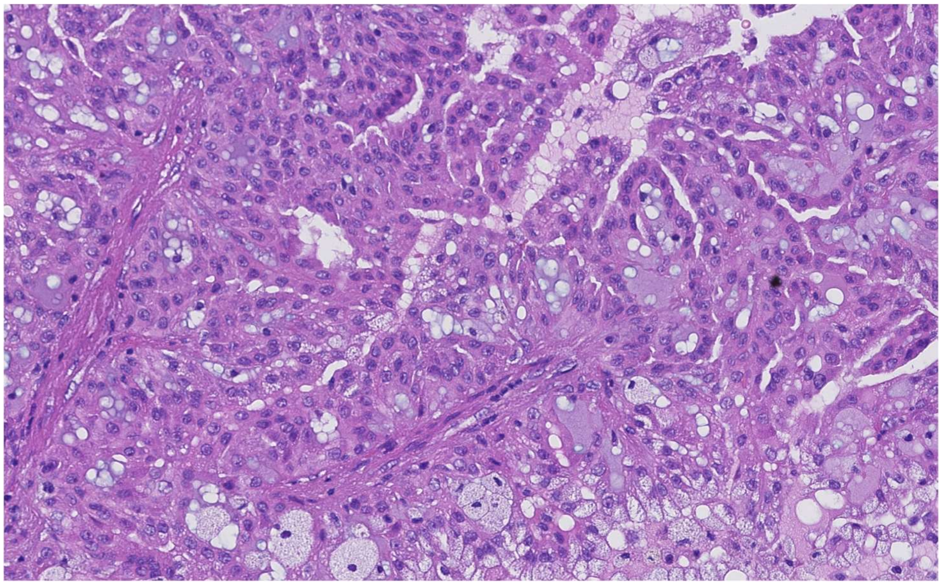

mixRCC_2028_03(1)

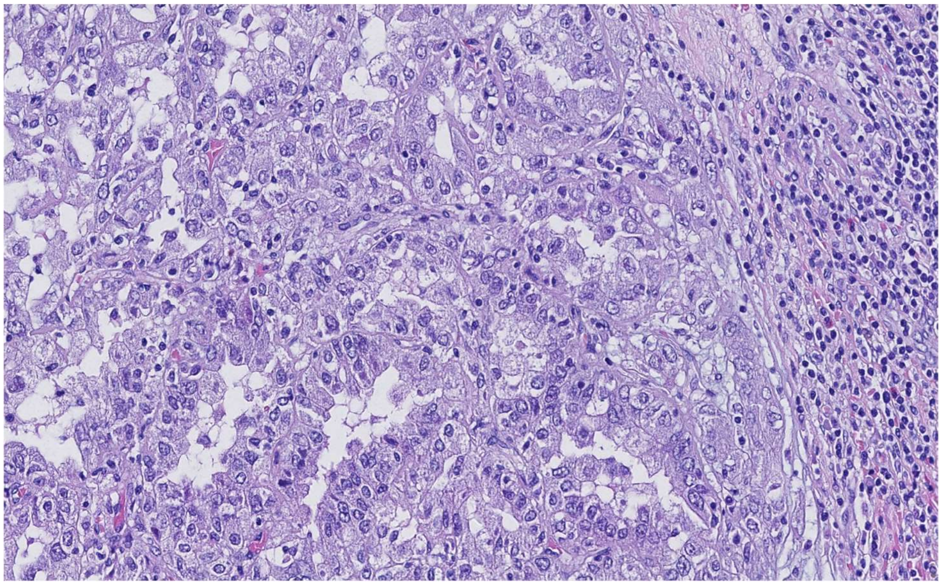

mixRCC_2028_03

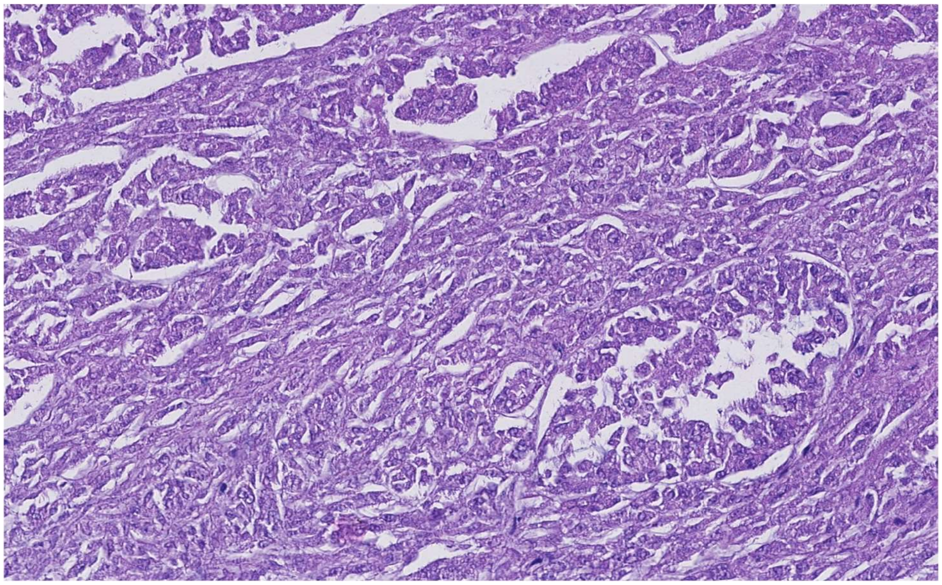

pRCC1_1394_02

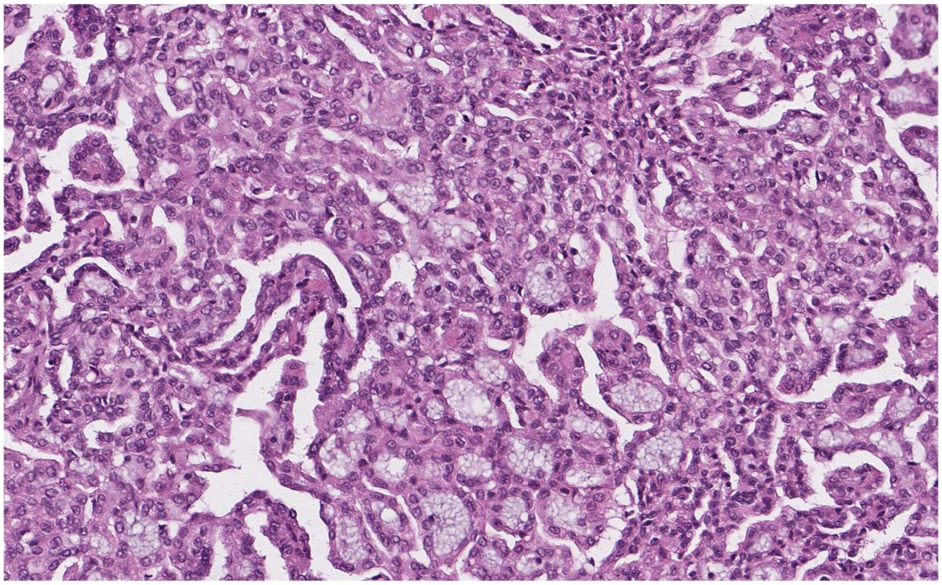

pRCC1_1401_01

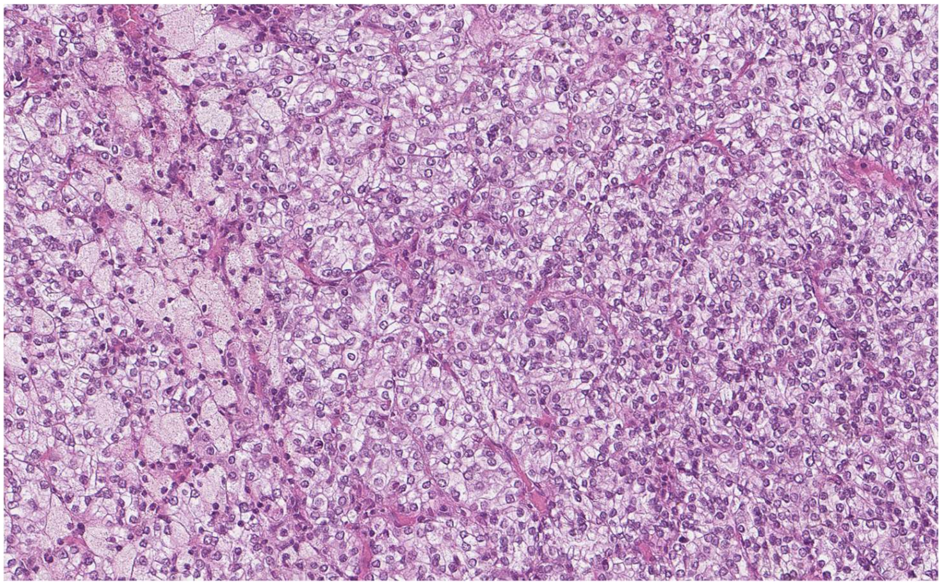

pRCC1_1402_01

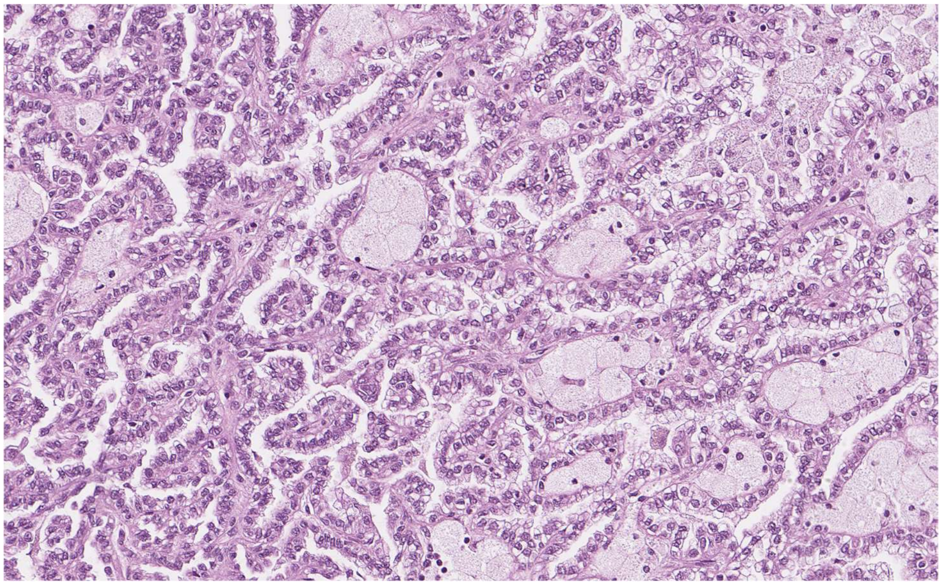

pRCC1_1405_01

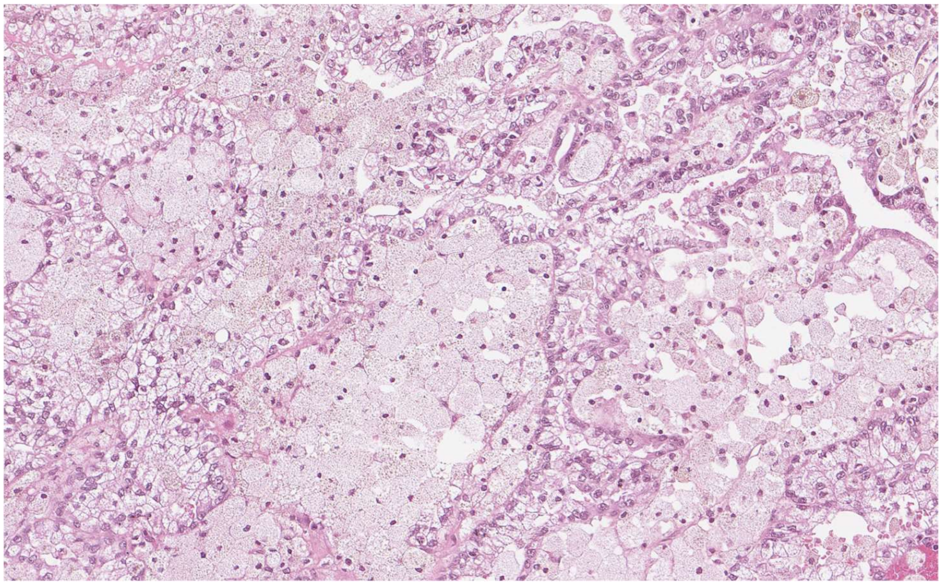

pRCC1_1416_04

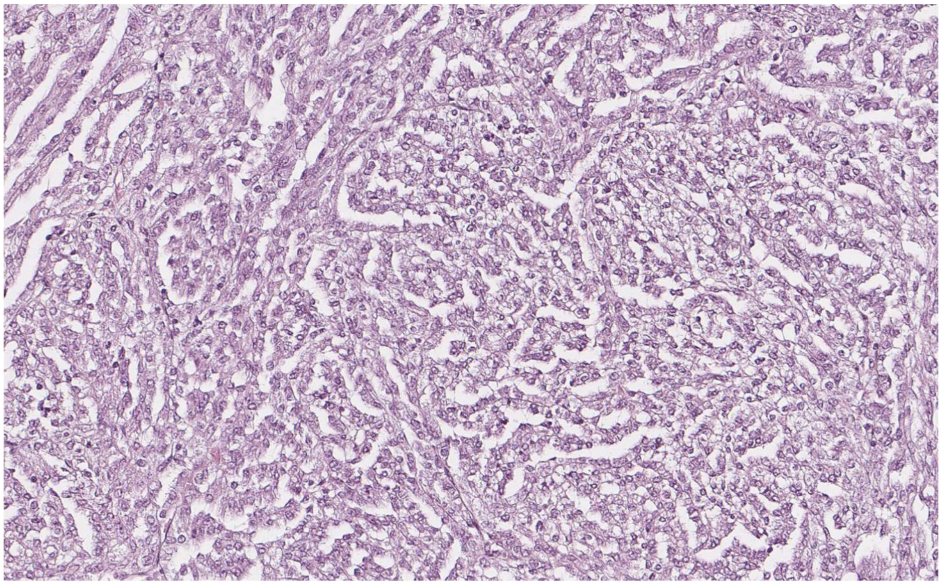

pRCC1_1418_01

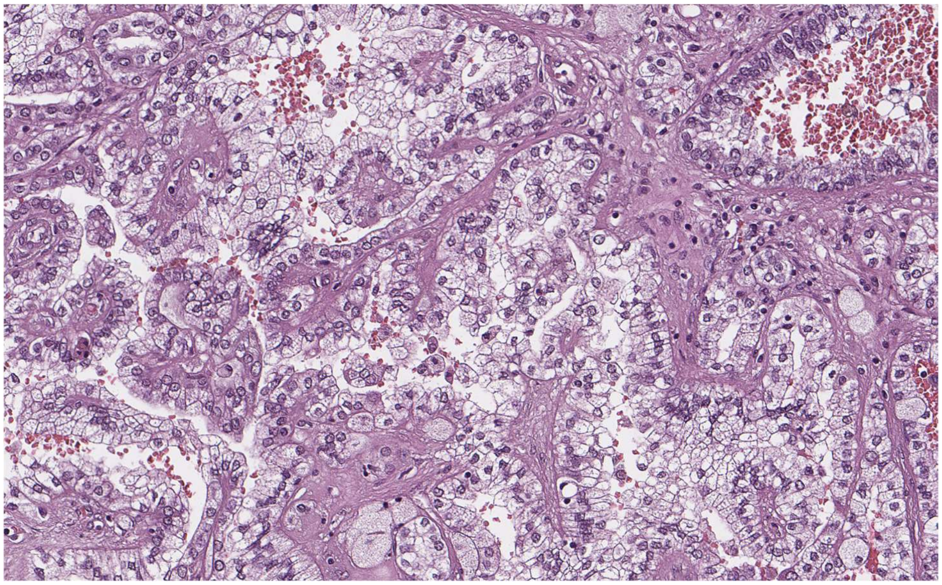

pRCC1_1426_01

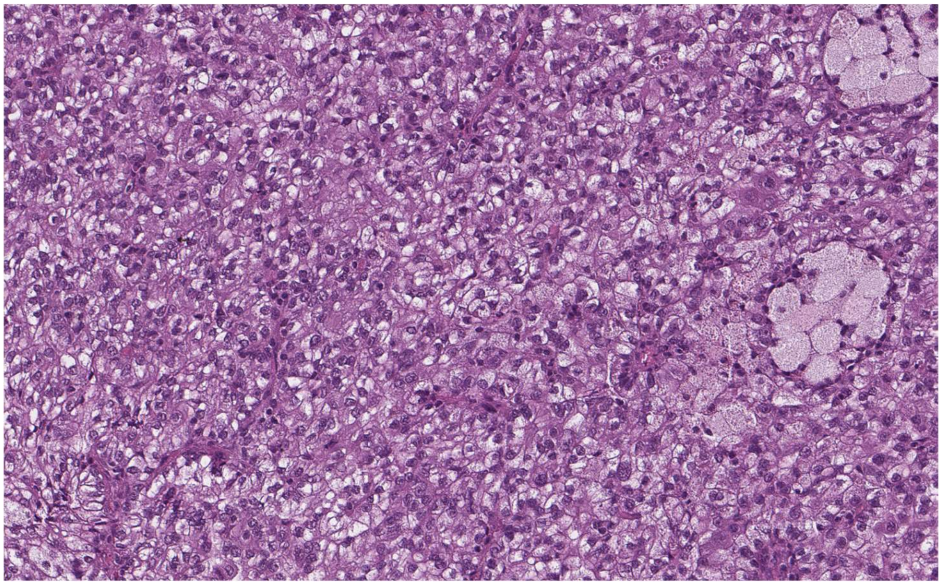

pRCC1_1434_01

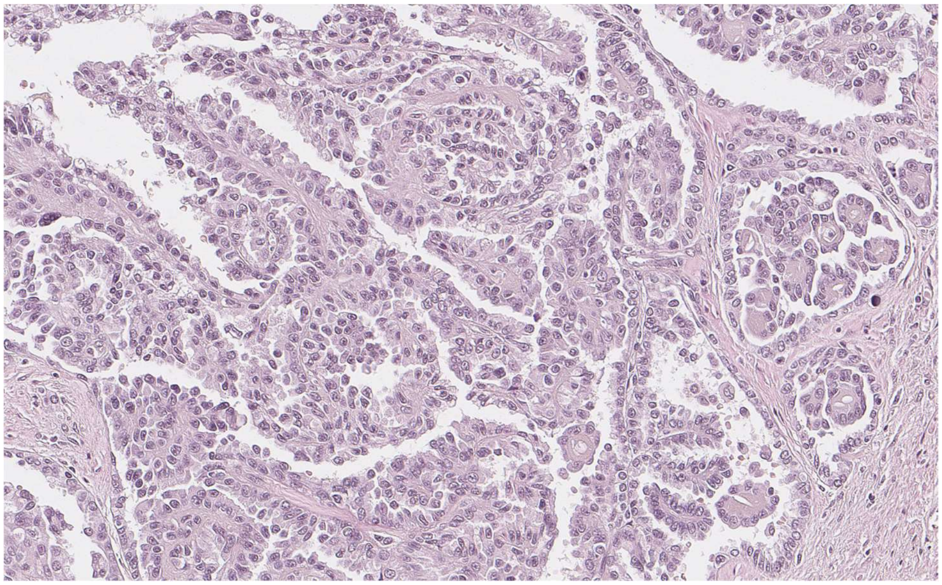

pRCC1_1459_01

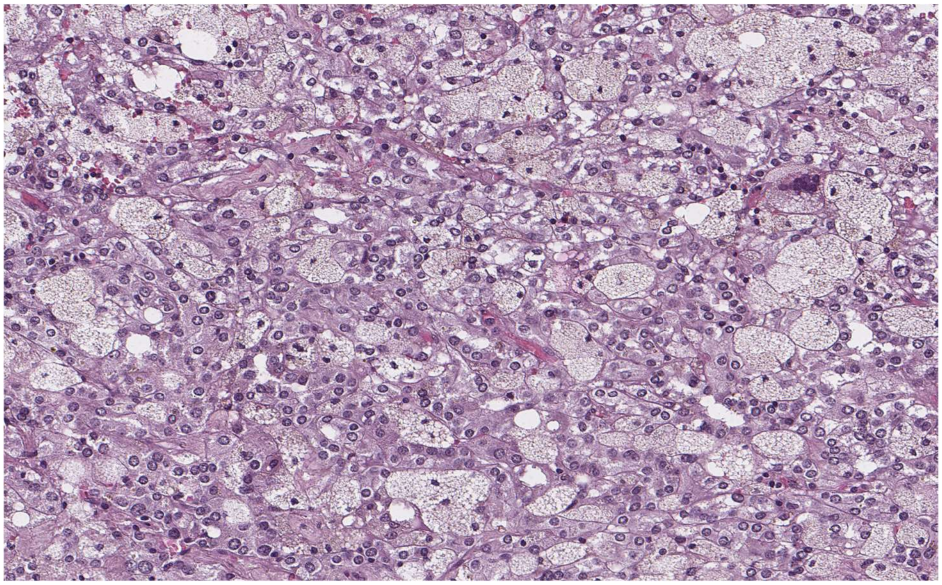

pRCC1_1464_01

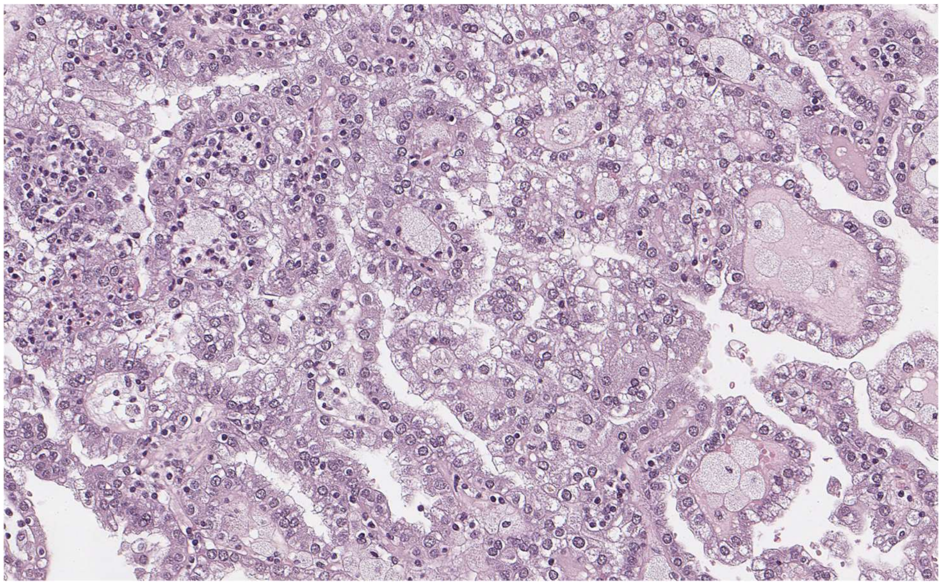

pRCC1_1472_01

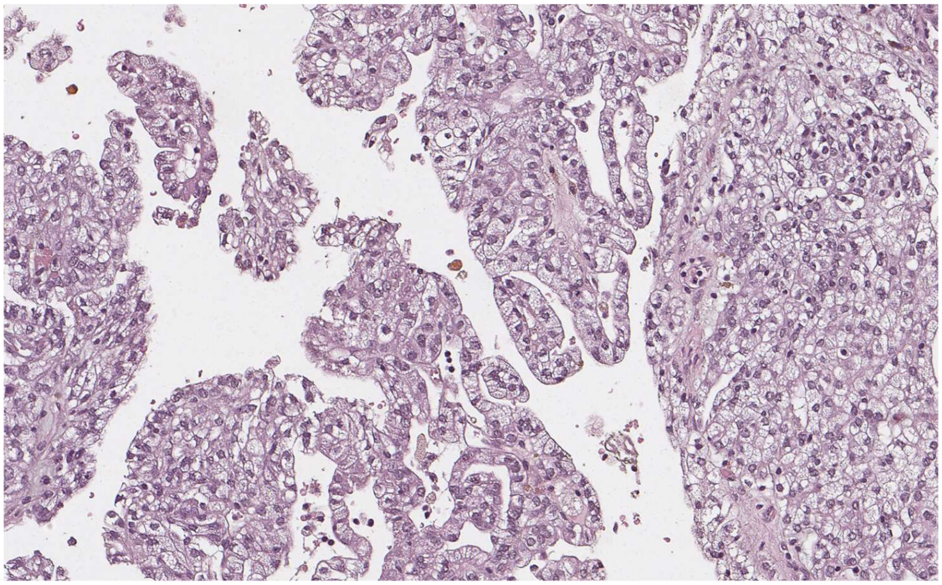

pRCC1_1501_01

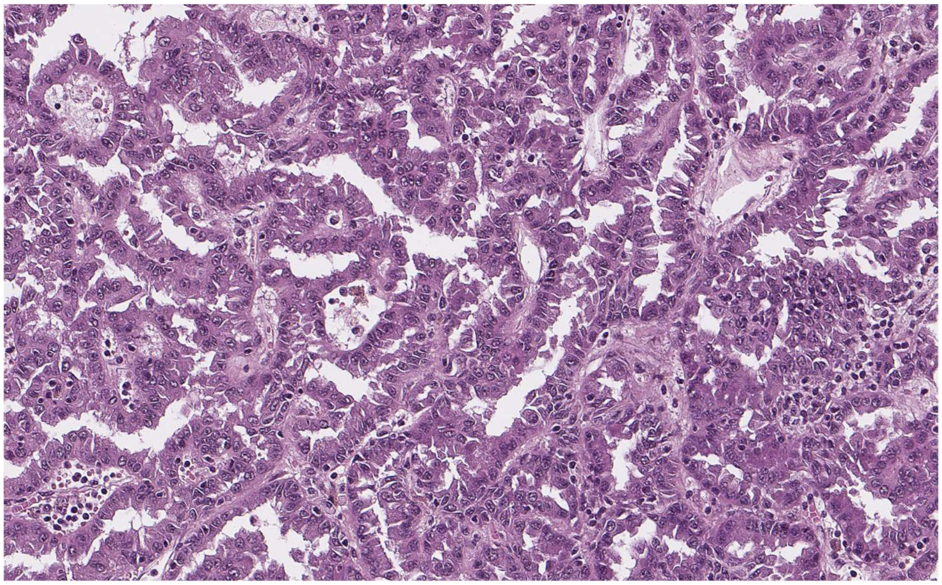

pRCC1_1546_01

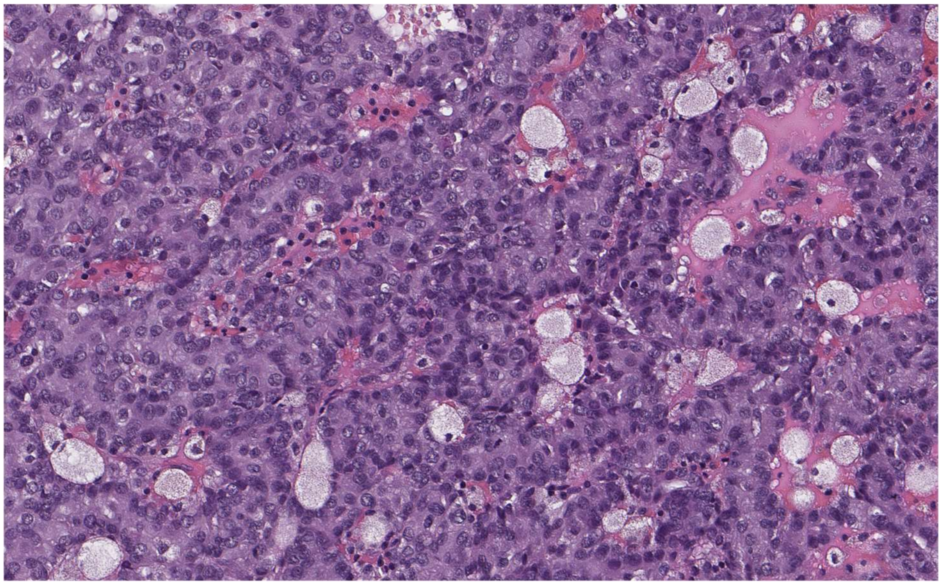

pRCC1_1550_02

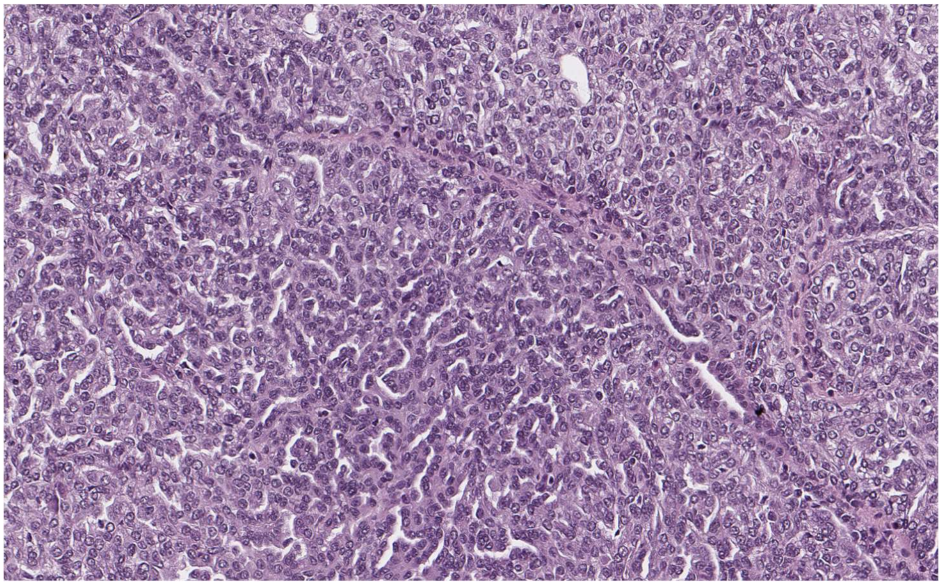

pRCC1_1654_01

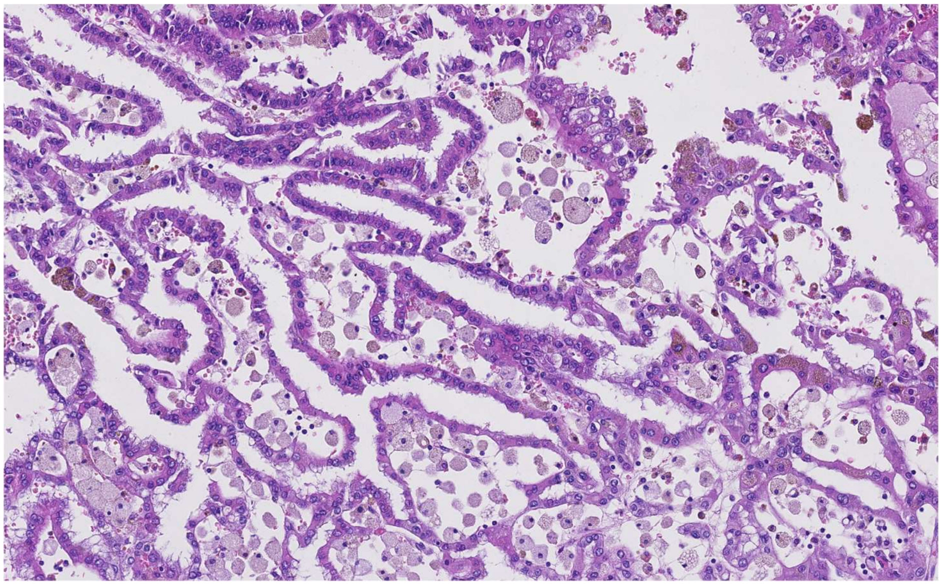

pRCC1_1656_01

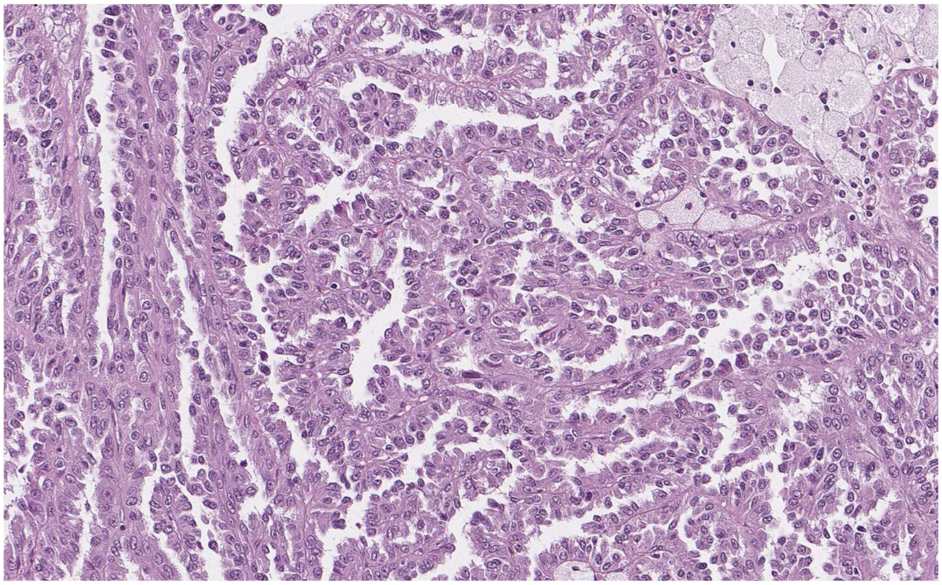

pRCC1_1662_01

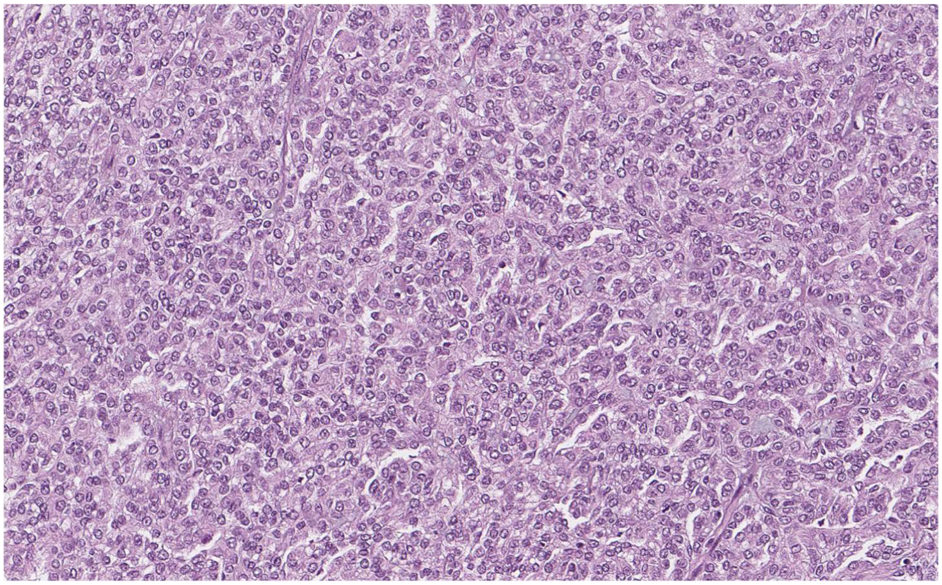

pRCC1_1671_08

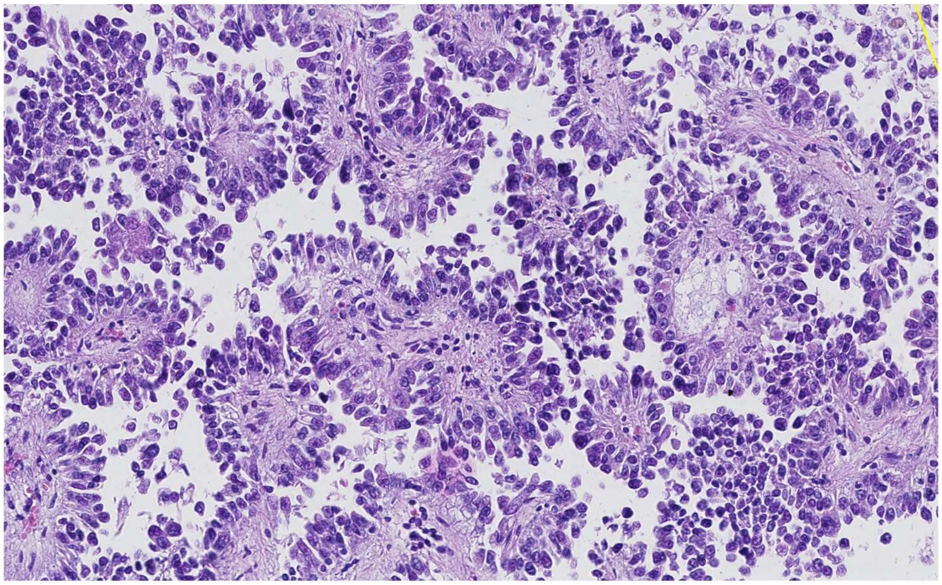

pRCC1_1689_06

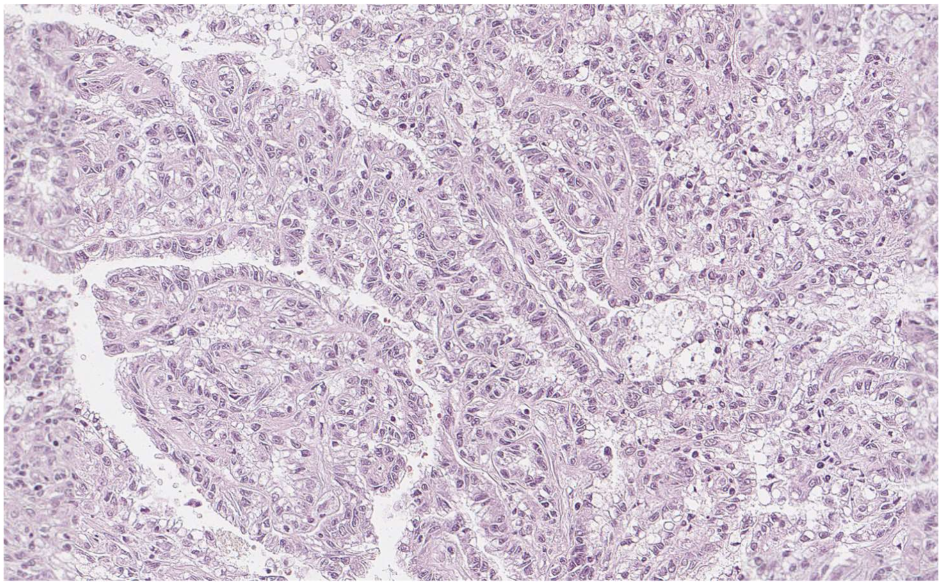

pRCC1_1699_01

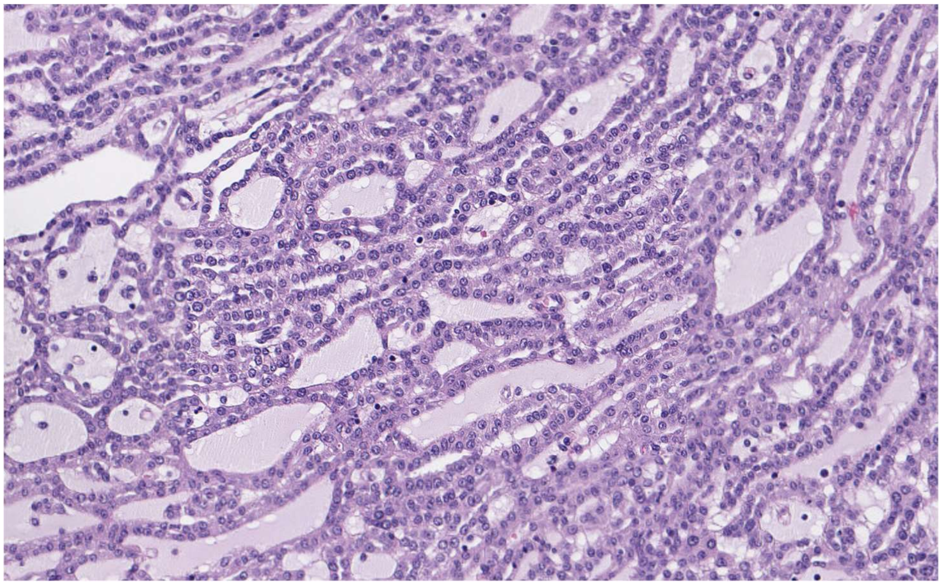

pRCC1_1708_01

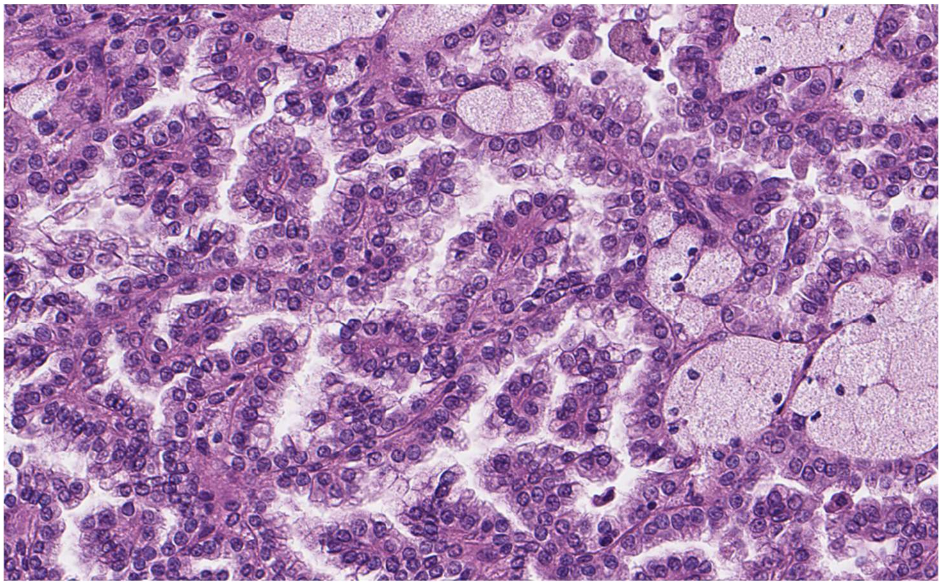

pRCC1_1735_01

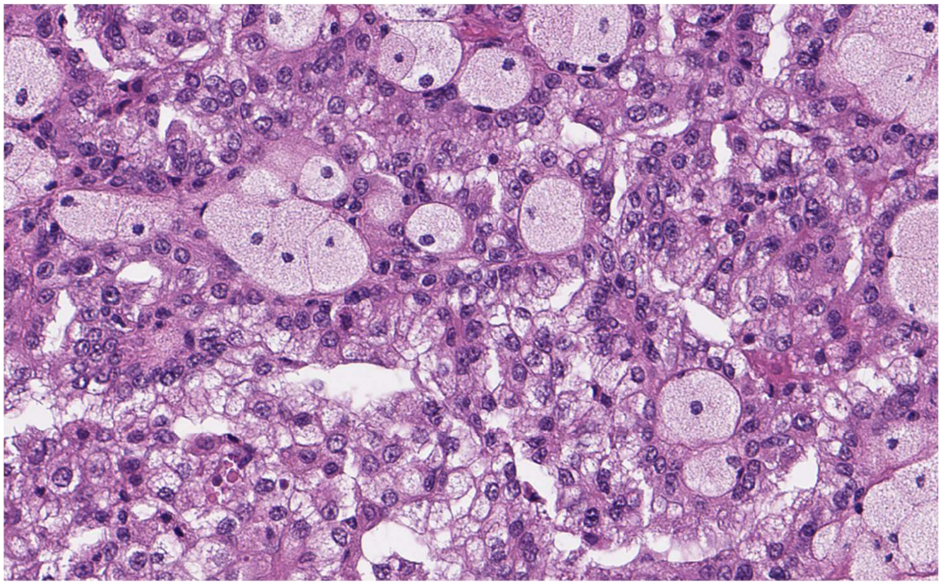

pRCC1_1844_01

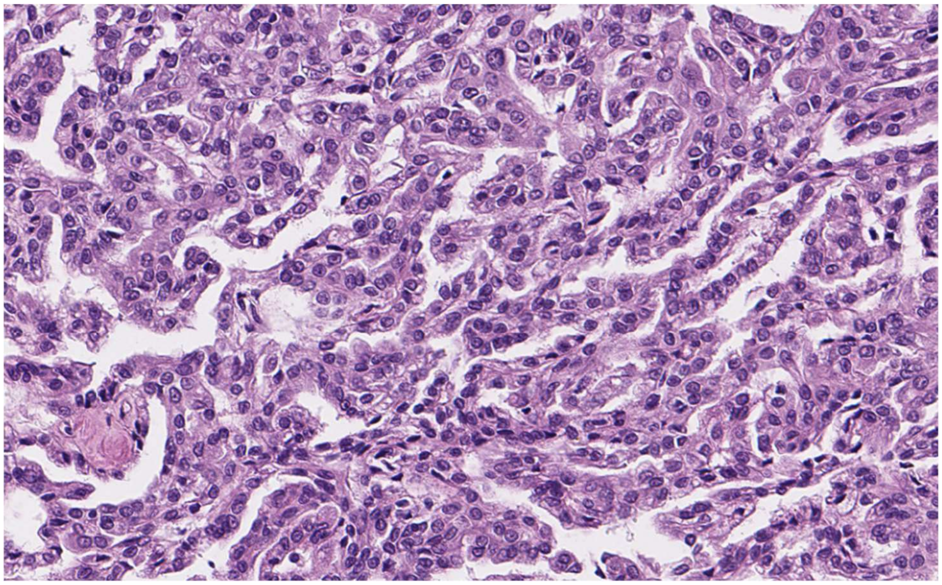

pRCC2_1298_01

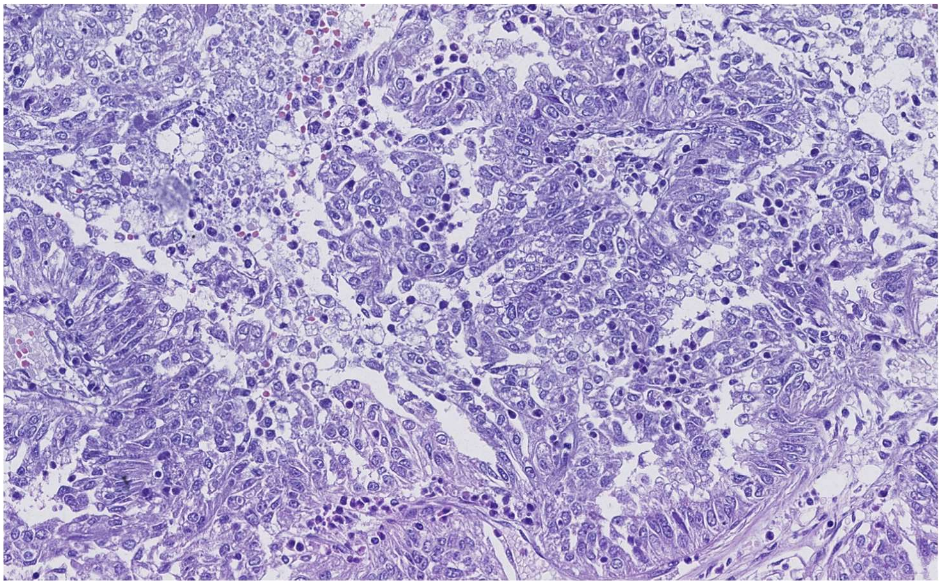

pRCC2_1396_01

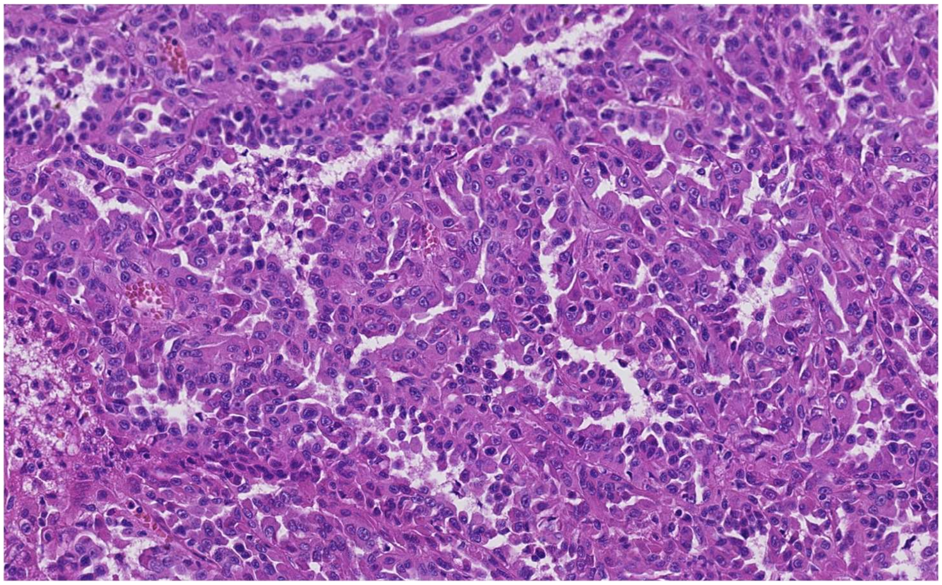

pRCC2_1410_02

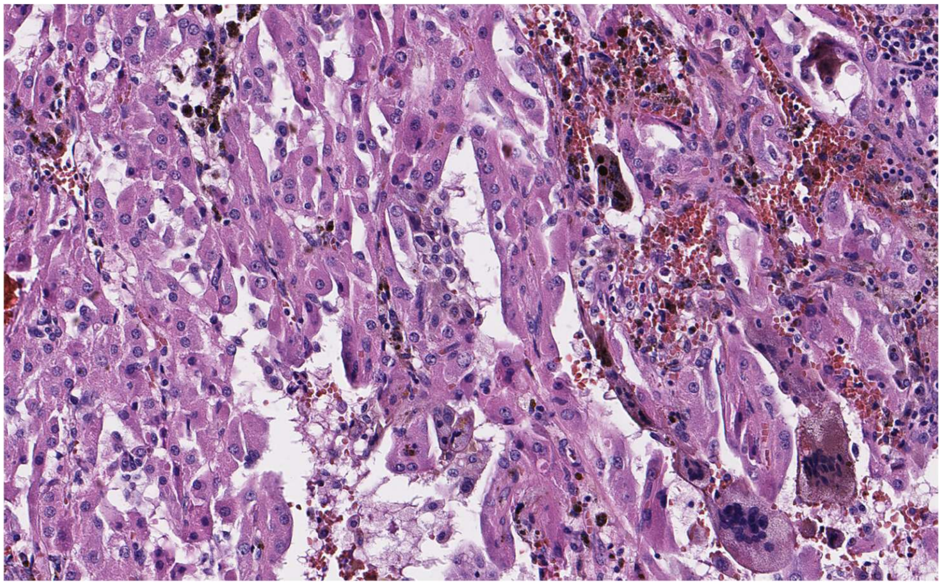

pRCC2_1429_03

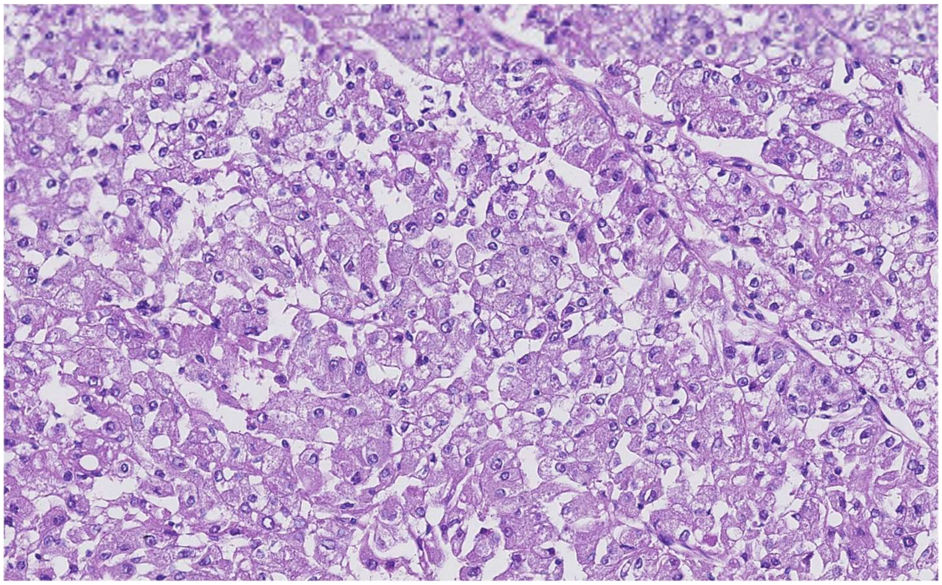

pRCC2_1479_03

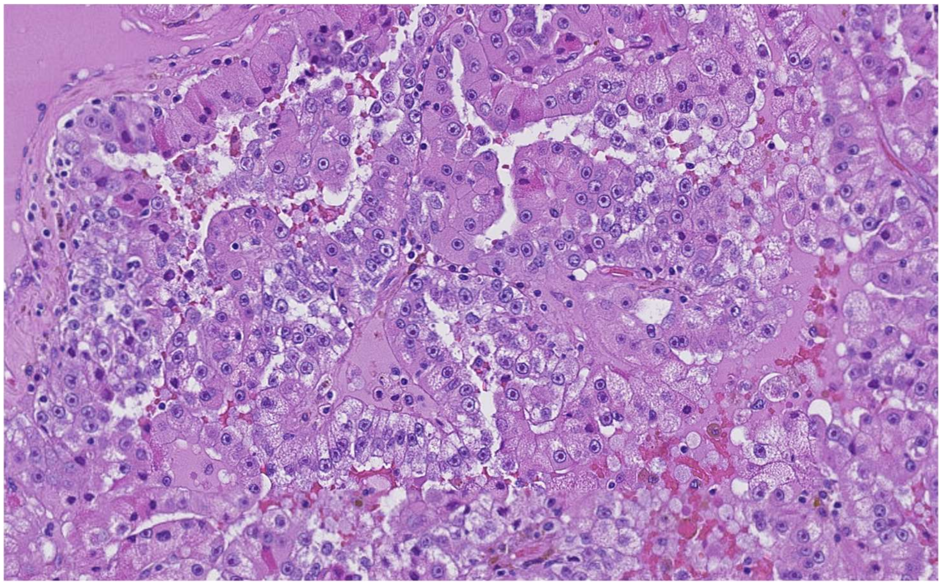

pRCC2_1494_04

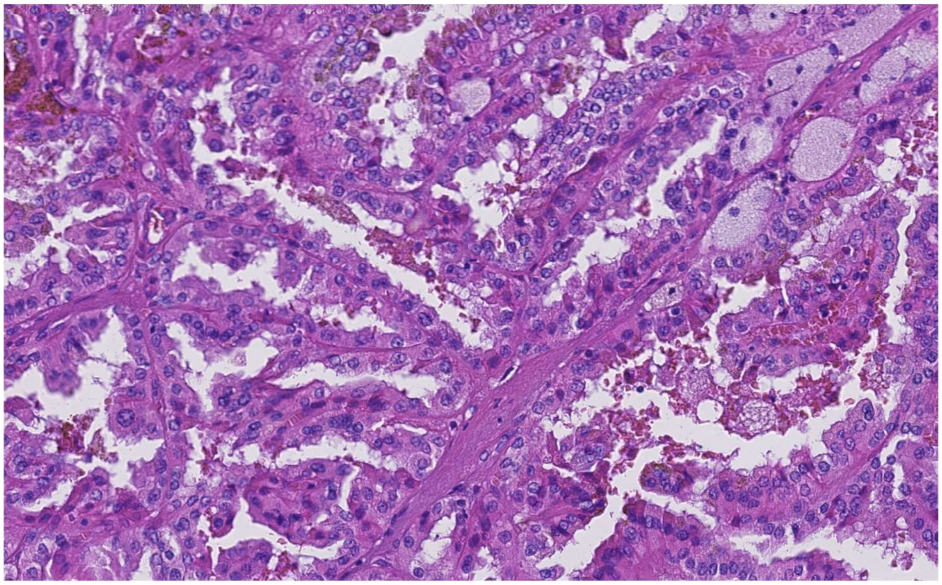

pRCC2_1552_03

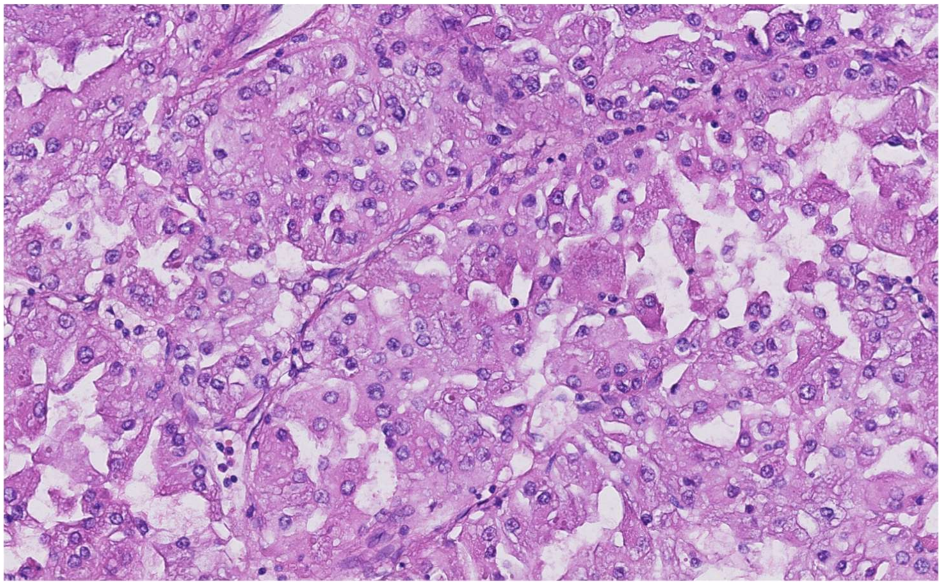

pRCC2_1568_04

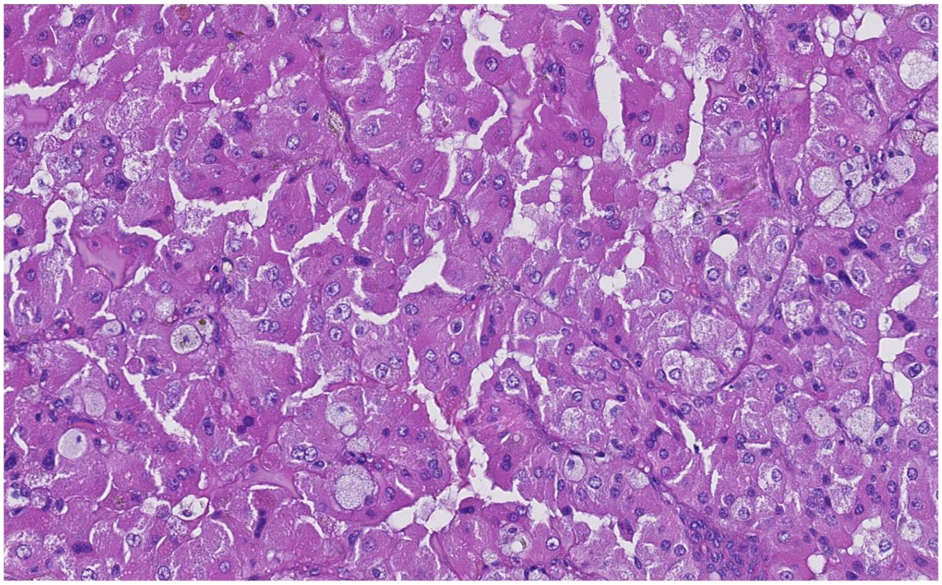

pRCC2_1782_08

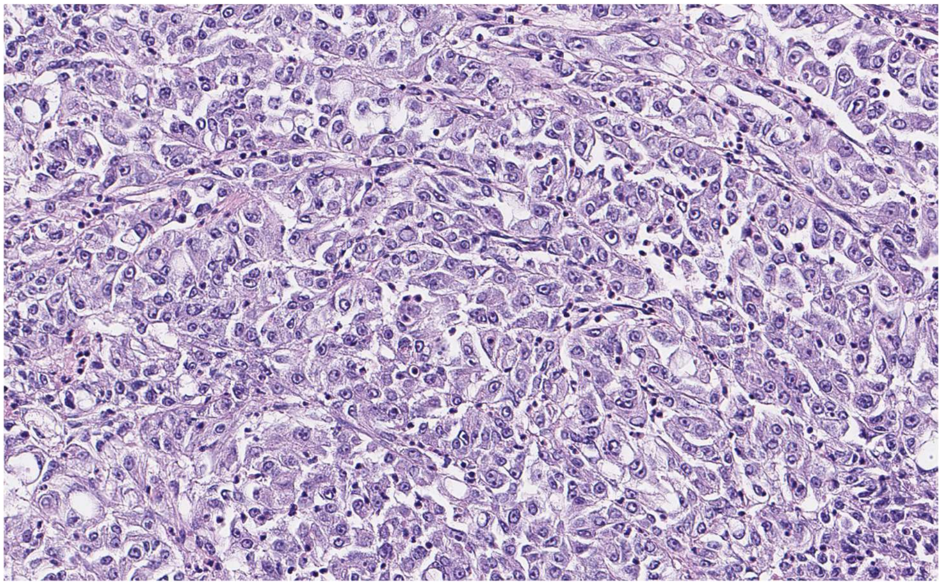

pRCC2_1799_02

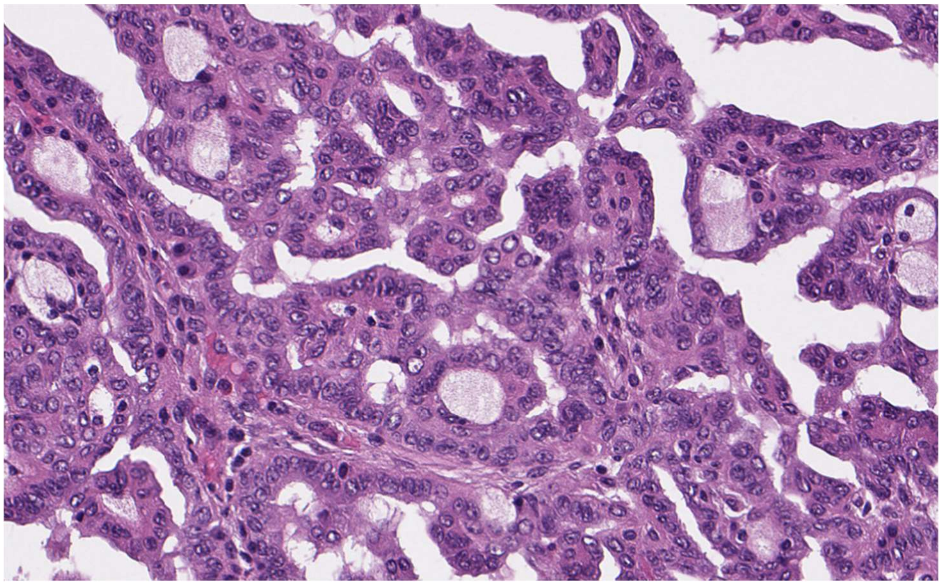

pRCC2_1824_13

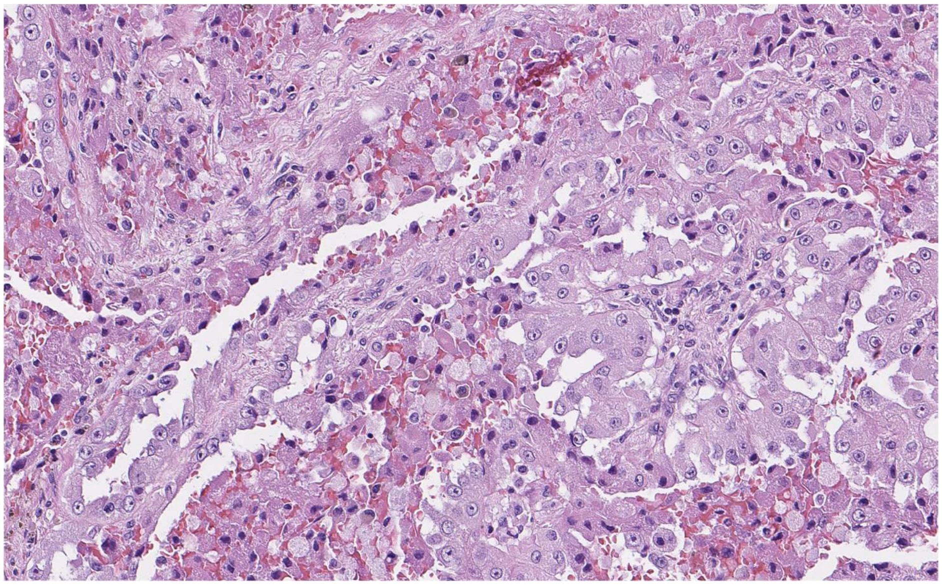

pRCC2_1851_04

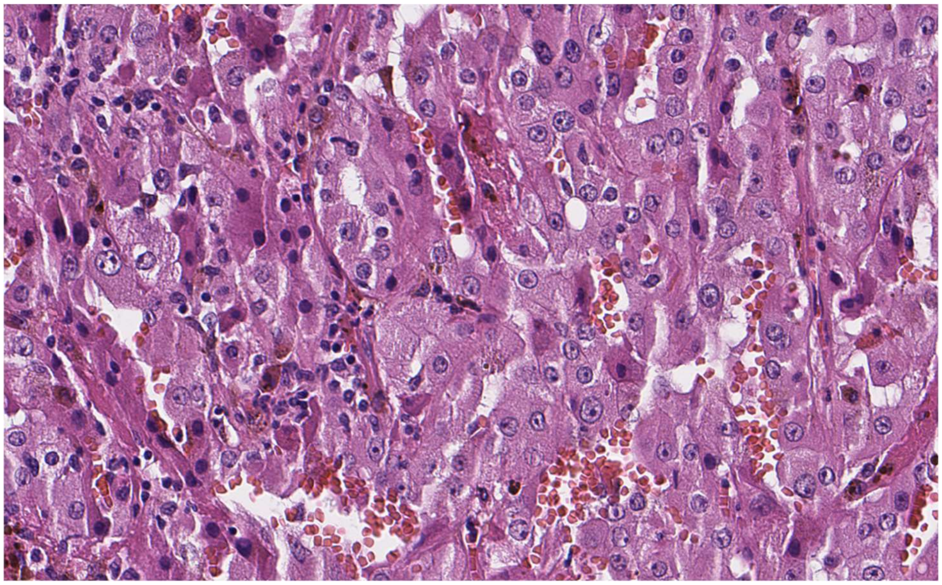

rSRC_1697_10

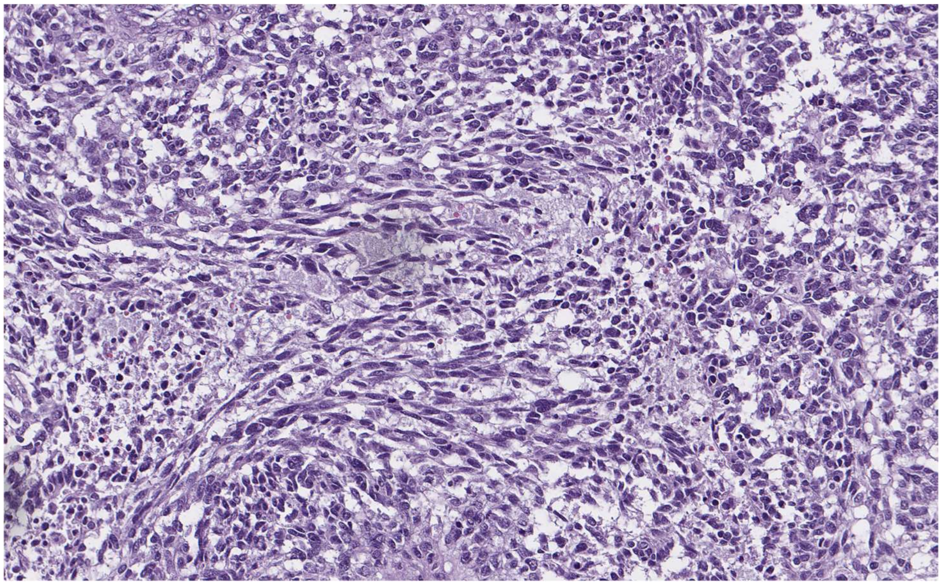

**Supplementary Figure 26: Venn Diagram of genomic data available across subjects.** The figure describes the intercept of whole genome sequencing, deep targeted sequencing, methylation array and genotyping array data across all 29 subjects.

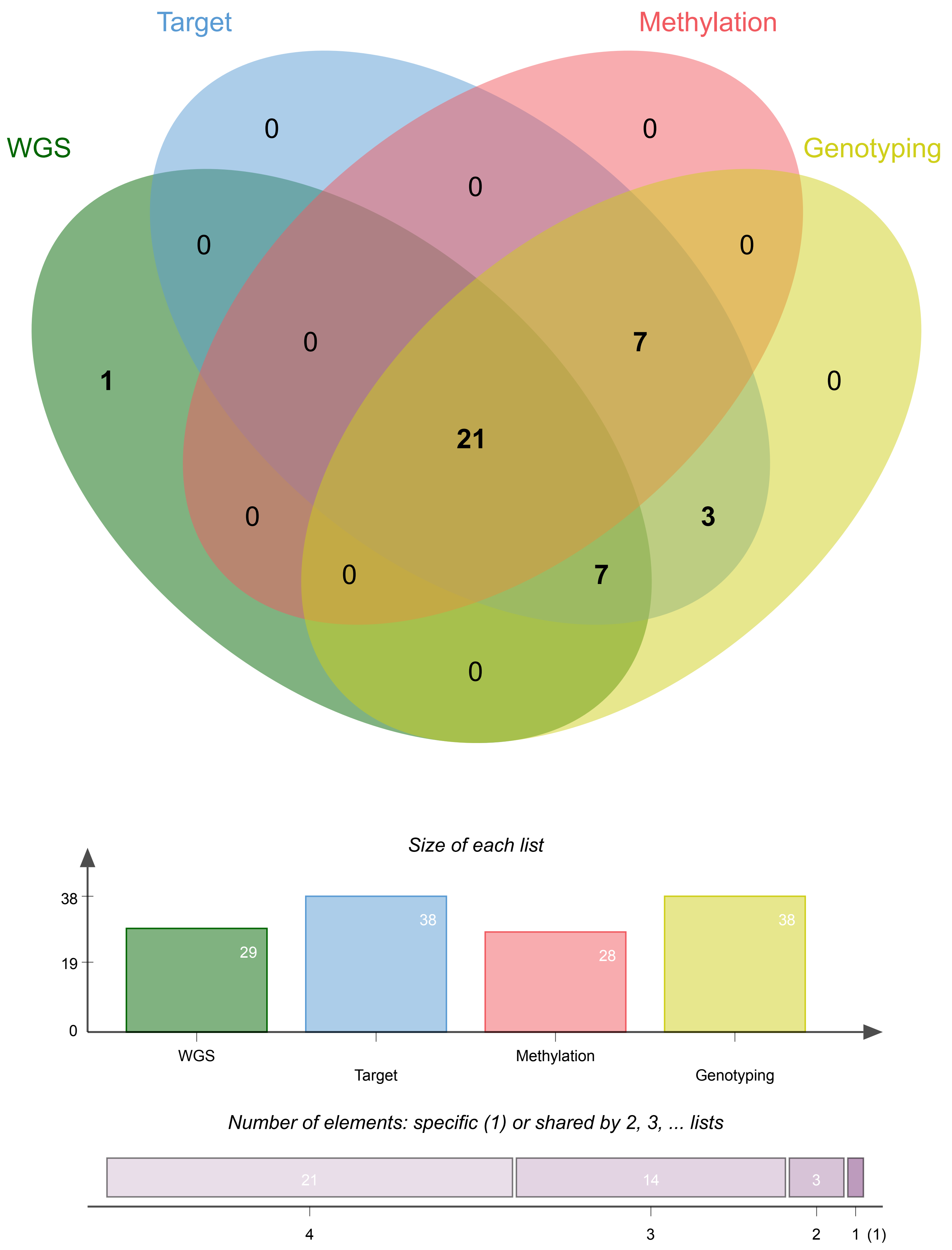
Target
Methylation
WGS
Genotyping
0
0
0
0
0
0
7
1
0
21
0
3
0
7
0
Size of each list
38
19
0
29
38
28
38
WGS
Target
Methylation
Genotyping
Number of elements: specific (1) or shared by 2, 3, ... lists
21
14
3
4
3
2
1 (1)

**Supplementary Figure 27: Topography of mutational signatures. (a)** The distributions of nucleosome density signals (y-axes) are shown in a 2 kb window centered on each mutation (position 0 on the x-axes), for each signature. The averaged signal was calculated as the total amount of signal observed at each point divided by total number of mutations contributing to that signal. **(b)** Distribution of the base substitution signatures across the cell cycle. Replication domains were identified by using conservatively defined transition zones in DNA replication time data. Data were separated into deciles, with each segment containing exactly 10% of the observed replication time signal. Normalized mutation density per decile is presented for early (left) to late (right) replication domains. (c) Transcription and replication strand-bias for mutational signatures and types of single-point somatic mutations. We performed two-tailed Fisher's exact test and the p-values were adjusted for multiple hypothesis testing using the Benjamini–Hochberg false discovery rate (FDR) procedure. Exact adjusted p-values which are significant (adjusted p-value<=0.05 represented by '*') for each plot are listed as follows. C>A: $2.61 \times 10^{-3}$, C>T: $1.12 \times 10^{-9}$, T>A: $6.40 \times 10^{-3}$, T>C: $1.21 \times 10^{-3}$, T>G: $5.13 \times 10^{-5}$ in plot of "Transcribed vs. Untranscribed All Mutations"; C>A: $3.28 \times 10^{-2}$, C>T: $8.42 \times 10^{-3}$, T>G: $3.10 \times 10^{-4}$ in plot of "Lagging vs. Leading All Mutations"; SBS-01: $8.42 \times 10^{-3}$, SBS-05: $3.25 \times 10^{-4}$, SBS-40: $1.37 \times 10^{-4}$ in plot of "Transcribed vs. Untranscribed All Signatures"; SBS-40: $1.38 \times 10^{-2}$ in plot of "Lagging vs. Leading All Signatures".

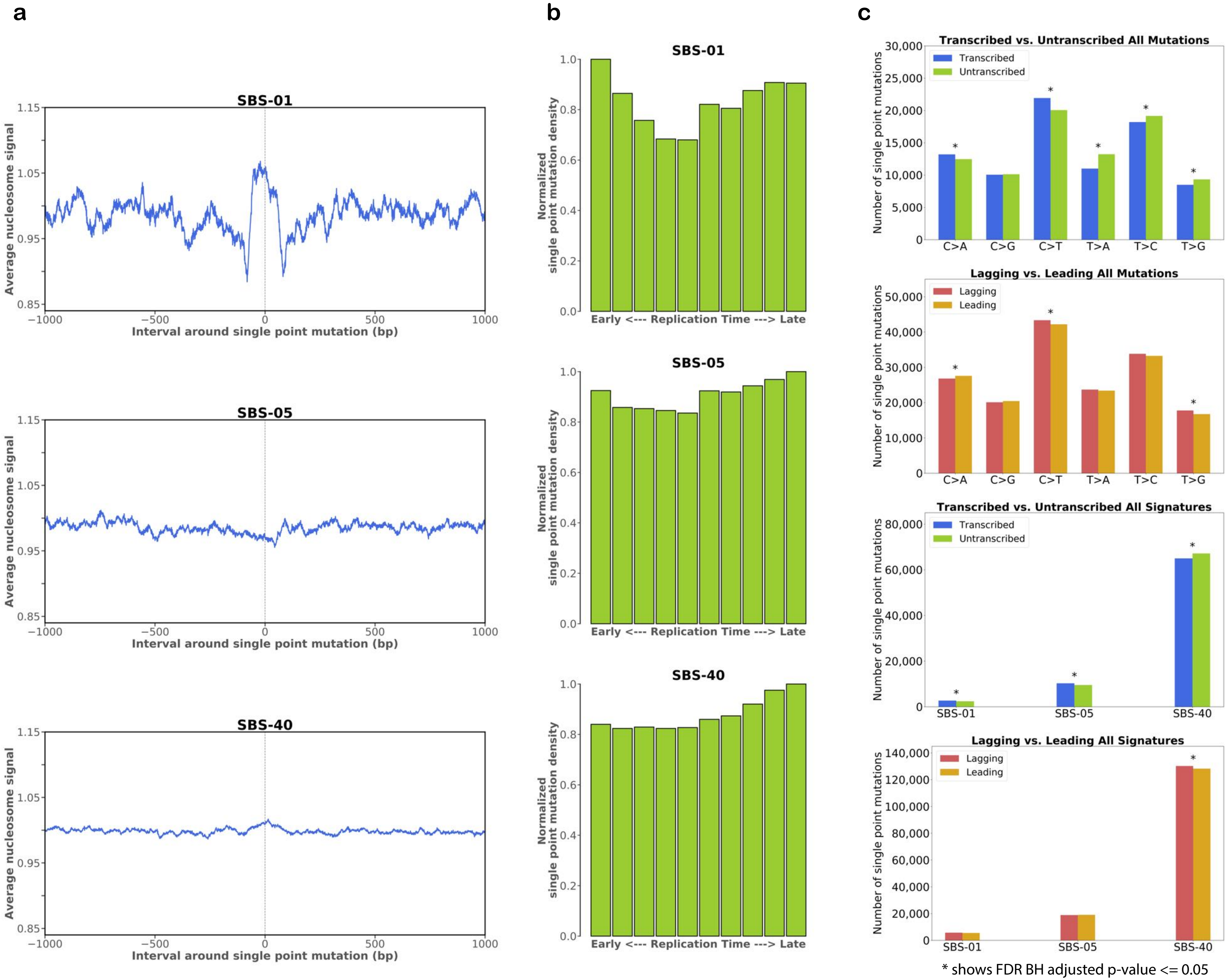
a
b
c
SBS-01
SBS-05
SBS-40
Average nucleosome signal
Interval around single point mutation (bp)
Normalized single point mutation density
Early <--- Replication Time ---> Late
Transcribed vs. Untranscribed All Mutations
Lagging vs. Leading All Mutations
Transcribed vs. Untranscribed All Signatures
Lagging vs. Leading All Signatures
Transcribed
Untranscribed
Lagging
Leading
Number of single point mutations
C>A
C>G
C>T
T>A
T>C
T>G
* shows FDR BH adjusted p-value <= 0.05

**Supplementary Figure 28: De novo extraction of indel mutational signatures.** Profile of the three *de novo* extracted signatures of small insertions and deletions (indels) is provided. Indels were classified as deletions or insertions and, when of a single base, as C or T and according to the length of the mononucleotide repeat tract in which they occurred. Longer indels were classified as occurring at repeats or with overlapping microhomology at deletion boundaries, and according to the size of indel, repeat, and microhomology.

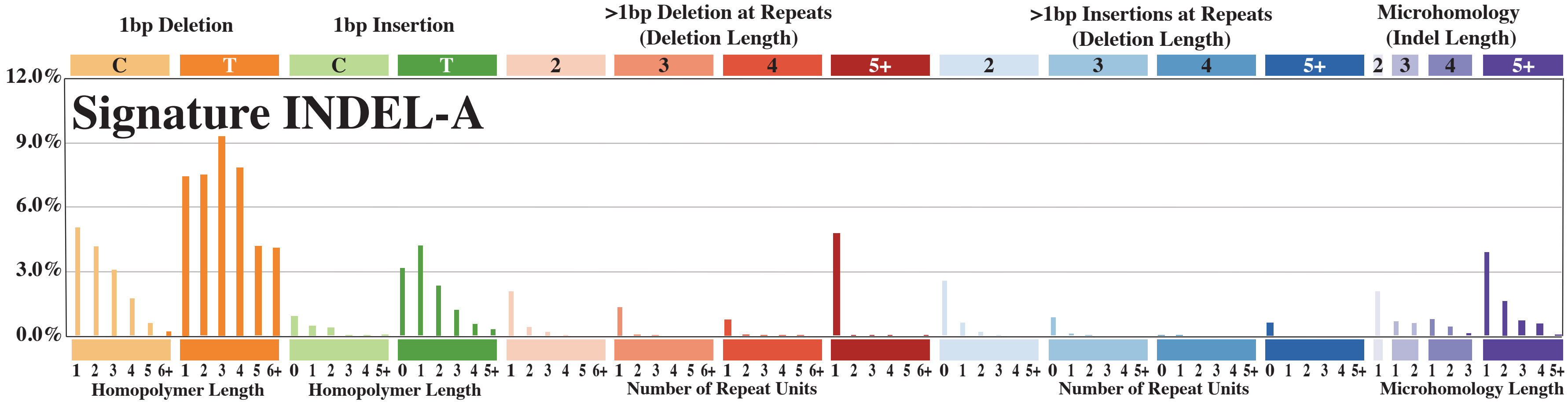
Signature INDEL-A
1bp Deletion
1bp Insertion
>1bp Deletion at Repeats (Deletion Length)
>1bp Insertions at Repeats (Deletion Length)
Microhomology (Indel Length)
C T C T 2 3 4 5+ 2 3 4 5+ 2 3 4 5+
12.0%
9.0%
6.0%
3.0%
0.0%
Homopolymer Length
Homopolymer Length
Number of Repeat Units
Number of Repeat Units
Microhomology Length

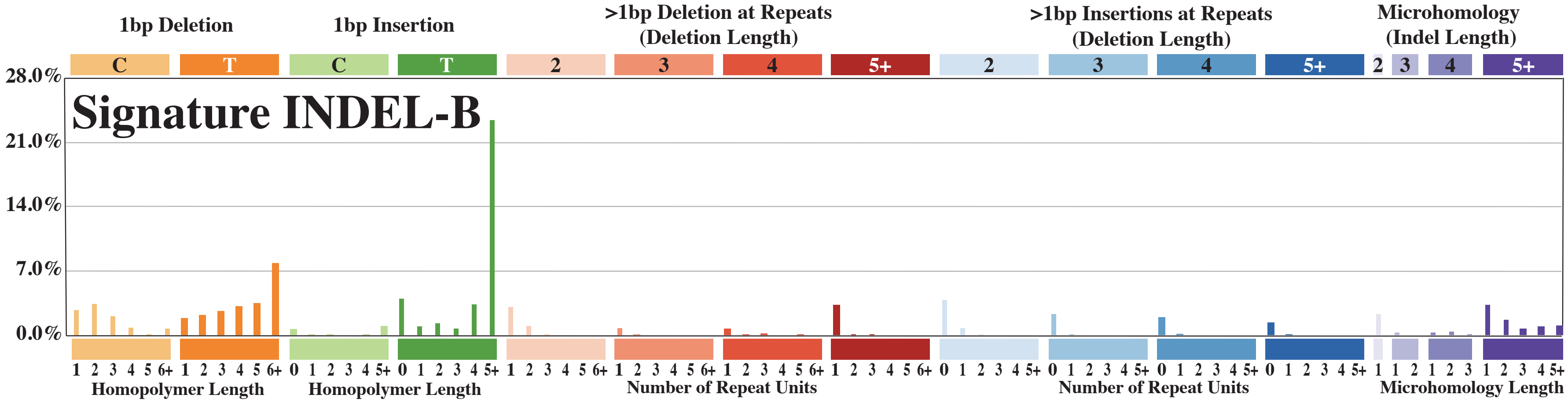
Signature INDEL-B
1bp Deletion
1bp Insertion
>1bp Deletion at Repeats (Deletion Length)
>1bp Insertions at Repeats (Deletion Length)
Microhomology (Indel Length)
C T C T 2 3 4 5+ 2 3 4 5+ 2 3 4 5+
28.0%
21.0%
14.0%
7.0%
0.0%
Homopolymer Length
Homopolymer Length
Number of Repeat Units
Number of Repeat Units
Microhomology Length

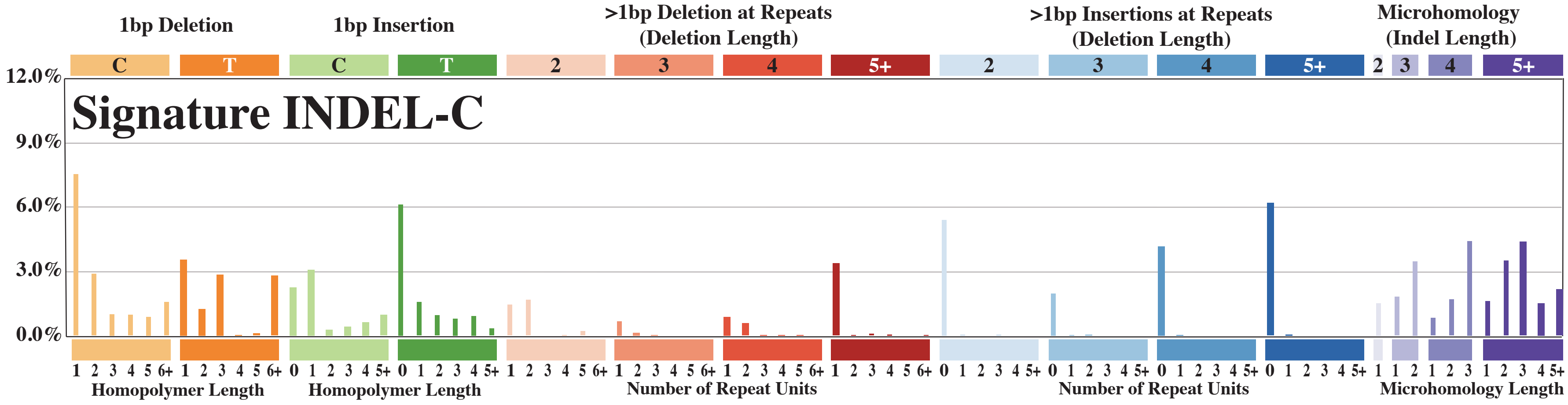
Signature INDEL-C
1bp Deletion
1bp Insertion
>1bp Deletion at Repeats (Deletion Length)
>1bp Insertions at Repeats (Deletion Length)
Microhomology (Indel Length)
C T C T 2 3 4 5+ 2 3 4 5+ 2 3 4 5+
12.0%
9.0%
6.0%
3.0%
0.0%
Homopolymer Length
Homopolymer Length
Number of Repeat Units
Number of Repeat Units
Microhomology Length
